# Compact type II-C Cas9 nucleases with expanded PAM access and high fidelity for therapeutic genome editing

**DOI:** 10.64898/2026.08.17.745178

**Authors:** Qiaochu Wang, Sivakrishna Rao Gundra, Rashid Aman, Ahmed Saleh, Ahmed M. Kazlak, Maazallah Masood, Norhan Hassan, Magdy M. Mahfouz

**Author notes:** Corresponding author: Magdy M. Mahfouz, Laboratory for Genome Engineering and Synthetic Biology, KAUST, Thuwal 23955-6900, Saudi Arabia. These authors contributed equally (Qiaochu Wang, Sivakrishna Rao Gundra).

## Abstract

Compact type II-C Cas9 nucleases are attractive for therapeutic genome editing because their small size enables packaging into adeno-associated viral (AAV) vectors, and their extended protospacer-adjacent motifs (PAMs) reduce off-target cleavage while expanding targeting scope. Yet characterized type II-C orthologs have edited mammalian cells far less efficiently than the canonical SpCas9. Here, we used embedding-based metagenomic mining of >4.7 × 10 proteins, combined with AlphaFold3 structure prediction and locus-context analysis, to identify three previously uncharacterized compact type II-C Cas9 orthologs, NsuCas9 (1,092 aa), PsuCas9 (1,084 aa), and GfoCas9 (1,074 aa), and benchmarked them *in vitro* and in human HEK293T cells. All three are robust RNA-guided nucleases with distinct PAM specificities (N CC, N NYAA, and N RHAA, respectively), divergent thermal profiles, and asymmetric sgRNA cross-compatibility. In human cells, PsuCas9 with an N ATAA PAM reaches 78.4% indels and matches or exceeds SpCas9 at multiple loci, representing the first natural compact type II-C ortholog reported to do so, while GfoCas9 and NsuCas9 add complementary coverage. All three show a strong deletion-biased repair signature and no detectable editing across 33 predicted off-target sites. These compact, high-fidelity nucleases expand the CRISPR targeting space for AAV-deliverable therapeutic editing.

## Introduction

Programmable genome editing with CRISPR-Cas9 has revolutionized biomedical research and is now advancing into clinical therapeutics.^1, 2^ Most of this progress has been built on a single nuclease, the type II-A Cas9 from *Streptococcus pyogenes* (SpCas9), which combines high activity in human cells with broad genomic targeting through its short 5’-NGG-3’ PAM.^3, 4^ Yet the very features that made SpCas9 the workhorse of the field also limit its therapeutic deployment. Its 1,368-amino-acid coding sequence approaches the packaging capacity of adeno-associated viral vectors when combined with guide RNA and regulatory elements,^5, 6^ while its short PAM increases the abundance of near-cognate off-targets across the human genome.^7, 8^ Compact Cas9 nucleases that are AAV-deliverable as a single construct, that recognize PAMs orthogonal to NGG, and that retain high specificity are therefore needed to translate the full promise of CRISPR into safe *in vivo* therapies.

Type II-C Cas9 enzymes are intrinsic candidates for these requirements.^9, 10^ They are typically 900–1,100 amino acids in length, several hundred residues smaller than SpCas9, and recognize extended PAMs of four to six bases that constrain the genomic search space and reduce promiscuous cleavage.^4, 11^ Despite these advantages, the subtype has remained markedly under-exploited. The type II-C orthologs validated in human cells to date, including Nme1Cas9,^12^ Nme2Cas9,^10^ CjCas9,^9^ PpCas9,^13^ CdCas9,^14^ GeoCas9,^15^ BlatCas9,^16^ and CoCas9,^17^ typically recognize complex PAMs (for example, 5’-N GATT-3’ for Nme1Cas9, 5’-N RYAC-3’ for CjCas9, and 5’-N CRAA-3’ for GeoCas9) and edit mammalian cells at frequencies substantially below SpCas9.^4, 18^ The unmet need is a compact type II-C nuclease that combines SpCas9-level on-target activity with the subtype’s intrinsic fidelity and PAM diversity. Two complementary strategies have been pursued to close this gap. The first is protein engineering of established type II-C orthologs: rational engineering or directed evolution of CjCas9 has produced variants such as enCjCas9 (5’-N VRYA-3’),^19^ evoCjCas9 (5’-N AH-3’ and 5’-N HA-3’)^20^ and UltraCjCas9 (5’-N RYAY-3’)^21^ with enhanced activity and relaxed PAMs, and domain swaps between closely related orthologs have generated chimeras such as Nsp2–SmuCas9 with simplified 5’-N C-3’ PAMs.^22^ These engineered variants expand the targeting range, but the gain in PAM flexibility frequently comes at the cost of specificity.^20^ The complementary strategy is discovery of new natural orthologs, which preserves wild-type fidelity but has been pursued largely on a one-at-a-time basis.^22–25^ Recent biochemical surveys have begun to enable systematic discovery,^11^ yet the breadth of microbial sequence space accessible through modern embedding-based homology search, including the >4 × 10 protein sequences now in MGnify, remains largely untapped for compact, high-fidelity, broadly targeting Cas9 nucleases.

Here we apply this approach. Embedding-based Dense Homology Retrieval (DHR) of more than 4.7 × 10 proteins in the MGnify metagenomic repository, integrated with AlphaFold3 structure prediction,^26^ CRISPR-locus context analysis, and protein-language-model PAM prediction^27^, prioritized three previously uncharacterized compact type II-C Cas9 orthologs: NsuCas9 (1,092 aa), PsuCas9 (1,084 aa), and GfoCas9 (1,074 aa), named for their closest BLASTp hits (*Neisseria subflava*, *Phascolarctobacterium succinatutens*, and *Gemmiger formicilis*, respectively). All three are robust RNA-guided dsDNA nucleases *in vitro*, with distinct PAM specificities (5’-N CC-3’ for NsuCas9, 5’-N NYAA-3’ for PsuCas9, and 5’-N RHAA-3’ for GfoCas9), divergent thermal profiles, and asymmetric sgRNA cross-compatibility. In human HEK293T cells, PsuCas9 paired with a 5’-N ATAA-3’ PAM matches or exceeds SpCas9 at multiple endogenous loci and reaches indel frequencies up to 78.4%, to our knowledge representing the first natural compact type II-C Cas9 ortholog reported to outperform SpCas9 in mammalian editing. None of the three nucleases produced detectable editing across 33 computationally predicted off-target sites, including sites bearing only two spacer mismatches. Together, these enzymes provide a complementary set of compact, high-fidelity editors that substantially expand the sequence space accessible to CRISPR-Cas9 and qualify as candidate scaffolds for AAV-deliverable therapeutic genome editing and for adaptation into base-editor, prime-editor, and epigenome-editor derivatives.

## Results

### Metagenomic mining identifies three uncharacterized compact type II-C Cas9 orthologs

To identify novel compact Cas9 nucleases, we performed large-scale *in silico* mining of the MGnify metagenomic protein repository using a curated set of 49 representative Cas9 sequences spanning the diversity of CRISPR-Cas9 subtypes as queries.^28^ Searches were carried out with an embedding-based Dense Homology Retrieval (DHR) framework, which detects remote homologs that retain conserved structural features despite extensive sequence divergence. From a database of more than 4.7 × 10 full-length proteins we recovered the top 10 hits per query and enriched for compact variants by removing proteins longer than 1,100 amino acids, leaving 1.25 × 10 sequences after deduplication. Sequence-and structure-based annotation excluded candidates lacking a complete RuvC catalytic triad or HNH nuclease domain, retaining 4.5 × 10 high-confidence Cas9 homologs. CRISPR arrays and tracrRNA loci were detected at 35% and 28% of these sites, respectively. AlphaFold3 structure prediction,^26^ structural alignment to reference orthologs (SaCas9 [PDB: 5AXW],^29^ Nme1Cas9 [PDB: 6JDV],^30^ GeoCas9 [PDB: 8UZA]^15^), and Protein2PAM-based prediction of PAM diversity^27^ prioritized three high-confidence novel candidates for experimental characterization: NsuCas9 (1,092 aa), PsuCas9 (1,084 aa), and GfoCas9 (1,074 aa) (Data S1 and Table S1). The three orthologs adopt conserved three-dimensional architectures (Figure S1), display direct-repeat and tracrRNA features consistent with closely related orthologs (Figure S2), and are predicted to recognize distinct PAMs (Figure S3A).

### Phylogeny, locus organization, and in vitro activity

To establish that the three candidates are bona fide type II-C Cas9 enzymes, we first characterized their evolutionary placement and locus organization. Maximum-likelihood phylogenetic reconstruction with 122 reference Cas9 proteins (Data S1) confirmed that all three orthologs belong to the type II-C clade containing NmeCas9, AtCas9, and CjCas9, but occupy distinct evolutionary lineages: NsuCas9 groups with the *Neisseria*-derived nucleases (Nme1Cas9, Nme2Cas9), whereas PsuCas9 and GfoCas9 cluster together adjacent to AtCas9 and more distantly from the NmeCas9 clade (Figure 1A). Locus analysis with CRISPRCasTyper^31^ showed that each Cas9 gene is embedded in a canonical type II-C architecture comprising *cas1*, *cas2*, a CRISPR array, and a tracrRNA region (Figure 1B). Domain annotation by sequence and structural alignment confirmed that all three proteins contain the expected REC and nuclease lobes, including RuvC and HNH catalytic domains together with WED and PAM-interacting domains (Figure 1C).

**Figure 1.**
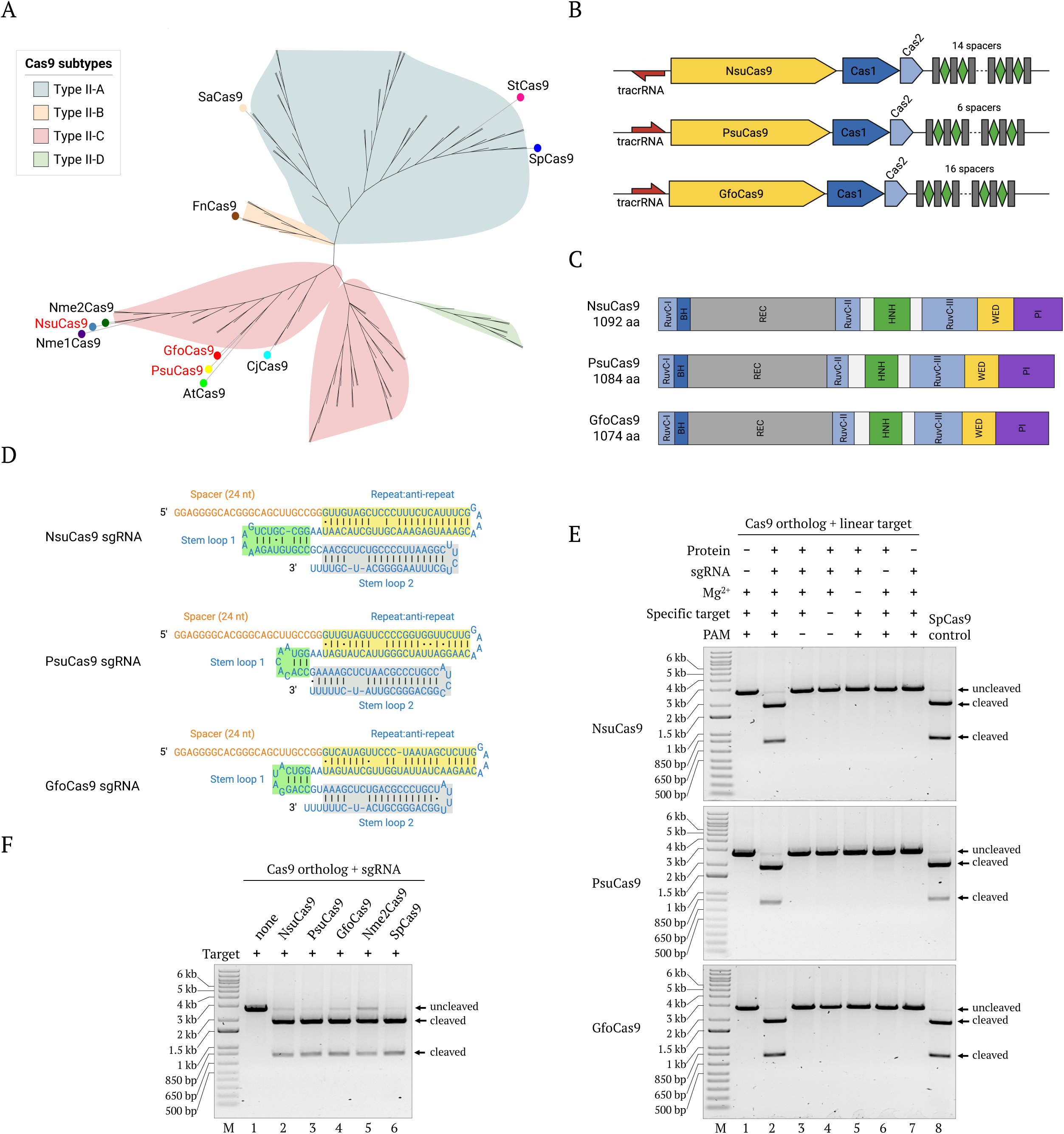
Identification and initial characterization of three compact type II-C Cas9 orthologs. (A) Maximum-likelihood phylogeny of NsuCas9, PsuCas9 and GfoCas9 with representative Cas9 proteins from type II-A, II-B, II-C and II-D subtypes (color-coded: II-A, blue; II-B, orange; II-C, pink; II-D, green); the three orthologs identified in this study are highlighted in red. (B) Organization of the CRISPR-Cas loci of NsuCas9, PsuCas9 and GfoCas9 predicted by CRISPRCasTyper. Yellow arrows, Cas9 genes; blue arrows, accessory genes including the *cas1*– *cas2* adaptation module; black rectangles, CRISPR repeats; green diamonds, spacers; red arrows, predicted tracrRNA transcription direction. (C) Domain architecture of NsuCas9, PsuCas9 and GfoCas9; boundaries inferred from multiple sequence alignment and refined manually. (D) Predicted sgRNA secondary structures: spacer (orange), repeat:anti-repeat duplex (yellow), stem-loop 1 (green), stem-loop 2 (gray). (E) *In vitro* cleavage activity on XmnI-linearized plasmid substrates; SpCas9 included as control. “Specific target,” plasmid carrying a sequence complementary to the spacer; “PAM,” substrate carrying a compatible PAM. (F) Side-by-side comparison of cleavage activity for NsuCas9, PsuCas9, GfoCas9, Nme2Cas9 and SpCas9 under identical reaction conditions. See also Figures S1–S5 and Tables S1–S3.

Having confirmed type II-C identity, we next tested whether the three proteins function as RNA-guided nucleases. sgRNAs were designed by fusing the predicted crRNA repeat to the cognate tracrRNA via a 5’-GAAA-3’ tetraloop (Figure 1D and Figure S3B and Table S2). The resulting scaffolds, each carrying a 24-nt repeat:anti-repeat duplex followed by two stem-loops, share the canonical type II-C architecture.^11, 32, 33^ Each Cas9 was expressed in *Escherichia coli* and purified to homogeneity (Figure S4). Reconstituted ribonucleoprotein (RNP) complexes cleaved linearized plasmid substrates only when Cas9, sgRNA, target DNA, a compatible PAM, and Mg² were all present, confirming RNA-guided, target-specific nuclease activity (Figure 1E). After optimization of buffer composition, sgRNA-to-Cas9 stoichiometry, and RNP concentration (Figure S5), the three new nucleases displayed robust *in vitro* cleavage comparable to SpCas9 and exceeding Nme2Cas9 under identical conditions (Figure 1F).

### PAM specificities define orthogonal targeting ranges

To map the targeting space accessible to each nuclease, we determined PAM requirements with an *in vitro* PAM screen using a randomized N library upstream of a fixed protospacer.^34^ RNPs assembled with a 24-nt-spacer sgRNA were incubated with the library, and cleaved fragments were enriched, sequenced, and analyzed by a custom pipeline. Position-specific enrichment, visualized using Logomaker,^35^ revealed distinct PAM preferences for the three nucleases (Figure 2A). As with other characterized type II-C orthologs,^9, 12^ specificity was concentrated at PAM positions 5–8. NsuCas9 strongly preferred 5’-N CC-3’, mirroring its closest characterized ortholog Nme2Cas9.^10^ PsuCas9 and GfoCas9 instead recognized AT-rich PAMs of the form 5’-N VYAA-3’ (V = A/G/C; Y = C/T) and 5’-N RHAA-3’ (R = A/G; H = A/C/T), respectively, that are orthogonal to the PAMs used by current type II-A and type II-C editors and broaden the addressable AT-rich genomic space.

**Figure 2.**
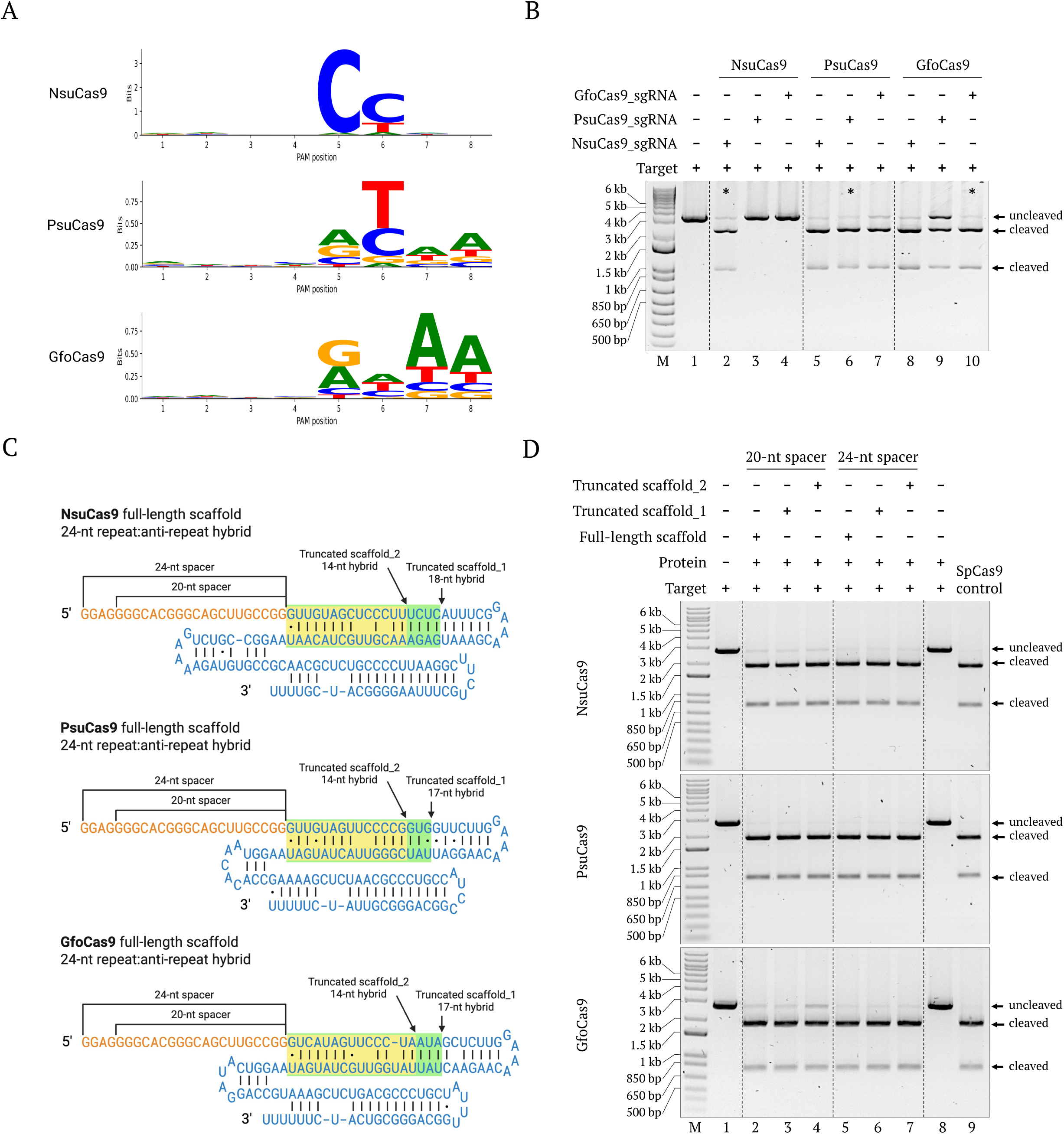
In vitro PAM characterization, sgRNA cross-compatibility, and scaffold optimization of type II-C Cas9 orthologs. (A) Sequence-logo representation of *in vitro* PAM preferences for NsuCas9, PsuCas9, and GfoCas9, derived from enriched cleavage reads using a DNA library containing an 8-nt randomized PAM (N) adjacent to a fixed protospacer. (B) *In vitro* cleavage assays of NsuCas9, PsuCas9, and GfoCas9 with each of the three sgRNA scaffolds. Lanes 2, 6, and 10 (asterisks) correspond to each Cas9 paired with its cognate sgRNA scaffold. (C) Schematic of full-length and truncated sgRNA designs. The full-length scaffold (blue) contains a 24-nt repeat:anti-repeat duplex, whereas truncated scaffold_1 (green box) and truncated scaffold_2 (yellow box) contain 17–18 nt and 14 nt duplexes, respectively. All constructs include a 5’-GAAA-3’ tetraloop linker and either a 20-nt or 24-nt spacer. (D) *In vitro* cleavage assays using full-length and truncated sgRNA scaffolds with 20-nt or 24-nt spacers under identical conditions. See also Figure S6 and Tables S4–S8.

Cleavage assays with linearized plasmid substrates carrying defined PAM variants confirmed the sequencing-based preferences (Figure S6). GfoCas9 was strictly limited to 5’-N RHAA-3’ PAMs, whereas PsuCas9 exhibited substantially broader PAM compatibility, supporting efficient cleavage across multiple variants, including targets with 5’-N NYAA-3’, 5’-N NAYA-3’, and 5’-N NYAY-3’ PAMs (with the exception of 5’-N CCAT-3’), as well as 5’-N GTTT-3’. Cleavage was attenuated for 5’-N NAAA-3’ PAMs, indicating that excessive adenine content reduces rather than enhances PsuCas9 activity. From these data we selected several uncommon but robust PAMs, notably 5’-N AYAA-3’ and 5’-N GAAA-3’, for evaluation in mammalian cells.

### Asymmetric sgRNA cross-compatibility and tolerance to scaffold truncation

Closely related Cas9 orthologs can sometimes interchange sgRNA scaffolds.^36^ We tested this systematically by pairing each of the three new nucleases with all three sgRNA scaffolds. Compatibility was strongly asymmetric (Figure 2B), as NsuCas9 was strictly dependent on its cognate sgRNA, whereas PsuCas9 and GfoCas9 retained activity with all three scaffolds, with PsuCas9 the most tolerant overall. Notably, the NsuCas9 sgRNA supported efficient cleavage by all three enzymes, including GfoCas9, which was paradoxically less active with the more closely related PsuCas9 sgRNA scaffold. The broad compatibility of the NsuCas9 scaffold may be attributed to its extended stem-loop 1 relative to the other two scaffolds; previous work has shown that incorporating the Nme1Cas9 stem-loop 1 into engineered sgRNAs improves cleavage efficiency in heterologous Cas9 systems,^37^ suggesting that this stem-loop strengthens RNP assembly across orthologs.

To test whether the scaffold could be further compacted, we shortened the repeat:anti-repeat duplex from 24 nt (full-length scaffold) to 17–18 nt (truncated scaffold_1) or 14 nt (truncated scaffold_2), and combined each scaffold with a 20-nt or 24-nt spacer (Figure 2C). Cleavage was minimally perturbed by duplex truncation (Figure 2D), but 24-nt spacers consistently outperformed 20-nt spacers, in agreement with prior observations for other type II-C orthologs.^9, 10^ All subsequent *in vitro* experiments therefore used 24-nt spacers.

### Distinct temperature ranges and blunt cleavage three nucleotides upstream of the PAM

The three nucleases displayed strikingly different thermal profiles *in vitro* (Figure S7). NsuCas9 was the most thermostable, retaining activity up to 50 °C and showing detectable cleavage at 55 °C. PsuCas9 was unusual in retaining robust activity at 20 °C, a temperature at which most characterized Cas9 enzymes are essentially inactive.^11^ GfoCas9 was the most narrowly tuned, with optimal cleavage between 30 °C and 40 °C. These differences likely reflect the distinct environmental niches of the source microbes and provide a practical handle for selecting enzymes for low-or high-temperature editing applications.

To map the cleavage geometry of each nuclease, the RNP of each enzyme was incubated with two plasmid substrates carrying different high-activity PAMs. The cleaved fragments were gel-purified and Sanger-sequenced. Mapping confirmed that all three nucleases generate blunt-ended double-strand breaks three nucleotides upstream of the PAM (Figure S8), consistent with many characterized type II-C Cas9 enzymes. The shared blunt-cut geometry positions all three orthologs for direct adaptation into base-editing, prime-editing, and epigenome-modulating fusion architectures, where the relative position of nickase activity is essential.

### Genome editing in human cells: PsuCas9 outperforms SpCas9 at multiple loci

We next assayed editing of endogenous human loci in HEK293T cells, with SpCas9-NGG and Nme2Cas9-N CC as benchmarks. We initially tested six target sites across four genes (*CLTA* t1–t2; *HBB* t1–t2; *AIFM* t1; *Casp-3* t1) under ten conditions: untransfected control, SpCas9-NGG, Nme2Cas9-N CC, NsuCas9-N CC, and PsuCas9 or GfoCas9 each paired with three candidate PAMs (5’-N ATAA-3’, 5’-N ACAA-3’, and 5’-N GAAA-3’). Seventy-two hours after transfection, T7EI assays revealed strong cleavage by PsuCas9-N ATAA, PsuCas9-N ACAA, GfoCas9-N GAAA, and SpCas9 (Figure 3A); NsuCas9 and Nme2Cas9 produced weaker but detectable products at some loci.

**Figure 3.**
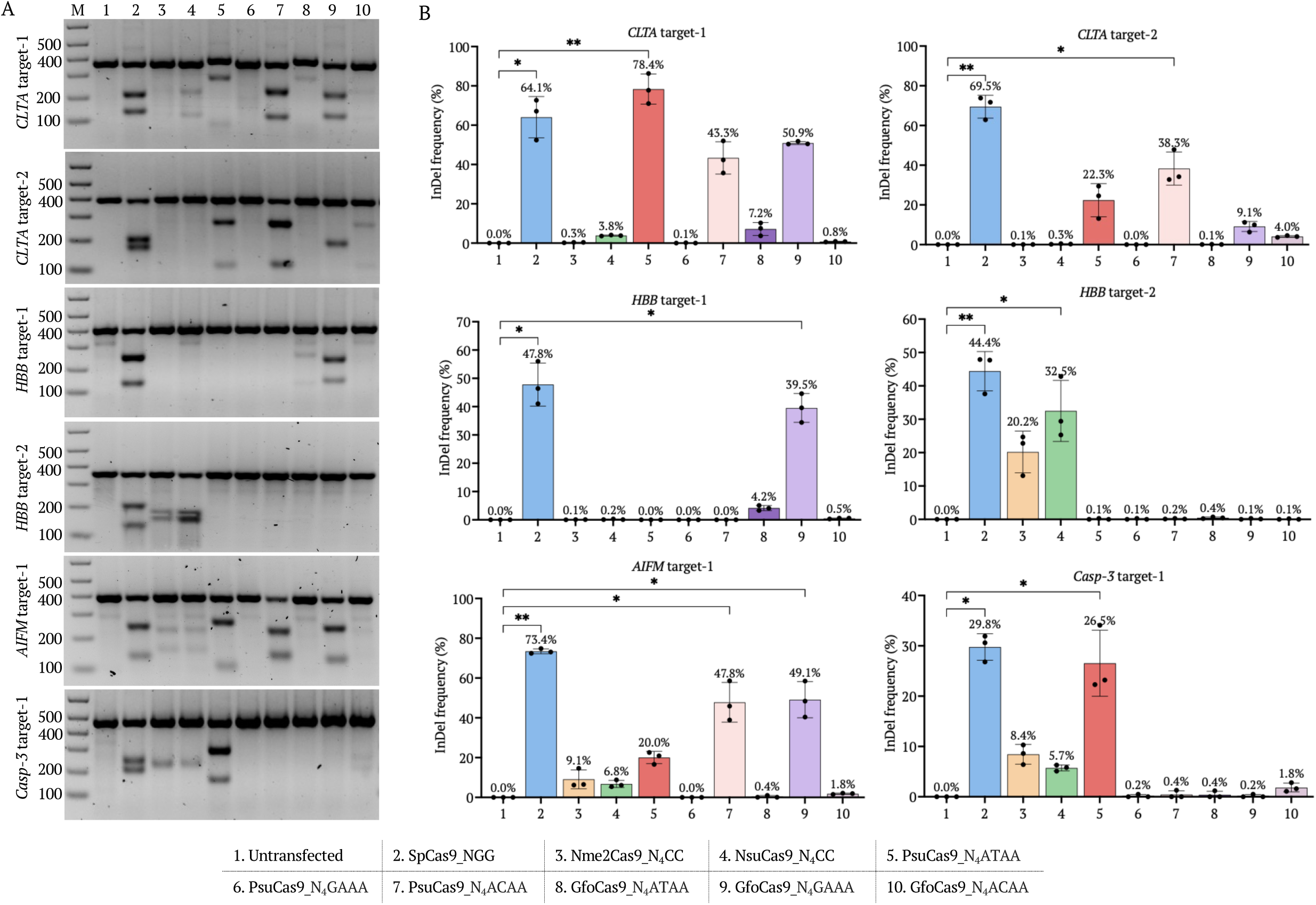
Editing of six endogenous human target sites by SpCas9, Nme2Cas9, NsuCas9, PsuCas9 and GfoCas9. (A) Representative T7EI gels for *CLTA* t1–t2, *HBB* t1–t2, *AIFM* t1, and *Casp-3* t1. Lanes: M, 1 kb plus marker; 1, untransfected; 2, SpCas9-NGG; 3, Nme2Cas9-N CC; 4, NsuCas9-N CC; 5, PsuCas9-N ATAA; 6, PsuCas9-N GAAA; 7, PsuCas9-N ACAA; 8, GfoCas9-N ATAA; 9, GfoCas9-N GAAA; 10, GfoCas9-N ACAA. (B) Indel frequencies (%) at each target by amplicon deep sequencing; bars, mean of three biological replicates; error bars, SEM; individual replicates plotted. Statistical significance: Kruskal–Wallis with Dunn’s post hoc; *p < 0.05, **p < 0.01, ***p < 0.001, ****p < 0.0001. See also Figures S9 and S10 and Tables S9–S15.

Amplicon deep sequencing quantified these editing outcomes precisely (Figure 3B). Among the type II-C editors, PsuCas9-N ATAA was the highest performer, exceeding SpCas9 at *CLTA* t1 (78.4% vs. 64.1% indels) and reaching 20.0%, 22.3% and 26.5% at *AIFM* t1, *CLTA* t2 and *Casp-3* t1, respectively. PsuCas9-N ACAA was moderately active at *CLTA* t1 (43.3%), *CLTA* t2 (38.3%), and *AIFM* t1 (47.8%), while PsuCas9-N GAAA was largely inactive. GfoCas9 displayed the opposite preference: GfoCas9-N GAAA reached 50.9%, 49.1%, and 39.5% at *CLTA* t1, *AIFM* t1, and *HBB* t1, respectively, whereas GfoCas9-N ATAA and-N ACAA were essentially inactive. NsuCas9 peaked at *HBB* t2 (32.5%), while Nme2Cas9 produced moderate editing only at *HBB* t2 (20.2%) and *AIFM* t1 (9.1%). SpCas9-NGG yielded indel frequencies of 29.8–73.4% across these loci. PsuCas9-N ATAA therefore matched or exceeded SpCas9 at the most permissive targets and consistently outperformed all other type II-C editors tested. To stress-test the standout enzyme/PAM combinations (PsuCas9-N ATAA, PsuCas9-N ACAA, and GfoCas9-N GAAA), we designed a second panel of seven targets that included a new locus, *EMX*, and trimmed the conditions to the three best-performing PAMs (Figure 4). T7EI assays again showed strong PsuCas9-N ACAA bands across all seven targets, while PsuCas9-N ATAA and GfoCas9-N GAAA showed locus-dependent activity (Figure 4A). Deep sequencing confirmed that PsuCas9-N ATAA achieved 77.5% editing at *EMX* t1 (vs. 63.9% for SpCas9) and 59.9% at *CLTA* t3 (vs. 55.5% for SpCas9), but only 7.0–13.0% at the two *HBB* t3/t4 sites. PsuCas9-N ACAA was modestly active at *CLTA* t4 (39.1%) and *Casp-3* t2 (32.4%), and GfoCas9-N GAAA peaked at *EMX* t1 (24.4%). Both Nme2Cas9 and NsuCas9 also showed their best activity at *EMX* t1 (18.3% and 22.9%, respectively), suggesting that local chromatin or sequence context, not merely PAM availability, strongly modulates type II-C activity (Figure 4B). The PAM ranking established in the initial screen was fully recapitulated, identifying PsuCas9-N ATAA and GfoCas9-N GAAA as the top type II-C performers in this expanded panel.

**Figure 4.**
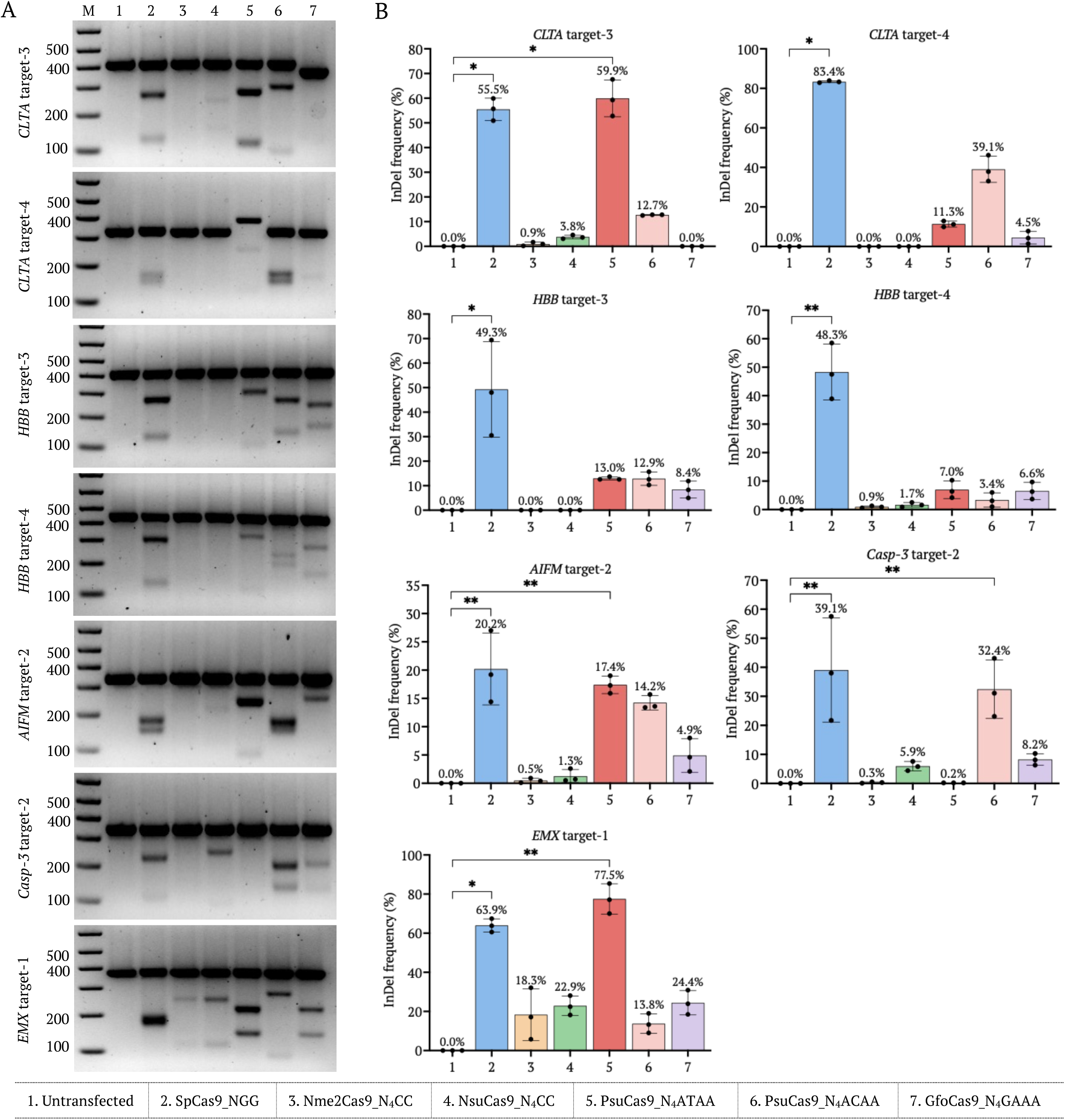
Editing of seven additional endogenous targets identifies PsuCas9-N ATAA as the top type II-C performer. (A) Representative T7EI gels for *CLTA* t3–t4, *HBB* t3–t4, *AIFM* t2, *Casp-3* t2 and *EMX* t1. Lanes: M, marker; 1, untransfected; 2, SpCas9-NGG; 3, Nme2Cas9-N CC; 4, NsuCas9-N CC; 5, PsuCas9-N ATAA; 6, PsuCas9-N ACAA; 7, GfoCas9-N GAAA. (B) Indel frequencies (%) by amplicon deep sequencing; bars, mean of three biological replicates; error bars, SEM. Statistical significance: Kruskal–Wallis with Dunn’s post hoc; *p < 0.05, **p < 0.01, ***p < 0.001, ****p < 0.0001.

### Type II-C orthologs generate a deletion-biased editing signature

To characterize repair outcomes, we performed in-depth mutation profiling at four representative loci (*CLTA* t2, *HBB* t1, *AIFM* t1, *EMX* t1). Deletions dominated the mutation spectrum of every type II-C ortholog at every target (Figure S9A). For PsuCas9 and GfoCas9 with their preferred PAMs, insertions typically accounted for less than 15% of editing outcomes, whereas SpCas9 generally produced a higher proportion of insertions, with insertion frequencies approaching deletion frequencies at some target sites. Substitutions were rare (<2%) regardless of the nuclease.

Aggregating across all targets (Figure S10), insertion-to-deletion ratios decreased from 0.73 for SpCas9 to 0.31, 0.26, 0.15, and 0.09 for Nme2Cas9, NsuCas9, PsuCas9, and GfoCas9, respectively, corresponding to a 2-to 8-fold stronger bias toward deletions than observed for SpCas9. Per-residue mutation maps showed that mutations cluster within 5–10 bp of the predicted cut site for all enzymes, with PsuCas9-N ATAA and GfoCas9-N GAAA producing the densest indel patterns at sites where they edit efficiently (Figure S9B). Indel size distributions were also distinct (Figure S9C): PsuCas9-N ATAA and GfoCas9-N GAAA generated deletions concentrated between −1 and −15 bp, whereas PsuCas9-N ACAA at AIFM t1 produced a longer-tailed distribution extending beyond −20 bp, suggesting that PAM-dependent cleavage or product-release dynamics may shape DNA repair outcomes. Nme2Cas9 and NsuCas9, when active, also favored small deletions. The deletion-biased outcomes shared by all three type II-C orthologs are well suited to gene-disruption applications, including the removal of regulatory elements or pathogenic insertions.

### No detectable off-target editing at 33 predicted sites

To assess specificity, we computationally predicted off-target sites for 11 productive on-target loci by searching the human reference genome (GRCh38/hg38) for sequences with the cognate PAM and 2–5 mismatches in the protospacer. The top 2–3 candidates per guide (33 sites in total) were amplified and analyzed by deep sequencing (Figure 5). None of the 33 sites showed editing above the untransfected baseline for PsuCas9-N ATAA, PsuCas9-N ACAA, GfoCas9-N GAAA, or NsuCas9-N CC, including off-targets bearing only two spacer mismatches (for example NsuCas9 at the *AIFM* locus). Two compounding factors plausibly account for this stringent on/off-target discrimination. First, the preferred PAMs of PsuCas9 and GfoCas9, such as 5’-N ATAA-3’ and 5’-N GAAA-3’, occur substantially less frequently than the canonical SpCas9 NGG PAM, thereby reducing the number of potential genomic target sites. Second, the requirement for extended PAM recognition together with spacer complementarity further restricts the pool of permissive off-target sites. Together, these properties account for the absence of detectable off-target editing across all five tested loci.

**Figure 5.**
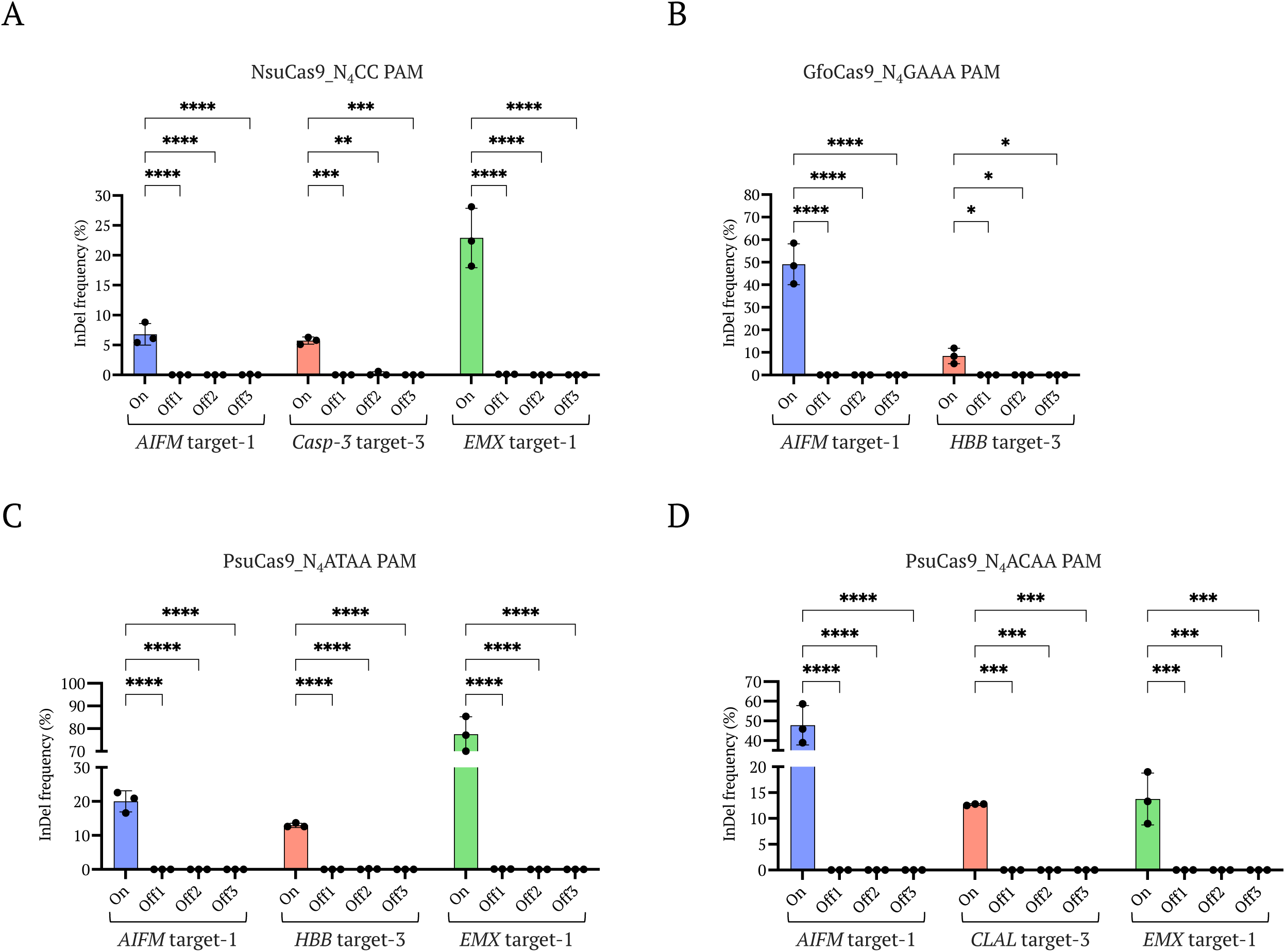
Off-target editing analysis of type II-C Cas9 orthologs at predicted genomic sites. Indel frequencies determined by amplicon deep sequencing at 11 on-target and 33 corresponding off-target sites for NsuCas9-N CC (A), GfoCas9-N GAAA (B), PsuCas9-N ATAA (C) and PsuCas9-N ACAA (D). Off-target sites were predicted by searching the human reference genome for matches with the cognate PAM and 2–5 spacer mismatches. Bars, mean (n = 3 biological replicates); error bars, SD; individual replicates plotted. Statistical significance: two-way ANOVA with Bonferroni’s post hoc; ns, not significant; *p < 0.05, **p < 0.01, ***p < 0.001, ****p < 0.0001. See also Tables S16 and S17.

## Discussion

PsuCas9 paired with its preferred 5’-N ATAA-3’ PAM matches or exceeds SpCas9 at multiple human loci and reaches indel frequencies up to 78.4%, establishing to our knowledge the first natural compact type II-C Cas9 ortholog reported to outperform SpCas9 in mammalian editing while retaining a sub-1,100 amino acid size, an orthogonal AT-rich PAM, and undetectable off-target activity at 33 predicted sites. This result challenges the long-held assumption that compactness in the type II-C subtype necessarily comes at the cost of catalytic potency in human cells^4, 18^ and reframes natural ortholog discovery, rather than directed evolution of established enzymes, as a productive route to high-performance compact editors. The accompanying enzymes NsuCas9 (5’-N CC-3’) and GfoCas9 (5’-N GAAA-3’) extend this conclusion to additional PAM landscapes, providing a complementary suite of compact, high-fidelity editors recovered from a single discovery pipeline.

PAM compatibility is the dominant determinant of editing efficiency for these enzymes. PsuCas9 and GfoCas9 share evolutionary ancestry yet display strikingly opposite PAM preferences (PsuCas9: N ATAA > N ACAA N GAAA; GfoCas9: N GAAA N ATAA ≈ N ACAA), implying that small differences in their PAM-interacting domains specify these distinct readouts. This natural pairing offers a tractable structural problem: co-crystal or cryo-EM structures of PsuCas9 and GfoCas9 RNPs bound to their cognate PAM duplexes will define the molecular basis of the specificity flip and can guide rational engineering of expanded-specificity variants in either direction. Such structure-guided engineering, layered onto the natural ortholog backbone, would further extend the AT-rich and GA-rich PAM space accessible to compact type II-C editors.

PsuCas9-N ATAA editing was strongly target-dependent, ranging from 78.4% at the most permissive sites to 0–13% at others. Such target-to-target variability is intrinsic to all Cas9 enzymes, including SpCas9 itself (which spanned 20.2–83.4% across the same panel), and likely reflects the combined contributions of local chromatin state, sequence context, and sgRNA folding. More distinctive is the repair signature: all three type II-C orthologs produced a pronounced deletion-biased mutation spectrum (insertion-to-deletion ratios 0.09–0.31 vs. 0.73 for SpCas9) and almost never produced the +1 templated insertion characteristic of SpCas9. The end-processing differences likely reflect both the geometry of the cleaved ends, which are blunt and located three nucleotides upstream of the PAM based on our cleavage-site mapping, and the kinetics of product release to the cellular repair machinery. For applications that disrupt rather than rebuild a sequence, including exon skipping, removal of pathogenic insertions, ablation of regulatory elements, or knockout screening, this deletion bias is a feature, not a liability.

The most consequential property of these enzymes for therapeutic development is their specificity. Across 33 computationally predicted off-target sites with two to five mismatches and matched cognate PAMs, none of the four high-activity enzyme/PAM combinations produced editing detectable above the untransfected baseline, including off-targets bearing only two spacer mismatches. Two factors may contribute to the stringent on/off-target discrimination observed for PsuCas9 and GfoCas9. First, their extended PAM requirements impose substantially greater sequence constraints than the canonical NGG PAM of SpCas9, thereby reducing the number of potential genomic binding sites. Second, these PAM constraints act in concert with guide RNA complementarity, further restricting the pool of permissive off-target sequences. Similar improvements in specificity associated with longer PAM requirements have been reported for Nme2Cas9 and SaCas9.^10, 18, 38^ Our 33-site validation provides strong but locally focused evidence; a definitive genome-wide specificity assessment by methods such as GUIDE-seq, DISCOVER-seq, or CHANGE-seq, and ultimately *in vivo* efficacy and safety studies in disease-relevant models, will be required before therapeutic application. With those caveats explicitly noted, the combination of compact size, robust on-target activity, deletion-biased repair, and the apparent specificity make PsuCas9 a particularly compelling candidate scaffold for AAV-deliverable therapeutic genome editing.

Several practical advantages emerge from the broader characterization. The asymmetric sgRNA cross-compatibility we observed argues against a single universal type II-C scaffold, but identifies the NsuCas9 sgRNA, whose extended stem-loop 1 supports activity across all three new orthologs, as a promising chassis for scaffold engineering. The wide thermal range of NsuCas9 (active up to 50–55 °C) and the unusual cold tolerance of PsuCas9 (active at 20 °C) suggest opportunities for low-temperature editing in primary cells, organoid cultures, and ectotherm systems where standard Cas9 enzymes are inefficient. The shared blunt-end cleavage three nucleotides upstream of the PAM positions all three enzymes for direct adaptation into base-editing, prime-editing, and epigenome-modulating fusions, where retention of catalytic activity at the predicted offset is essential.

More broadly, this work illustrates the value of large-scale embedding-based metagenomic mining for systematic genome-editing discovery. The DHR framework retrieves homologs that retain conserved structural features despite extensive sequence divergence and is in principle applicable to any RNA-guided nuclease class. Combined with predictive structural modeling and protein-language-model PAM inference, embedding-based search transforms a needle-in-the-haystack problem into a tractable triage pipeline. Given the continued growth of metagenomic databases, with MGnify alone expanding by more than an order of magnitude over the past five years, the pool of compact, specific, and orthogonally targetable Cas9 nucleases awaiting characterization is likely to remain large, and pipelines such as the one described here are well positioned to keep pace.

In summary, we report the discovery and comprehensive biochemical and cellular characterization of three previously uncharacterized compact type II-C Cas9 nucleases, NsuCas9, PsuCas9, and GfoCas9, identified through embedding-based mining of >4.7 × 10 metage omic proteins. PsuCas9 with its preferred 5’-N ATAA-3’ PAM achieves indel frequencies up to 78.4% in human cells and matches or exceeds SpCas9 at multiple endogenous loci, establishing the first natural compact type II-C ortholog reported to outperform SpCas9 in mammalian editing. All three enzymes show pronounced deletion-biased repair (insertion-to-deletion ratios 0.09–0.26 vs. 0.73 for SpCas9) and undetectable off-target editing across 33 predicted sites, including sites with only two spacer mismatches. Together, NsuCas9 (5’-N CC-3’), PsuCas9 (5’-N NYAA-3’), and GfoCas9 (5’-N RHAA-3’) provide a complementary set of compact, high-fidelity editors that substantially expand the sequence space accessible to CRISPR-Cas9. Their combination of small size, high on-target activity, expanded targeting range, deletion-biased repair, and undetectable off-target editing positions these enzymes, and PsuCas9 in particular, as compelling candidate scaffolds for AAV-deliverable therapeutic genome editing and for future adaptation into base-editor, prime-editor, and epigenome-editor derivatives.

## Materials and methods

### Identification and prioritization of Cas9 orthologs

Forty-nine representative Cas9 sequences spanning type II-A, II-B, and II-C subtypes^28^ were used to query the MGnify protein database (v2023.05) using a Dense Homology Retrieval framework. The top 10 hits per query were retrieved, deduplicated, and filtered to proteins ≤ 1,100 aa. Sequence-based annotation used HMM profiles of HNH, REC, RuvC-I/II/III and CTD domains constructed with MAFFT v7.487^39^ and HMMER v3.4;^40^ structure-based annotation used Foldseek^41^ against PDB Cas9 structures. Candidates lacking a complete RuvC catalytic triad or HNH domain were excluded. CRISPR arrays and tracrRNA elements were detected using CRISPRtracrRNA;^42^ candidates were retained when CRISPR-associated elements were within 1 kb of the Cas9 ORF. Sixteen candidates were structurally modelled with AlphaFold3^26^ and aligned to reference Cas9 structures using PyMOL v2.5.5 (Schrödinger, LLC) and ChimeraX;^43^ PAM preferences were predicted with Protein2PAM.^27^ Three orthologs (NsuCas9, PsuCas9, GfoCas9) were prioritized based on model confidence, structural similarity, and predicted PAM diversity.

### Phylogenetic analysis

The three new Cas9 proteins were aligned with 122 reference sequences spanning type II-A, II-B, II-C and II-D subtypes (Data S1). Sequences were clustered at 70% identity with CD-HIT v4.8.1,^44^ aligned with MAFFT v7.487 (L-INS-i, 1000 iterations),^39^ and trimmed with TrimAl v1.4.^45^ Maximum-likelihood phylogenies were inferred with IQ-TREE v2.2.0^46^ using the LG+F+R4 model selected by ModelFinder^47^ and assessed with 1,000 ultrafast bootstrap replicates. Trees were visualized in iTOL.^48^

### Protein expression and purification

Genes encoding NsuCas9, PsuCas9, GfoCas9, Nme2Cas9 and SpCas9 were codon-optimized for *E. coli*, synthesized as gBlocks (Integrated DNA Technologies, Coralville, IA, USA), and cloned into the pET-His-MBP-SUMO-SII backbone (Addgene #209290; Table S3). Proteins were expressed in *E. coli* Rosetta (DE3) at 30 °C for 4 h after induction with 0.25 mM IPTG, captured on HisTrap HP, cleaved with ULP1 protease, repassed over IMAC, and polished by HiTrap SP HP cation exchange and Superdex 200 size exclusion (GE Healthcare, Chicago, IL, USA). Purified proteins were flash-frozen and stored at −80 °C (Figure S4).

### sgRNA design and in vitro transcription

tracrRNA orientation and boundaries predicted by CRISPRtracrRNA^42^ were manually refined using canonical Rho-independent terminator features (GC-rich hairpin followed by ≥4 Ts) within 200 nt of the predicted anti-repeat.^49^ Direct-repeat and tracrRNA sequences were further validated by alignment to closely related orthologs (Figure S2). For each Cas9, sgRNAs were assembled by joining the direct repeat and tracrRNA via a 5’-GAAA-3’ tetraloop, with the repeat:anti-repeat duplex trimmed to 24 nt unless otherwise indicated (Table S2). Secondary structures were predicted with the RNAfold web server^50^ (Figure S3B). Designs that retained a stable repeat:anti-repeat duplex, a nexus, GC-rich stem-loops and a poly-U tail were used for subsequent functional assays. sgRNAs containing 20-nt or 24-nt spacers were designed based on PAM library targets and generated by *in vitro* transcription from PCR-amplified templates using HiScribe T7 Quick High Yield RNA Synthesis Kits (New England Biolabs, Ipswich, MA, USA; #E2050S), followed by RNA purification using the Monarch Spin RNA Cleanup Kit (New England Biolabs; #T2040L) according to the manufacturer’s instructions. The sgRNA templates and amplification primers are provided in Tables S4–S6.

### In vitro PAM screen and cleavage assays

The PAM library (Addgene #160132)^34^ contains a randomized 8-nt sequence flanking the protospacer. RNPs (150 nM) were assembled at 1.2:1 sgRNA:Cas9 at room temperature for 15 min, incubated with 1 µg of library DNA at 37 °C for 1 h, end-repaired with T4 polymerase, dA-tailed with Ex Taq, and ligated to a 3’-dT adapter. Adapter-tagged fragments were enriched by two PCR rounds with Illumina indexing and sequenced on an Illumina MiSeq. Reads were quality-filtered (Q ≥ 20), backbone-matched, and the 8-nt PAM extracted; the top 10% most-abundant PAMs were retained, normalized to a no-guide library, and visualized with Logomaker.^35^ For directed cleavage, RNPs (150 nM) were incubated with 200 ng XmnI-linearized plasmid in 20 µL 1× NEB Isothermal Amplification buffer at 37 °C for 45 min, treated with RNase A and Proteinase K, and resolved on 0.9% agarose gels. Sequence details for the *in vitro* targets, adapters and primers are listed in Tables S7–S8.

### Cleavage-site mapping

Two XmnI-linearized target plasmids per enzyme, each carrying a different high-activity PAM, were cleaved as above. Gel-purified fragments (QIAquick Gel Extraction Kit; QIAGEN, Hilden, Germany; #28704) were Sanger-sequenced and aligned in SnapGene to determine cleavage positions on both strands.

### Mammalian expression vectors and sgRNAs

Human-codon-optimized NsuCas9, PsuCas9 and GfoCas9 ORFs (Twist Bioscience; South San Francisco, CA, USA; Table S9) were cloned into the pX330 backbone (Addgene #52970) under the CMV promoter, replacing the original FokI-Cas9 ORF; SpCas9 (Addgene #87108) and Nme2Cas9 (Addgene #119923) plasmids were obtained directly. For guide RNA design, complementary oligos targeting five endogenous human loci (*CLTA*, *HBB*, *AIFM*, *Casp-3* [*Caspase-3*], and *EMX*) were designed and synthesized by Integrated DNA Technologies (Table S10). The sgRNA cassettes were assembled in U6-driven backbones with ortholog-matched scaffolds (Table S11), and the resulting U6-sgRNA cassettes were PCR-amplified for transfection using primers from Table S12. All sgRNA and PAM sequence details for each Cas9 ortholog across the target sites are summarized in Table S14.

### Cell culture, transfection and editing assays

HEK293T cells (passages 5–15) were maintained in DMEM with 10% FBS and 1% penicillin-streptomycin. Cells were seeded at 8 × 10 per well in 0.1%-gelatin-coated 24-well plates and transfected at 70% confluence using FuGENE 4K (Promega, Madison, WI, USA) with 250 ng Cas9 plasmid and 250 ng of U6-sgRNA amplicon per well. Cells were harvested 72 h after transfection and genomic DNA was extracted with the Monarch Genomic DNA Purification Kit (New England Biolabs #T3010L). On-target regions (Table S13) were PCR-amplified with primers in Table S15. The resulting amplicons were subjected to T7EI assays and analyzed on 1.5% agarose gels.

### Amplicon deep sequencing and statistics

Each target was amplified in three sequential PCRs (target-specific, adapter-overhang, and indexing). Pooled libraries were sequenced on an Illumina MiSeq, merged with FLASH, and analysed with CRISPResso2 v2.3.2.^51^ Indels <0.1% frequency or present in untransfected controls were discarded. Data are mean ± SEM of three biological replicates. Editing differences across Cas9 conditions at each target were tested with the Kruskal–Wallis test followed by Dunn’s post hoc correction. No statistical methods were used to predetermine sample size; experiments were not randomized; investigators were not blinded to allocation during experiments and outcome assessment.

### Off-target prediction and validation

Off-target sites were predicted by searching GRCh38/hg38 for sequences with the cognate PAM and ≤5 spacer mismatches; the top 2–3 sites per guide were selected (33 sites total; Tables S16– S17) and amplified from on-target genomic DNA. Reads were processed with CRISPResso2 as above, and indel frequencies were compared between off-target and matched on-target loci.

## Data availability

All next-generation sequencing data generated in this study have been deposited at the NCBI Sequence Read Archive (SRA) under BioProject accession PRJNA1439601 (SRA: PRJNA1439601) and are publicly available as of the date of publication. Plasmids constructed in this study are available from the corresponding author on reasonable request. All other data supporting the findings of this study are available within the article and its supplemental information.

## Supporting information

Cas9c_Dataset S1

## Acknowledgments

We thank all members of the Laboratory for Genome Engineering and Synthetic Biology at KAUST for useful discussions and technical assistance, and the KAUST Bioscience Core Lab for sequencing support. This work was supported by the King Abdullah University of Science and Technology (KAUST) baseline research fund BAS/1/1035-01-01 to M.M.M.

## Author contributions

M.M.M. conceived the research and supervised the project. Q.W., S.R.G. and A.S. designed the experiments. Q.W., S.R.G., R.A., A.S. and N.H. performed the experiments. A.M.K., Q.W., S.R.G., A.S. and M.M. carried out the statistical and bioinformatic analyses. M.M.M., Q.W. and S.R.G. analysed the data and wrote the manuscript with input from all authors. All authors read and approved the final manuscript.

