## Supplemental Material for "Compact type II-C Cas9 nucleases with expanded PAM access and high fidelity for therapeutic genome editing"

**Short title:** Compact high-fidelity type II-C Cas9 nucleases

**Supplemental Figures**

**Figure S1. Structural alignment of predicted Cas9 models with reference proteins**

The three-dimensional structures of NsuCas9 (A), PsuCas9 (B), and GfoCas9 (C) were predicted using AlphaFold3 ^1^ and structurally aligned with reference Cas9 proteins, including SaCas9 (PDB: 5AXW) ^2^, Nme1Cas9 (PDB: 6JDV) ^3^, and GeoCas9 (PDB: 8UZA) ^4^, using PyMOL (v2.5.5, Schrödinger, LLC). (D) Predicted TM-scores (pTM) for each model using AlphaFold3 and root-mean-square deviation (RMSD) values relative to the reference structures (PyMOL).

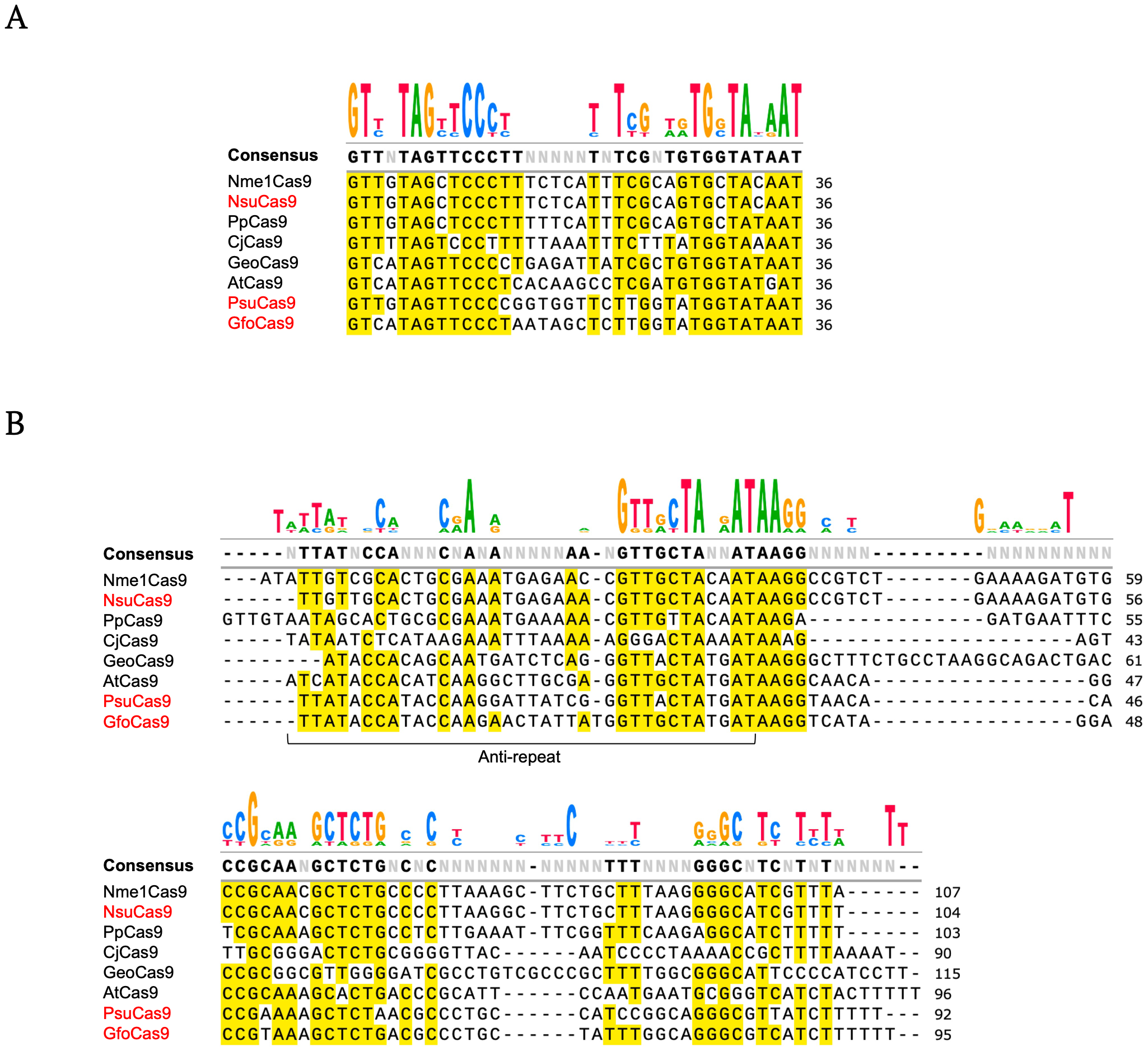

**Figure S2. Multiple sequence alignment of direct repeat and predicted tracrRNA sequences of type II-C Cas9 orthologs**

Direct repeat (DR) sequences (A) and tracrRNA sequences (B) from NsuCas9, PsuCas9, and GfoCas9 were identified using the CRISPRtracrRNA ^5^ pipeline and aligned with those of closely related Cas9 orthologs, including Nme1Cas9 ^6^, PpCas9 ^7^, CjCas9 ^8^, GeoCas9 ^4, 9^, and AtCas9 ^10^. Multiple sequence alignment was performed using MAFFT (v7.505) with the E-INS-i algorithm ^11^. Alignments were visualized in SnapGene, with conserved nucleotides highlighted in yellow. The newly identified candidates are labeled in red. The alignments reveal a high degree of conservation in both DR and tracrRNA sequences across type II-C Cas9 orthologs.

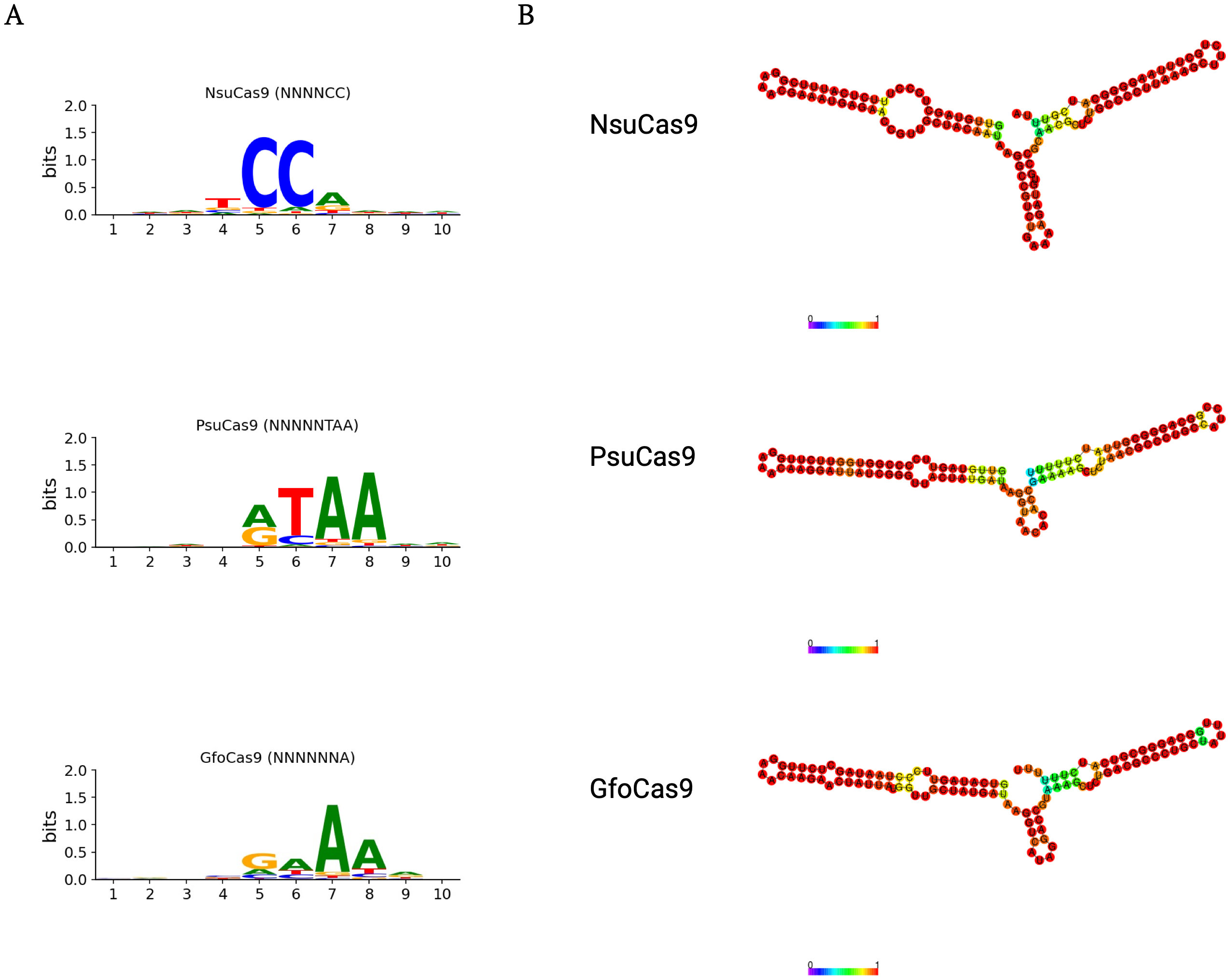

**Figure S3. Predicted PAM preferences and sgRNA secondary structures of new type II-C Cas9 orthologs**

(A) Predicted PAM preferences for NsuCas9 (top), PsuCas9 (middle), and GfoCas9 (bottom), generated using Protein2PAM v1.0^12^. PAM sequences used for subsequent in vitro assays were designed based on these predictions. (B) Centroid secondary structures of sgRNAs for NsuCas9 (top), PsuCas9 (middle), and GfoCas9 (bottom), predicted using the RNAfold web server ^13^ with the Andronescu energy model (2007) ^14^.

**Figure S4. Expression and purification of new type II-C Cas9 orthologs from E. coli**

(A) Schematic overview of the purification workflow applied to all Cas9 proteins. Proteins were purified using two sequential rounds of immobilized metal affinity chromatography (IMAC). (B) Representative SDS–PAGE gels showing protein fractions collected after the first (left) and second (right) rounds of IMAC. (C) Chromatograms showing absorbance (mAU) during the first (left) and second (right) rounds of IMAC. Red arrow indicates the 6×His–MBP–ULP1–Cas9 fusion protein; black arrow indicates cleaved Cas9 following ULP1 protease treatment; pink arrow indicates the cleaved 6×His–MBP–ULP1 tag. (D) SDS–PAGE analysis of purified NsuCas9, PsuCas9, and GfoCas9 proteins.

**Figure S5. Optimization of reaction conditions for *in vitro* cleavage assays**

(A) *In vitro* cleavage assays of NsuCas9, PsuCas9, and GfoCas9 performed in different buffer conditions, showing the effect of buffer composition on nuclease activity. (B) *In vitro* cleavage assays using ribonucleoprotein (RNP) complexes assembled with varying sgRNA-to-Cas9 ratios. (C) *In vitro* cleavage assays across a range of RNP concentrations to evaluate the effect of enzyme concentration on cleavage efficiency.

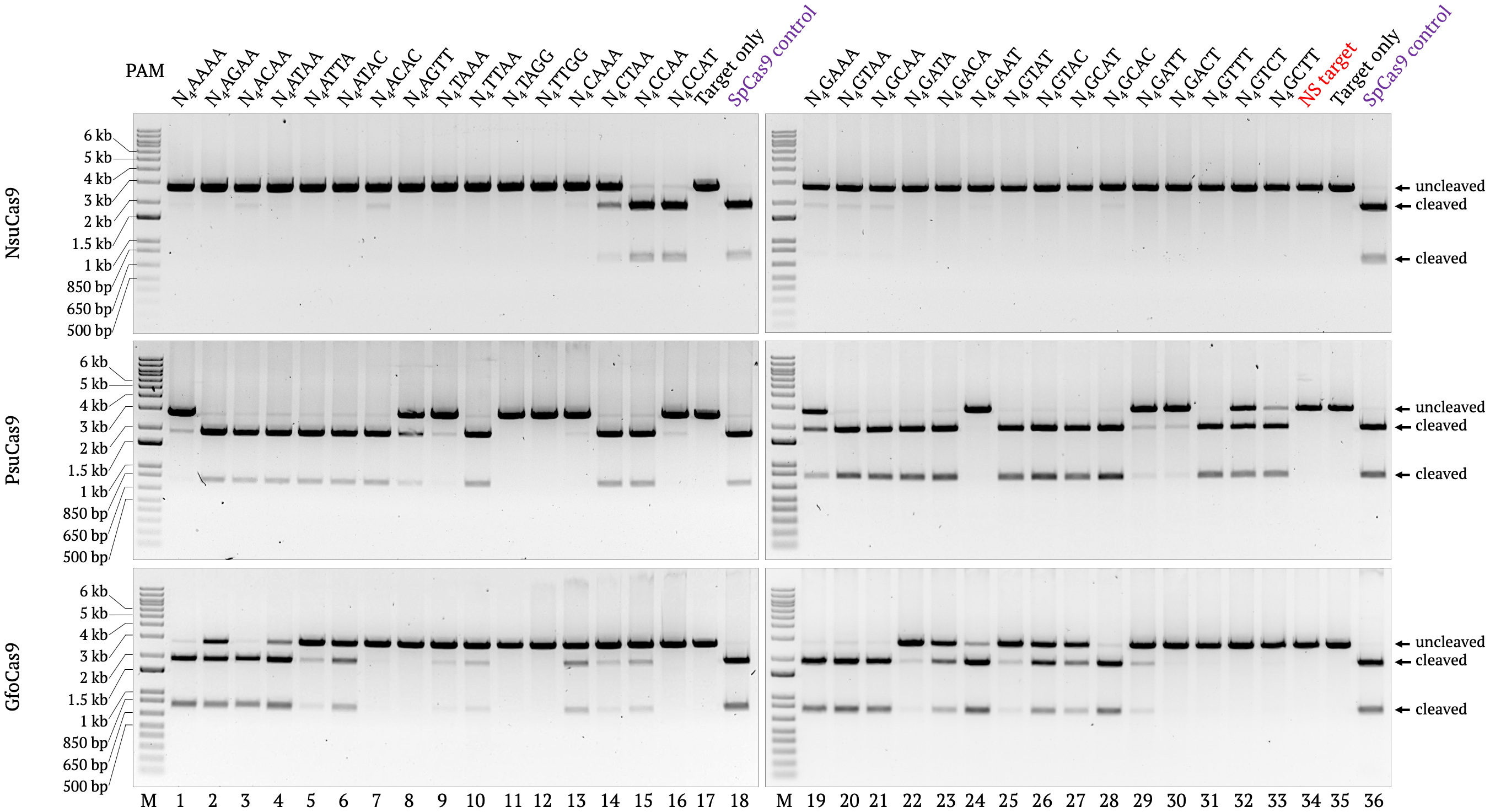

**Figure S6. *In vitro* validation of PAM preferences for compact type II-C Cas9 orthologs**

PAM preferences of NsuCas9 (top), PsuCas9 (middle), and GfoCas9 (bottom) were validated by *in vitro* cleavage assays using plasmid substrates containing representative PAM variants. dsDNA substrates were linearized with XmnI prior to Cas9 cleavage. SpCas9 was included as a positive control under identical conditions. “NS target” denotes a non-specific control plasmid lacking the cognate spacer.

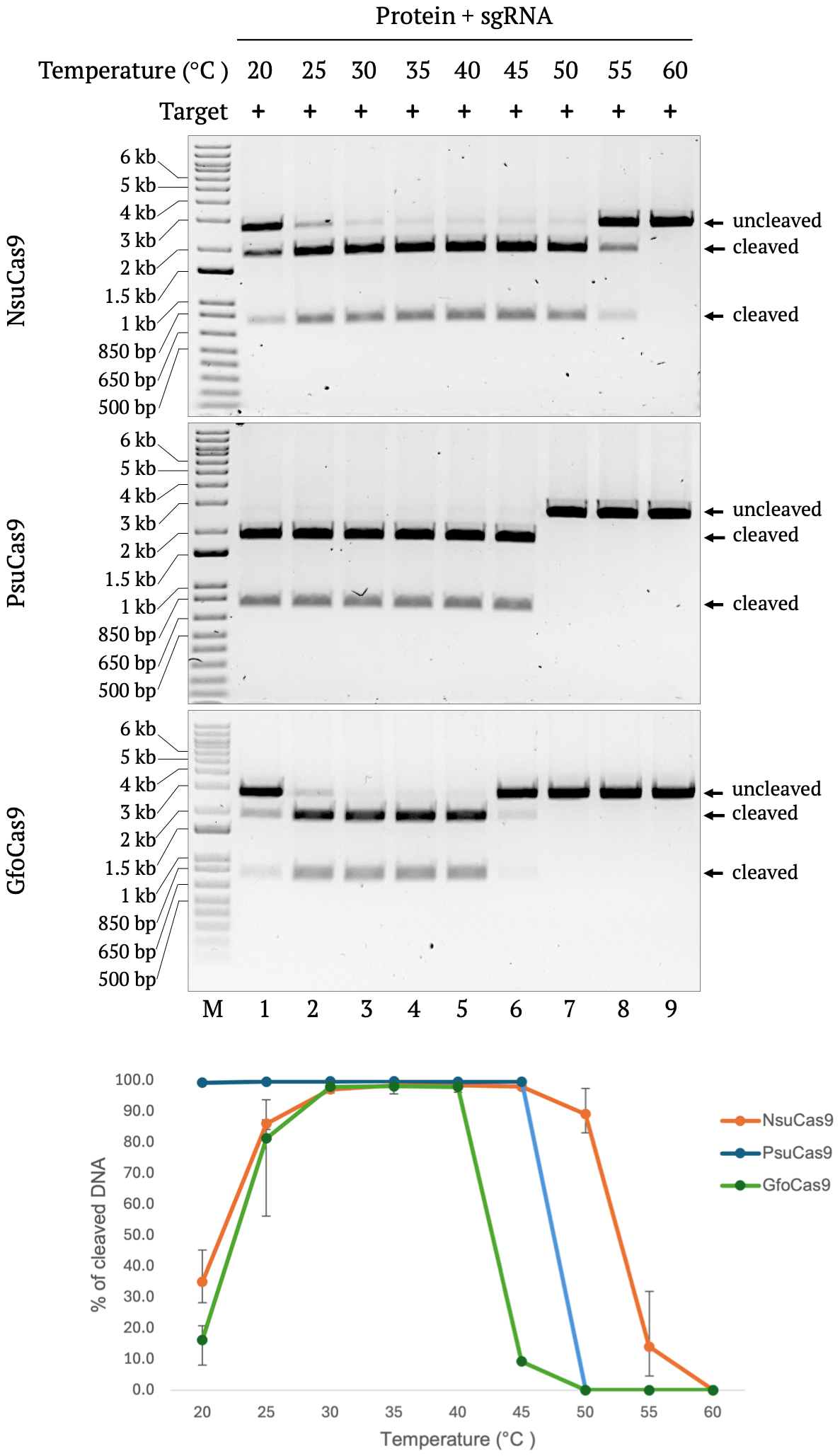

**Figure S7. Temperature-dependent cleavage of Type II-C Cas9 orthologs**

Representative gel images and quantification of *in vitro* cleavage by NsuCas9 (top), PsuCas9 (middle) and GfoCas9 (bottom) across temperatures from 20 °C to 60 °C. Quantification: mean ± s.d. of three independent experiments (n = 3).

**Figure S8. Cleavage site mapping of type II-C Cas9 orthologs on *in vitro* dsDNA targets**

*In vitro* cleavage site mapping was performed for NsuCas9 (A, B), PsuCas9 (C, D), and GfoCas9 (E, F). For NsuCas9, target plasmids containing PAM 20 (N₄CCAT) and PAM 11 (N₄CCAA) were used; for PsuCas9, PAM 29 (N₄ACAA) and PAM 23 (N₄ATAA); and for GfoCas9, PAM 27 (N₄GAAA) and PAM 23 (N₄ATAA). In each panel, the upper section shows a schematic of sgRNA-guided double-stranded DNA cleavage, with the target strand (TS) in red, the non-target strand (NTS) in black, and the PAM region highlighted in blue. Red arrows indicate cleavage positions determined by Sanger sequencing. The lower section shows representative Sanger sequencing chromatograms of purified cleavage products, visualized using SnapGene. Cleavage sites were identified by alignment to the reference plasmid sequence. Asterisks denote sequencing artifacts.

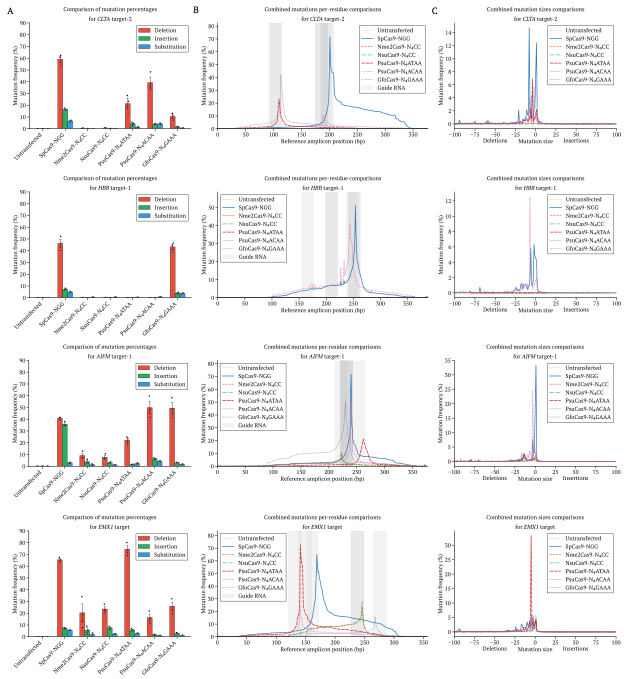

**Figure S9. Mutation profiles of Type II-C and benchmark Cas9 orthologs at four representative loci**

Composite mutation analysis at *CLTA* t2, *HBB* t1, *AIFM* t1 and *EMX* t1 across seven conditions: untransfected, SpCas9-NGG, Nme2Cas9-N₄CC, NsuCas9-N₄CC, PsuCas9-N₄ATAA, PsuCas9-N₄ACAA and GfoCas9-N₄GAAA. (A) Relative proportions of deletions, insertions and substitutions per condition (mean ± s.e.m., n = 3 biological replicates; individual replicate values plotted). (B) Per-residue mutation frequency across the PCR amplicon; gray shading marks the sgRNA target window; untransfected baseline subtracted. c Indel size distribution: positive values, insertions; negative values, deletions; mean of three biological replicates.

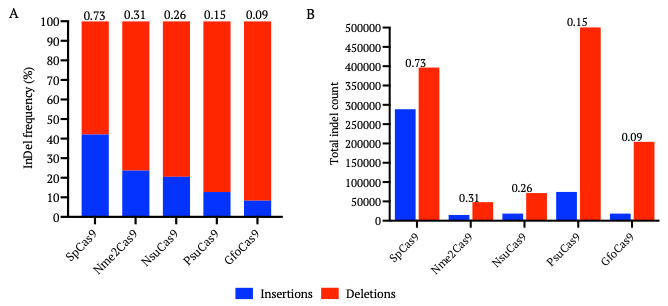

**Figure S10. Insertion-to-deletion analysis across all the Cas9 orthologs**

Aggregate insertion and deletion counts were calculated from amplicon deep sequencing data across all the target sites and biological replicates for each Cas9 orthologue. SpCas9 (NGG) was tested across 13 target sites (39 samples; 3 replicates each). Nme2Cas9 (N₄CC) and NsuCas9 (N₄CC) were each tested at 13 targets (39 samples each). PsuCas9 was tested with three PAM sequences (N₄ATAA, N₄GAAA, N₄ACAA) across multiple targets (96 samples), and GfoCas9 with three PAMs (N₄ATAA, N₄GAAA, N₄ACAA) across multiple targets (75 samples), resulting in higher total indel counts for these orthologs. (A) A stacked bar chart showing the proportion of insertions (blue) and deletions (red) for each Cas9 variant, normalized to 100%. Ins:Del ratios are indicated above each bar. All Type II-C orthologs display a deletion bias relative to SpCas9. (B) A grouped bar chart showing total insertion and deletion counts. Ins:Del ratios are indicated above each group. Note that PsuCas9 and GfoCas9 totals reflect testing across three PAM conditions per target, whereas SpCas9, Nme2Cas9, and NsuCas9 were each tested with a single PAM per target.
**Supplemental Tables**

**Table S1.** **Protein coding sequences of novel type II-C Cas9 orthologs**

| **Name** | **Sequence** |
| --- | --- |
| NsuCas9 (1092 aa) | MAAFKPNPINYIIGLDIGIASVGWAMVEIDEEENPIRLIDLGVRVFERAEVPKTGDSLAAARRLARSVRRLTRRRAHRLLRARRLLKREGVLQAADFDENGLIKSLPNTPWQLRAAALDRKLTPLEWSAVLLHLIKHRGYLSQRKNEGETADKELGALLKGVADNAHALQTGDFRTPAELALNKFEKESGHIRNQRGDYSHTFSRKDLQAELNLLFEKQKEFGNPHISDGLKEGIETLLMTQRDSIPDESAILRMQGKCTFEKNKSRAAKHSWSAERFIWLSKLANLRITDEFGTRKLTEIERQILIDLPYKLKTVNYQQVRNALTTLSPNALFNIRYHQKDKKGEIKSSDKIEKDTKFIEMKFWHKIKETLEGNGRKTEWQSLSSNHNLLDEIGTVFTICKNDERRKALLSGKIKEDQDILNILCNHVDFTCTVNLSLEALKNILPFMEQGDDYDTAWRKVYPPQSTKKESVLPPIPADEIRNPVVLRALSQARKVINSVVRRYGSPARIHIETAREVGKSFKDRKEIEKRQEENRKDREKAAAKFREYFPNFVGEPKSKDILKLRLYEQQHGKCLYSGKEINLGRLNEKGYVEIDHALPFSRTWDDSFNNKVLVLGSENQNKGNQTPYEYFNGKDNSREWQEFKARVETSRFPRSKKQRILLQKFDEDGFKERNLNDTRYVNRFLCQFVADHMLLTGKGKRRVFASNGQITNLLRGFWGLRKVRTENDRHHALDAVVVACSTVAMQQKITRFVRYKEMNAFDGKTIDKETGEVLHQKAHFPQPWEFFAQEVMIRVFGKPDGKPEFEEADTPEKLRTLLAEKLSSRPEAVHEYVTPLFVSRAPNRKMSGAHKDTLRSAKRFVKHNEKISVKRVWLTEIKLADLENMVNYKNGREIELYEALKARLGAYGGNAKQAFDPKDNPFYKKGGQLVKAVRVEKTQESGVLLNKKNAYTIADNGDMVRVDVFCKVDKKGKNQYFIVPIYAWQVAENILPDIDCKGYRIDDSYTFCFSLHKYDLIAFQKDEKSKVEFAYYINCDSSNGRFYLAWHDKGSKEQQFRISTQNLVLMQKYQIDELGKEIRPCRLKKRPPVR |
| PsuCas9 (1084 aa) | MKYAIGLDIGIASVGYAVLALDHEENPWGIIRLGSRIFDVAENPKDGASLALPRREARSVRRRLRRHHHRLERIKNLLINTELITKDELLHLYDGKLSDVYELRVKALDYSVTNAELTRILLHLAQRRGFKSNRKSDAGDKEAGQLLEAVSANAKRMQENHYRTVGEMFYKDDLFSKYKRNKGGTYLTTVHRDMIAAEARTILEKQFALGNDICTQDFIDKYLSILLSQRQFDEGPGEPSPYAGNQIANMIGKCTFEPNEYRAAKASYSFERFNLLQKVNHLRLLLEGKSIALDNEQRKKIIALAHDKADLRYSHIRKALGLDEKVLFNTITYNDDVSVIEKKTKFNFLQAYHQIKKVLAEDMQKLTTEQLDNIGQILSTYKSDNKRTEELSALGLEKKIIDALLGINGLSKFAHLSLKALRKINPYLEEGKIYNEACAAAGYDFKAHANTQKTELLPAYKEEMDDITSPVARRAIAQSIKVINAIIREQKCSPVYINIELAREMAKGFDERTQIDKANKENQAKNERIMERIRTEFHKSNPTGMDLIKLKLWEEQDGRSPYSQKAISINRLFEPGYVDIDHIVPYSISFDDSFKNKVLVFSDENRDKGNRLPIAYLQSKFGAKAAENFIIWVQSNIKDYKKRQKLLKREITEEDINKFKERNLQDTKTISRFLYNYINDYLLFAPSDTGKKKRVTAVNGTITAYLRKRWGINKIRANGDKHHAVDAVVIACTTDKMIKDLSSFSQYHELEYTHTDTESLLVNSLTGEILKRFPYPWEDFRPELMARLSDNPADALRKLNLIFYHGTDLSTIKPIFVSRMPRHKVTGAAHKATIKSARCLNNGIVICKTSLQNLKLDKEGEIANYYAPESDTILYNALKERLRAYDNKPAKAFAEPFYKPKADGTPGPLVKKVKVYEKSTLNVAVQQNTAVADNDSMVRVDVFHVKGDGYYLVPIYIADTLKKELPNKAIVAYKPYADWPVMDDSNFIFSLYPNDLIKFEHRNGVKFAKVNKESSLAETYLTKSELVYYKGTNITVGSTSIITNDNSYTVKSLGVKTLSNLEKYQVDVLGNYTKVKKEIRRPFR |
| GfoCas9 (1074 aa) | MKHPYGIGLDIGIASVGWAVVALNENAEPYGLIRCGSRIFDKAEQPKTGDSLAAPRREARSARRRLRRRSLRKADLYELMEKNGLPGKAEIEQAVQAGHLPDVYALRVQALDGPVTALDFARILLHLMQRRGFRSNRKADDAQKDGKLLQAIDANTRRMEANRYRTVGEMMYRDPVFAEHKRNKAENYLSTVKRDQIIDEARLVFAAQRQYGATWASPEMEAEYLCILTRQRSFAEGPGKGSPYSGSNRVGTCTLEGKSEQRAAKAAFSFEYFTLLQKINHIRIAENGTSRTLTPAERQVLLSACYQTDKLDFARIRKALALPEEARFNMVRYRGEQTAEACEKKEKITALPCYHKMRKALNTLRKDHIRNISREQLDAAGAALTNPENEDKLREALKQAEFEPLEIEALLTLPSFAGYGHISVKACRKLIPYLEQGMNYNDACQAAGYDFQGSQNGEKAQFLPASTEEMEDITSPVVRRAVAQTIKVVNAIIREQGESPVSIHLELAREMNKNFQQRSELDKAMRDNSAENERLMKELNELFPGRTVTGQDLVKYRLWKEQDGRCAYSIQPLELDKVITVSGYAEVDHIVPYSISFDDRRTNKVLVLASENRQKGNRLPLQYLQGKRRDDFIVYTKANVKNFRKRQNLLKERLSEEDGKGYIQRNLQDTQYIAAFMLNYIRNHLAFADCSGAGKRRVVAVNGAVTAFLRKRWGLSKVRADGDLHHAADAAVIACTTQGMIKRVSDFCKRAETTAVRNEHFPEPWPRFRDELTQRLSACPQENLMQINPVYYATVDISSIQPVFASRMPRHKVTGAAHKETIKSRLDDTHVVQRRNITELKLDKDGEIAGYFNRSSDTLLYNALKARLLAFGGDGKKAFAEPFYKPRADGTPGARVQKVKICDKVTSTVPVHGGKGVADNDTMVRMDVYYVPGDGYYWIPVYVADTVKPELPSKAVVAYKSSAEWKEMKDEDFLFSLCQHDLVRIESKRLMKFKVQNRDSTLEKEMPVNKILAYFEGGNISTGAITVTTHDDAYIADGLGFKTLQKVQKYQVDVLGNYTPVKKEKRQTFPAQRR |

**Table S2. Sequences of direct repeat, tracrRNA and sgRNA scaffolds**

| **Name** | **Sequence (5’---3’)** |
| --- | --- |
| NsuCas9 direct repeat | GTTGTAGCTCCCTTTCTCATTTCGCAGTGCTACAAT |
| NsuCas9 tracrRNA | TTGTTGCACTGCGAAATGAGAAACGTTGCTACAATAAGGCCGTCTGAAAAGATGTGCCGCAACGCTCTGCCCCTTAAGGCTTCTGCTTTAAGGGGCATCGTTTT |
| NsuCas9 sgRNA full-length scaffold | GTTGTAGCTCCCTTTCTCATTTCGGAAACGAAATGAGAAACGTTGCTACAATAAGGCCGTCTGAAAAGATGTGCCGCAACGCTCTGCCCCTTAAGGCTTCTGCTTTAAGGGGCATCGTTTT |
| PsuCas9 direct repeat | GTTGTAGTTCCCCGGTGGTTCTTGGTATGGTATAAT |
| PsuCas9 tracrRNA | TTATACCATACCAAGGATTATCGGGTTACTATGATAAGGTAACACACCGAAAAGCTCTAACGCCCTGCCATCCGGCAGGGCGTTATCTTTTT |
| PsuCas9 sgRNA full-length scaffold | GTTGTAGTTCCCCGGTGGTTCTTGGAAACAAGGATTATCGGGTTACTATGATAAGGTAACACACCGAAAAGCTCTAACGCCCTGCCATCCGGCAGGGCGTTATCTTTTT |
| GfoCas9 direct repeat | GTCATAGTTCCCTAATAGCTCTTGGTATGGTATAAT |
| GfoCas9 tracrRNA | TTATACCATACCAAGAACTATTATGGTTGCTATGATAAGGTCATAGGACCGTAAAGCTCTGACGCCCTGCTATTTGGCAGGGCGTCATCTTTTTT |
| GfoCas9 sgRNA full-length scaffold | GTCATAGTTCCCTAATAGCTCTTGGAAACAAGAACTATTATGGTTGCTATGATAAGGTCATAGGACCGTAAAGCTCTGACGCCCTGCTATTTGGCAGGGCGTCATCTTTTTT |

Red: sequences trimmed from the full-length sgRNA scaffold containing the 24-nt repeat:anti-repeat duplex.

Underline: 5’-GAAA-3’ tetraloop linker

**Table S3. E. coli codon-optimized coding sequences of type II-C Cas9 proteins**

| **Name** | **Sequence (5’---3’)** |
| --- | --- |
| NsuCas9 | AAGT**GGATCC**ATGGCAGCGTTCAAACCGAATCCGATCAACTATATTATTGGCTTAGACATCGGTATTGCCAGTGTGGGCTGGGCCATGGTTGAAATCGATGAAGAGGAGAATCCTATTCGCCTTATCGATTTAGGCGTGCGCGTGTTTGAACGCGCTGAAGTCCCTAAAACGGGAGATAGCTTAGCGGCAGCCCGCCGCTTAGCGCGCTCCGTGCGCCGTCTTACGCGTCGTCGTGCACATCGTCTGCTTCGCGCACGCCGTTTGTTAAAGCGCGAAGGTGTACTGCAGGCGGCGGATTTCGATGAAAACGGTCTGATTAAGTCACTGCCGAACACGCCCTGGCAACTGCGCGCCGCCGCCTTAGATCGTAAATTGACCCCCTTAGAATGGAGCGCCGTGCTTTTGCATCTTATCAAACACCGTGGGTACTTATCGCAGCGCAAAAACGAGGGCGAGACGGCCGATAAGGAATTGGGAGCACTGCTGAAAGGAGTGGCAGATAACGCACATGCCCTTCAAACTGGGGATTTTCGCACCCCGGCTGAGCTGGCCCTGAACAAATTTGAAAAAGAAAGCGGACACATTCGTAATCAACGCGGTGATTATTCGCACACCTTTTCTCGTAAGGACCTGCAAGCCGAATTGAACCTTCTGTTCGAGAAACAAAAAGAATTTGGCAATCCCCATATTAGTGACGGTCTGAAAGAAGGCATCGAAACGCTGCTGATGACGCAGCGCGACAGCATTCCCGACGAAAGCGCCATTTTGCGCATGCAAGGTAAGTGCACCTTTGAGAAAAACAAAAGCCGCGCGGCGAAACATTCGTGGAGCGCTGAGCGTTTCATCTGGCTTAGCAAACTGGCAAATCTCCGCATTACTGATGAATTTGGGACCCGCAAGCTGACAGAAATTGAGCGTCAAATATTAATCGACTTACCATACAAATTGAAAACCGTAAACTATCAACAGGTTCGCAATGCCCTTACAACGTTGTCCCCAAATGCACTTTTCAACATTCGATATCATCAGAAGGACAAGAAGGGAGAAATCAAATCAAGTGATAAAATTGAGAAAGATACCAAGTTCATCGAAATGAAATTCTGGCATAAAATTAAAGAAACCTTAGAGGGTAATGGCCGCAAAACGGAATGGCAAAGTCTCTCTAGTAACCATAATCTGTTAGATGAGATTGGGACCGTATTCACGATATGTAAAAATGACGAACGTCGTAAGGCATTGCTGTCTGGCAAAATCAAAGAAGATCAGGACATTTTAAACATTCTTTGCAATCACGTGGATTTCACGTGCACGGTAAATCTGAGCTTAGAGGCTCTGAAGAATATCCTTCCTTTCATGGAGCAGGGTGACGATTATGATACTGCGTGGAGAAAAGTCTACCCTCCTCAATCAACTAAAAAAGAGTCTGTTCTGCCGCCGATTCCGGCCGATGAGATACGCAACCCCGTAGTTCTTCGCGCACTTTCTCAGGCTCGTAAAGTCATCAATTCGGTGGTTCGGCGTTACGGCAGCCCGGCTCGCATACACATTGAAACAGCAAGAGAGGTTGGCAAATCATTTAAAGATCGGAAAGAAATTGAAAAACGCCAGGAAGAAAATAGAAAAGATCGTGAAAAAGCGGCCGCAAAATTTCGCGAATATTTCCCGAACTTTGTGGGTGAACCTAAATCGAAGGACATACTCAAACTTCGTTTATACGAACAACAGCACGGTAAATGTCTGTATAGCGGTAAAGAGATCAATTTAGGCCGCTTGAACGAAAAGGGCTATGTAGAGATCGATCATGCGTTACCATTCTCTCGCACTTGGGACGACAGCTTTAATAATAAAGTGCTGGTCCTTGGTTCGGAAAACCAGAATAAAGGTAATCAGACTCCATACGAATATTTCAATGGCAAGGACAATAGTCGCGAGTGGCAAGAGTTCAAGGCCCGTGTGGAGACTTCGCGTTTCCCGCGTTCTAAGAAACAGCGCATCCTGTTGCAGAAATTTGATGAAGACGGTTTCAAAGAAAGAAATCTGAATGATACACGTTATGTCAATCGTTTTCTGTGTCAGTTTGTTGCCGATCACATGCTTCTGACAGGGAAGGGCAAACGGCGTGTTTTCGCATCTAACGGGCAGATTACAAATCTCCTGCGCGGTTTTTGGGGTTTGCGGAAGGTGCGCACCGAGAATGATCGCCATCATGCCCTGGATGCTGTCGTAGTTGCCTGTTCTACGGTGGCAATGCAACAGAAGATCACTCGTTTTGTGCGTTACAAAGAGATGAATGCCTTTGATGGAAAAACGATAGATAAAGAGACAGGGGAAGTACTGCACCAAAAGGCCCATTTTCCACAGCCGTGGGAGTTTTTTGCACAAGAAGTTATGATTCGGGTCTTTGGAAAGCCAGATGGCAAACCAGAATTTGAAGAAGCTGATACACCGGAAAAACTTCGTACGCTGCTGGCAGAGAAACTCTCATCACGCCCGGAAGCTGTACATGAATATGTAACTCCCCTGTTTGTGTCTAGAGCACCGAATCGGAAAATGAGTGGTGCGCATAAAGATACGCTTCGCTCCGCCAAGCGTTTCGTGAAGCATAACGAAAAAATCTCAGTTAAACGCGTGTGGCTTACCGAGATCAAACTGGCAGATCTGGAGAATATGGTCAATTATAAAAACGGCCGTGAAATCGAGCTCTATGAAGCGTTGAAAGCCCGTCTGGGAGCATATGGTGGAAATGCGAAACAAGCGTTTGATCCTAAAGACAATCCCTTTTACAAGAAGGGAGGTCAGTTAGTTAAAGCCGTCCGTGTGGAAAAGACCCAGGAATCGGGCGTACTCTTAAACAAGAAAAATGCGTACACAATTGCCGATAACGGTGACATGGTTAGAGTGGATGTCTTTTGTAAAGTTGACAAAAAAGGTAAAAATCAGTATTTTATCGTGCCAATCTATGCCTGGCAAGTGGCAGAAAATATTCTTCCAGATATAGATTGCAAAGGCTATCGTATCGATGATTCTTATACCTTTTGTTTTTCACTGCATAAATATGATTTAATAGCATTTCAGAAAGATGAAAAATCAAAAGTGGAGTTTGCATACTATATCAATTGTGACAGTAGTAATGGACGTTTCTATTTGGCTTGGCATGACAAAGGCTCCAAAGAACAACAGTTTCGCATCTCCACGCAGAATTTAGTACTTATGCAGAAATATCAGATCGATGAATTAGGAAAAGAAATTCGCCCGTGCCGGCTGAAGAAACGCCCTCCAGTTCGCTAA**TTAATTAA**CCTA |
| PsuCas9 | AAGT**GGATCC**ATGAAATACGCAATTGGCCTTGACATTGGCATTGCCTCGGTAGGATATGCAGTGCTGGCGCTTGATCATGAAGAAAATCCTTGGGGCATCATTCGTCTTGGCTCCCGCATCTTCGATGTTGCAGAAAACCCGAAGGACGGAGCGTCGCTGGCGTTACCGCGCCGTGAAGCACGCAGTGTCCGCCGCCGCCTGCGTCGCCATCACCATCGCCTTGAGCGCATTAAAAATTTACTTATCAATACGGAGCTGATTACTAAAGATGAATTACTGCACCTGTATGATGGAAAGCTGTCAGACGTGTATGAACTGCGTGTCAAAGCCCTGGATTACAGTGTAACGAACGCGGAACTGACGCGTATCCTGCTGCATTTAGCACAGCGCCGTGGCTTTAAATCGAATCGCAAGTCGGATGCCGGAGATAAAGAGGCCGGGCAACTGCTGGAAGCCGTCAGCGCGAATGCGAAGCGTATGCAGGAGAACCACTATCGCACTGTGGGTGAAATGTTCTACAAAGACGATTTGTTTTCAAAGTACAAACGTAATAAGGGTGGCACCTATCTGACCACTGTTCATCGTGATATGATTGCAGCCGAAGCCCGTACTATTCTTGAAAAACAGTTCGCCCTGGGCAATGATATTTGTACGCAGGACTTCATTGATAAATACCTGAGCATTCTTCTTAGCCAGCGCCAGTTTGATGAGGGTCCAGGGGAACCAAGTCCGTACGCAGGCAACCAGATTGCGAACATGATCGGCAAATGTACCTTTGAACCGAATGAATACCGTGCAGCAAAAGCCAGTTACAGCTTTGAGCGCTTTAATCTGCTGCAGAAGGTAAACCATTTGCGCTTGCTTTTAGAGGGCAAATCAATCGCACTGGACAATGAGCAGCGCAAAAAAATCATCGCGCTGGCCCACGATAAAGCCGATTTGCGCTACAGTCATATTCGTAAAGCGCTGGGCCTTGACGAGAAAGTGTTGTTTAACACCATTACTTATAATGATGATGTGAGCGTAATCGAAAAAAAAACTAAATTTAATTTTCTTCAAGCTTATCATCAAATTAAAAAAGTTTTGGCTGAAGATATGCAGAAGCTGACAACAGAACAACTTGACAATATTGGGCAGATCCTGTCTACGTATAAATCCGATAATAAACGCACAGAGGAGCTGTCTGCCCTGGGATTGGAAAAAAAAATAATTGATGCACTGCTTGGCATCAACGGCCTCAGCAAATTCGCTCACTTATCGTTGAAGGCCCTGCGCAAAATTAATCCTTATCTTGAAGAAGGCAAAATTTATAATGAAGCATGCGCAGCGGCAGGCTATGATTTTAAAGCCCATGCAAATACCCAGAAAACGGAACTGCTGCCAGCTTATAAAGAAGAGATGGATGATATAACGTCCCCCGTGGCACGGCGCGCGATTGCGCAGTCAATCAAAGTGATTAATGCAATTATCCGCGAGCAGAAATGTTCCCCCGTCTATATCAATATCGAGCTCGCCCGCGAAATGGCAAAAGGCTTCGATGAACGAACGCAGATCGATAAGGCAAATAAGGAAAACCAAGCAAAAAATGAACGGATCATGGAACGTATTCGTACAGAGTTCCATAAAAGTAATCCAACCGGTATGGATCTGATTAAACTGAAGCTGTGGGAGGAACAGGATGGACGCTCTCCATACTCACAGAAAGCGATTAGCATTAACCGCTTGTTTGAGCCCGGCTACGTTGATATCGACCATATCGTTCCCTATAGCATCAGCTTTGACGATTCCTTTAAGAATAAAGTCTTAGTTTTTAGTGATGAGAATAGAGATAAAGGGAACCGGCTGCCAATTGCTTACTTGCAGTCGAAATTTGGCGCAAAAGCAGCAGAAAACTTCATCATATGGGTTCAGAGCAACATTAAAGACTATAAAAAGCGTCAGAAATTACTTAAAAGAGAAATCACTGAAGAAGATATAAATAAGTTTAAAGAACGCAACCTGCAGGATACAAAAACAATCTCACGGTTTCTCTATAATTATATAAACGATTACTTACTCTTTGCTCCGTCAGACACAGGGAAAAAAAAACGGGTTACCGCCGTAAATGGGACTATAACAGCATATTTACGGAAACGTTGGGGAATAAATAAAATTCGAGCTAACGGCGACAAGCATCATGCTGTCGATGCCGTAGTGATAGCCTGCACAACTGATAAAATGATTAAAGATCTGAGCTCCTTTTCTCAGTACCATGAATTAGAATATACACATACCGATACTGAGAGTCTGCTGGTTAACTCATTGACCGGCGAAATCCTGAAACGTTTTCCGTATCCATGGGAGGACTTCCGTCCGGAACTGATGGCCCGTTTATCCGACAATCCGGCGGACGCCTTACGTAAGCTTAACCTTATTTTCTACCACGGTACAGATCTGTCAACTATTAAACCTATTTTTGTATCTCGCATGCCGCGCCACAAAGTCACGGGAGCAGCACATAAAGCTACCATTAAGTCTGCCCGCTGTTTGAACAACGGCATTGTGATCTGTAAAACAAGCTTACAGAATCTGAAACTGGACAAGGAAGGTGAAATTGCCAATTATTATGCACCGGAGTCTGACACGATCCTTTATAATGCGCTCAAAGAACGCCTGCGTGCATATGATAACAAACCCGCTAAAGCCTTTGCGGAGCCGTTTTATAAACCTAAAGCCGACGGTACACCGGGCCCACTTGTTAAAAAGGTGAAAGTCTATGAGAAATCAACTTTGAACGTCGCAGTACAACAAAACACAGCCGTGGCTGACAACGATTCAATGGTGAGAGTGGACGTTTTCCACGTAAAAGGTGATGGTTATTATCTTGTCCCGATTTATATTGCTGACACGCTGAAAAAGGAATTGCCCAACAAGGCTATCGTGGCTTACAAACCGTATGCAGATTGGCCGGTAATGGATGATTCCAATTTTATTTTTTCTTTATATCCGAATGATCTGATTAAATTCGAACACCGCAATGGCGTGAAATTTGCAAAAGTTAATAAAGAATCGTCACTGGCGGAAACGTATCTCACTAAATCCGAACTCGTTTATTATAAGGGTACCAATATCACGGTCGGTTCTACGAGTATTATTACTAATGACAATTCTTATACTGTTAAGTCGTTAGGTGTGAAAACACTGAGCAATCTGGAGAAATATCAGGTCGACGTACTTGGAAATTATACTAAAGTGAAGAAGGAAATTAGACGTCCTTTTCG**TTAATTAA**CCTA |
| GfoCas9 | AAGT**GGATCC**ATGAAACATCCGTATGGGATTGGTTTAGACATCGGCATCGCCAGCGTCGGTTGGGCGGTCGTGGCTCTGAATGAGAATGCCGAACCGTATGGCTTGATCCGTTGTGGTTCTCGCATTTTTGACAAAGCCGAGCAACCGAAAACCGGGGACTCGCTGGCGGCTCCTCGTCGTGAAGCGCGTTCAGCCCGCCGTCGTCTGCGTCGTCGCAGCCTGCGTAAAGCAGATTTATACGAATTAATGGAGAAAAATGGCTTGCCGGGGAAAGCAGAAATTGAACAAGCGGTGCAAGCAGGCCACCTGCCCGATGTTTACGCCCTGCGCGTACAGGCTTTAGATGGCCCGGTTACCGCCCTGGATTTTGCCCGTATTTTATTACACCTGATGCAGCGTCGTGGCTTTCGTAGTAATCGTAAAGCGGATGACGCTCAAAAAGATGGCAAACTGCTGCAGGCCATTGACGCGAATACGCGCCGCATGGAAGCGAACCGCTACCGCACAGTTGGAGAAATGATGTACCGCGATCCGGTTTTTGCCGAACATAAACGCAATAAAGCCGAAAATTATCTGAGTACCGTGAAACGTGACCAAATTATTGATGAGGCACGCCTGGTATTTGCGGCGCAGCGCCAATACGGTGCGACCTGGGCATCCCCGGAGATGGAAGCAGAGTATTTGTGTATCTTAACCCGCCAGCGCTCTTTTGCCGAAGGCCCTGGCAAAGGTAGTCCGTATAGTGGGTCGAACCGCGTAGGCACATGCACGCTGGAGGGCAAAAGTGAACAGCGTGCTGCCAAAGCCGCCTTCTCATTTGAGTATTTCACACTGTTGCAGAAAATCAATCACATTCGTATTGCGGAAAATGGCACCTCTCGTACTCTGACGCCGGCAGAGCGTCAGGTTTTGCTTTCAGCGTGTTATCAGACCGACAAACTTGACTTTGCCCGCATTCGCAAAGCGCTGGCCTTGCCTGAAGAAGCCCGTTTTAACATGGTCCGTTATCGTGGGGAACAGACCGCCGAGGCGTGTGAAAAGAAAGAGAAAATCACCGCCTTACCATGCTATCATAAAATGAGAAAGGCACTGAATACCCTTCGCAAGGACCACATTCGGAATATTTCCCGTGAGCAGTTGGATGCAGCCGGTGCTGCCCTTACCAATCCCGAAAATGAGGACAAACTTCGTGAAGCGTTAAAACAGGCTGAATTCGAACCCTTGGAAATCGAAGCGTTACTGACGTTGCCCTCATTTGCGGGATACGGTCACATCTCAGTTAAAGCGTGCCGTAAACTCATTCCTTATCTCGAACAGGGGATGAATTACAATGATGCGTGTCAGGCCGCTGGTTATGACTTTCAGGGAAGTCAGAACGGAGAGAAAGCGCAGTTCTTACCGGCCTCGACAGAAGAAATGGAAGACATTACTTCCCCAGTCGTTCGACGCGCCGTTGCCCAGACGATTAAAGTCGTGAATGCCATTATACGCGAACAAGGGGAGTCGCCAGTAAGTATTCACCTGGAATTAGCTAGAGAGATGAATAAAAACTTTCAGCAACGCTCTGAGCTGGATAAAGCAATGCGCGATAACTCCGCAGAAAACGAACGGCTCATGAAAGAACTTAATGAATTATTTCCGGGGCGCACCGTTACGGGCCAGGATCTGGTTAAATACCGGCTTTGGAAAGAGCAGGACGGTCGGTGCGCATATTCTATACAACCACTGGAACTTGACAAAGTGATAACGGTCAGCGGCTATGCCGAGGTTGATCATATCGTTCCCTACTCTATCTCATTTGATGATCGGAGAACAAATAAGGTTCTCGTGTTGGCAAGTGAAAATCGACAGAAAGGGAACCGCCTGCCTTTGCAGTATCTGCAAGGGAAACGCAGAGACGATTTCATTGTCTACACAAAAGCAAACGTAAAAAATTTTCGGAAGAGACAGAACCTCCTGAAAGAGCGACTGAGTGAAGAGGACGGTAAGGGCTATATCCAGAGAAATCTTCAAGACACCCAGTATATCGCCGCGTTTATGCTGAATTACATTCGGAATCACCTGGCTTTTGCAGACTGCTCCGGCGCCGGGAAACGTCGTGTGGTTGCGGTTAACGGGGCAGTTACGGCTTTTCTTCGTAAACGGTGGGGCCTGAGTAAAGTTCGTGCAGACGGCGACTTACATCATGCCGCCGATGCAGCCGTTATCGCCTGCACAACCCAGGGTATGATTAAACGCGTTAGTGATTTCTGTAAAAGAGCGGAAACCACGGCCGTACGTAACGAACATTTTCCAGAACCGTGGCCTCGGTTTAGAGACGAATTAACTCAGCGTCTGAGCGCGTGTCCCCAGGAAAATTTAATGCAGATCAACCCAGTGTATTACGCCACCGTGGATATTAGTAGTATTCAGCCGGTCTTCGCATCGCGGATGCCCAGACACAAAGTGACCGGTGCGGCCCACAAAGAGACAATCAAGAGCAGACTGGATGATACACATGTGGTCCAACGTCGTAACATAACGGAATTAAAGCTTGATAAGGACGGTGAAATTGCTGGTTATTTTAACCGTTCGAGCGACACGTTGCTGTATAATGCGTTAAAAGCTAGATTGCTGGCCTTTGGAGGAGATGGCAAAAAGGCGTTCGCGGAACCGTTTTATAAGCCGAGAGCGGACGGTACACCGGGTGCACGTGTACAGAAGGTGAAAATTTGTGATAAGGTGACATCAACAGTACCTGTCCACGGTGGGAAGGGTGTGGCAGATAACGACACAATGGTTCGTATGGACGTATATTACGTTCCGGGTGATGGATACTACTGGATTCCAGTCTACGTTGCAGACACGGTAAAACCGGAGCTTCCATCCAAAGCAGTTGTAGCATACAAATCCAGCGCTGAATGGAAGGAAATGAAGGATGAGGATTTTCTGTTCTCCCTGTGCCAGCATGATTTGGTTCGCATTGAGAGTAAGCGTCTCATGAAATTTAAAGTGCAGAACCGTGACAGCACGCTGGAAAAAGAGATGCCAGTAAACAAAATTCTGGCGTATTTTGAAGGTGGGAATATTTCAACCGGGGCGATCACGGTGACCACTCACGATGACGCCTACATTGCTGACGGACTGGGATTTAAGACACTGCAAAAGGTTCAAAAGTATCAGGTCGATGTATTGGGGAATTATACCCCAGTGAAGAAAGAAAAACGCCAGACCTTTCCTGCGCAGCGTCGCTGA**TTAATTAA**CCTA |

Yellow highlighted: BamHI restriction site

Pink highlighted: PacI restriction site

**Table S4.** **sgRNA expression templates designed for cloning into a T7 promoter–driven vector for in vitro transcription**

| **Name** | **Sequence (5’---3’)** |
| --- | --- |
| NsuCas9_full-length sgRNA_20nt_spacer | CACCGCTAGCTAATACGACTCACTATAGGGCACGGGCAGCTTGCCGGGTTGTAGCTCCCTTTCTCATTTCGGAAACGAAATGAGAAACGTTGCTACAATAAGGCCGTCTGAAAAGATGTGCCGCAACGCTCTGCCCCTTAAGGCTTCTGCTTTAAGGGGCATCGTTTTAAAAAGCTTGGAT |
| NsuCas9_full-length sgRNA_24nt_spacer | CACCGCTAGCTAATACGACTCACTATAGGAGGGGCACGGGCAGCTTGCCGGGTTGTAGCTCCCTTTCTCATTTCGGAAACGAAATGAGAAACGTTGCTACAATAAGGCCGTCTGAAAAGATGTGCCGCAACGCTCTGCCCCTTAAGGCTTCTGCTTTAAGGGGCATCGTTTTAAAAAGCTTGGAT |
| PsuCas9_full-length sgRNA_20nt_spacer | CACCGCTAGCTAATACGACTCACTATAGGGCACGGGCAGCTTGCCGGGTTGTAGTTCCCCGGTGGTTCTTGGAAACAAGGATTATCGGGTTACTATGATAAGGTAACACACCGAAAAGCTCTAACGCCCTGCCATCCGGCAGGGCGTTATCTTTTTAAAAAGCTTGGAT |
| PsuCas9_full-length sgRNA_24nt_spacer | CACCGCTAGCTAATACGACTCACTATAGGAGGGGCACGGGCAGCTTGCCGGGTTGTAGTTCCCCGGTGGTTCTTGGAAACAAGGATTATCGGGTTACTATGATAAGGTAACACACCGAAAAGCTCTAACGCCCTGCCATCCGGCAGGGCGTTATCTTTTTAAAAAGCTTGGAT |
| GfoCas9_full-length sgRNA_20nt_spacer | CACCGCTAGCTAATACGACTCACTATAGGGCACGGGCAGCTTGCCGGGTCATAGTTCCCTAATAGCTCTTGGAAACAAGAACTATTATGGTTGCTATGATAAGGTCATAGGACCGTAAAGCTCTGACGCCCTGCTATTTGGCAGGGCGTCATCTTTTTTAAAAAGCTTGGAT |
| GfoCas9_full-length sgRNA_24nt_spacer | CACCGCTAGCTAATACGACTCACTATAGGAGGGGCACGGGCAGCTTGCCGGGTCATAGTTCCCTAATAGCTCTTGGAAACAAGAACTATTATGGTTGCTATGATAAGGTCATAGGACCGTAAAGCTCTGACGCCCTGCTATTTGGCAGGGCGTCATCTTTTTTAAAAAGCTTGGAT |
| NsuCas9_18nt-hybrid_scaffold | CACCGCTAGCTAATACGACTCACTATAGGGTCTTCGAGAAGACCTGTTGTAGCTCCCTTTCTCGAAAGAGAAACGTTGCTACAATAAGGCCGTCTGAAAAGATGTGCCGCAACGCTCTGCCCCTTAAGGCTTCTGCTTTAAGGGGCATCGTTTTAAAAAGCTTGGAT |
| NsuCas9_14nt-hybrid_scaffold | CACCGCTAGCTAATACGACTCACTATAGGGTCTTCGAGAAGACCTGTTGTAGCTCCCTTGAAAAACGTTGCTACAATAAGGCCGTCTGAAAAGATGTGCCGCAACGCTCTGCCCCTTAAGGCTTCTGCTTTAAGGGGCATCGTTTTAAAAAGCTTGGAT |
| PsuCas9_17nt-hybrid_scaffold | CACCGCTAGCTAATACGACTCACTATAGGGTCTTCGAGAAGACCTGTTGTAGTTCCCCGGTGGAAATATCGGGTTACTATGATAAGGTAACACACCGAAAAGCTCTAACGCCCTGCCATCCGGCAGGGCGTTATCTTTTTAAAAAGCTTGGAT |
| PsuCas9_14nt-hybrid_scaffold | CACCGCTAGCTAATACGACTCACTATAGGGTCTTCGAGAAGACCTGTTGTAGTTCCCCGGAAACGGGTTACTATGATAAGGTAACACACCGAAAAGCTCTAACGCCCTGCCATCCGGCAGGGCGTTATCTTTTTAAAAAGCTTGGAT |
| GfoCas9_17nt-hybrid_scaffold | CACCGCTAGCTAATACGACTCACTATAGGGTCTTCGAGAAGACCTGTCATAGTTCCCTAATAGAAATATTATGGTTGCTATGATAAGGTCATAGGACCGTAAAGCTCTGACGCCCTGCTATTTGGCAGGGCGTCATCTTTTTTAAAAAGCTTGGAT |
| GfoCas9_14nt-hybrid_scaffold | CACCGCTAGCTAATACGACTCACTATAGGGTCTTCGAGAAGACCTGTCATAGTTCCCTAGAAATATGGTTGCTATGATAAGGTCATAGGACCGTAAAGCTCTGACGCCCTGCTATTTGGCAGGGCGTCATCTTTTTTAAAAAGCTTGGAT |

Underline: Spacers

Yellow highlighted: T7 promoter

Pink highlighted: NheI restriction sites

Red hilighted: BbsI restriction sites

Blue highlited: DraI restriction sites

Green highlighted: HindIII restriction sites

**Table S5.** **Complementary oligonucleotide pairs used for cloning 20-nt or 24-nt spacers into BbsI-digested sgRNA backbone vectors for *in vitro* assays**

| **Oligonucleotide Name** | **Sequence (5’---3’)** |
| --- | --- |
| Spacer_20nt_invitro_TOP | TATAGGGCACGGGCAGCTTGCCGG |
| Spacer_24nt_invitro_TOP | TATAGGAGGGGCACGGGCAGCTTGCCGG |
| GTTG_20nt_invitro_BOTTOM | CAACCCGGCAAGCTGCCCGTGCCC |
| GTTG_24nt_invitro_BOTTOM | CAACCCGGCAAGCTGCCCGTGCCCCTCC |
| GTCA_20nt_invitro_BOTTOM | TGACCCGGCAAGCTGCCCGTGCCC |
| GTCA_24nt_invitro_BOTTOM | TGACCCGGCAAGCTGCCCGTGCCCCTCC |

**Table S6. Primers used for PCR amplification of the *in vitro* sgRNA expression templates and for the Sanger sequencing of sgRNA expression plasmids**

| **Primer Name** | **Sequence (5’---3’)** | **Used for** |
| --- | --- | --- |
| RTW443_T7_F | GCTAGCTAATACGACTCACTATAG | PCR amplification of T7-driven sgRNA expression templates, forward primer |
| NsuCas9_sgRNA_R | ACGATGCCCCTTAAAGCAG | PCR amplification of NsuCas9 sgRNA expression templates, reverse primer |
| PsuCas9_sgRNA_R | GATAACGCCCTGCCGGATG | PCR amplification of PsuCas9 sgRNA expression templates, reverse primer |
| GfoCas9_sgRNA_R | GATGACGCCCTGCCAAATAG | PCR amplification of GfoCas9 sgRNA expression templates, reverse primer |
| M13 Reverse | CAGGAAACAGCTATGAC | Sanger sequencing |
| M13 Forward | TGTAAAACGACGGCCAGT | Sanger sequencing |

**Table S7.** **Complementary oligonucleotide pairs used for cloning target sequences with variable PAMs into the pUC19 backbone**

| **Oligonucleotide Name** | **Sequence (5’---3’)** | **PAM (5’---3’)** |
| --- | --- | --- |
| PAM1_Top | AATTCGGGAGGGGCACGGGCAGCTTGCCGGTGGTTCTGATGCGG | TGGTTCTG |
| PAM1_Bottom | GATCCCGCATCAGAACCACCGGCAAGCTGCCCGTGCCCCTCCCG |  |
| PAM2_Top | AATTCGGGAGGGGCACGGGCAGCTTGCCGGCGGATTTGATGCGG | CGGATTTG |
| PAM2_Bottom | GATCCCGCATCAAATCCGCCGGCAAGCTGCCCGTGCCCCTCCCG |  |
| PAM3_Top | AATTCGGGAGGGGCACGGGCAGCTTGCCGGTAAATTGGATGCGG | TAAATTGG |
| PAM3_Bottom | GATCCCGCATCCAATTTACCGGCAAGCTGCCCGTGCCCCTCCCG |  |
| PAM4_Top | AATTCGGGAGGGGCACGGGCAGCTTGCCGGCAAATAGGATGCGG | CAAATAGG |
| PAM4_Bottom | GATCCCGCATCCTATTTGCCGGCAAGCTGCCCGTGCCCCTCCCG |  |
| PAM5_Top | AATTCGGGAGGGGCACGGGCAGCTTGCCGGAAAATTGGATGCGG | AAAATTGG |
| PAM5_Bottom | GATCCCGCATCCAATTTTCCGGCAAGCTGCCCGTGCCCCTCCCG |  |
| PAM6_Top | AATTCGGGAGGGGCACGGGCAGCTTGCCGGGTAAAGTTATGCGG | GTAAAGTT |
| PAM6_Bottom | GATCCCGCATAACTTTACCCGGCAAGCTGCCCGTGCCCCTCCCG |  |
| PAM7_Top | AATTCGGGAGGGGCACGGGCAGCTTGCCGGTGGTGCACATGCGG | TGGTGCAC |
| PAM7_Bottom | GATCCCGCATGTGCACCACCGGCAAGCTGCCCGTGCCCCTCCCG |  |
| PAM8_Top | AATTCGGGAGGGGCACGGGCAGCTTGCCGGTGGAGTACATGCGG | TGGAGTAC |
| PAM8_Bottom | GATCCCGCATGTACTCCACCGGCAAGCTGCCCGTGCCCCTCCCG |  |
| PAM9_Top | AATTCGGGAGGGGCACGGGCAGCTTGCCGGTAAAACACATGCGG | TAAAACAC |
| PAM9_Bottom | GATCCCGCATGTGTTTTACCGGCAAGCTGCCCGTGCCCCTCCCG |  |
| PAM10_Top | AATTCGGGAGGGGCACGGGCAGCTTGCCGGCGGAATACATGCGG | CGGAATAC |
| PAM10_Bottom | GATCCCGCATGTATTCCGCCGGCAAGCTGCCCGTGCCCCTCCCG |  |
| PAM11_Top | AATTCGGGAGGGGCACGGGCAGCTTGCCGGGGGTCCAAATGCGG | GGGTCCAA |
| PAM11_Bottom | GATCCCGCATTTGGACCCCCGGCAAGCTGCCCGTGCCCCTCCCG |  |
| PAM12_Top | AATTCGGGAGGGGCACGGGCAGCTTGCCGGAGGTATTAATGCGG | AGGTATTA |
| PAM12_Bottom | GATCCCGCATTAATACCTCCGGCAAGCTGCCCGTGCCCCTCCCG |  |
| PAM13_Top | AATTCGGGAGGGGCACGGGCAGCTTGCCGGGGGTGATTATGCGG | GGGTGATT |
| PAM13_Bottom | GATCCCGCATAATCACCCCCGGCAAGCTGCCCGTGCCCCTCCCG |  |
| PAM14_Top | AATTCGGGAGGGGCACGGGCAGCTTGCCGGGTTTGATAATGCGG | GTTTGATA |
| PAM14_Bottom | GATCCCGCATTATCAAACCCGGCAAGCTGCCCGTGCCCCTCCCG |  |
| PAM15_Top | AATTCGGGAGGGGCACGGGCAGCTTGCCGGCGGTGACAATGCGG | CGGTGACA |
| PAM15_Bottom | GATCCCGCATTGTCACCGCCGGCAAGCTGCCCGTGCCCCTCCCG |  |
| PAM16_Top | AATTCGGGAGGGGCACGGGCAGCTTGCCGGGGGAGACTATGCGG | GGGAGACT |
| PAM16_Bottom | GATCCCGCATAGTCTCCCCCGGCAAGCTGCCCGTGCCCCTCCCG |  |
| PAM17_Top | AATTCGGGAGGGGCACGGGCAGCTTGCCGGTAAAGCTTATGCGG | TAAAGCTT |
| PAM17_Bottom | GATCCCGCATAAGCTTTACCGGCAAGCTGCCCGTGCCCCTCCCG |  |
| PAM18_Top | AATTCGGGAGGGGCACGGGCAGCTTGCCGGCAAAGTTTATGCGG | CAAAGTTT |
| PAM18_Bottom | GATCCCGCATAAACTTTGCCGGCAAGCTGCCCGTGCCCCTCCCG |  |
| PAM19_Top | AATTCGGGAGGGGCACGGGCAGCTTGCCGGAAAAGTCTATGCGG | AAAAGTCT |
| PAM19_Bottom | GATCCCGCATAGACTTTTCCGGCAAGCTGCCCGTGCCCCTCCCG |  |
| PAM20_Top | AATTCGGGAGGGGCACGGGCAGCTTGCCGGGTTTCCATATGCGG | GTTTCCAT |
| PAM20_Bottom | GATCCCGCATATGGAAACCCGGCAAGCTGCCCGTGCCCCTCCCG |  |
| PAM21_Top | AATTCGGGAGGGGCACGGGCAGCTTGCCGGAGGAGAATATGCGG | AGGAGAAT |
| PAM21_Bottom | GATCCCGCATATTCTCCTCCGGCAAGCTGCCCGTGCCCCTCCCG |  |
| PAM22_Top | AATTCGGGAGGGGCACGGGCAGCTTGCCGGGGGACAAAATGCGG | GGGACAAA |
| PAM22_Bottom | GATCCCGCATTTTGTCCCCCGGCAAGCTGCCCGTGCCCCTCCCG |  |
| PAM23_Top | AATTCGGGAGGGGCACGGGCAGCTTGCCGGTCAAATAAATGCGG | TCAAATAA |
| PAM23_Bottom | GATCCCGCATTTATTTGACCGGCAAGCTGCCCGTGCCCCTCCCG |  |
| PAM24_Top | AATTCGGGAGGGGCACGGGCAGCTTGCCGGTGGCGTAAATGCGG | TGGCGTAA |
| PAM24_Bottom | GATCCCGCATTTACGCCACCGGCAAGCTGCCCGTGCCCCTCCCG |  |
| PAM25_Top | AATTCGGGAGGGGCACGGGCAGCTTGCCGGCTAAGTATATGCGG | CTAAGTAT |
| PAM25_Bottom | GATCCCGCATATACTTAGCCGGCAAGCTGCCCGTGCCCCTCCCG |  |
| PAM26_Top | AATTCGGGAGGGGCACGGGCAGCTTGCCGGGCAAGCATATGCGG | GCAAGCAT |
| PAM26_Bottom | GATCCCGCATATGCTTGCCCGGCAAGCTGCCCGTGCCCCTCCCG |  |
| PAM27_Top | AATTCGGGAGGGGCACGGGCAGCTTGCCGGTGGTGAAAATGCGG | TGGTGAAA |
| PAM27_Bottom | GATCCCGCATTTTCACCACCGGCAAGCTGCCCGTGCCCCTCCCG |  |
| PAM28_Top | AATTCGGGAGGGGCACGGGCAGCTTGCCGGGTTTGCAAATGCGG | GTTTGCAA |
| PAM28_Bottom | GATCCCGCATTTGCAAACCCGGCAAGCTGCCCGTGCCCCTCCCG |  |
| PAM29_Top | AATTCGGGAGGGGCACGGGCAGCTTGCCGGCGGCACAAATGCGG | CGGCACAA |
| PAM29_Bottom | GATCCCGCATTTGTGCCGCCGGCAAGCTGCCCGTGCCCCTCCCG |  |
| PAM30_Top | AATTCGGGAGGGGCACGGGCAGCTTGCCGGCACGAAAAATGCGG | CACGAAAA |
| PAM30_Bottom | GATCCCGCATTTTTCGTGCCGGCAAGCTGCCCGTGCCCCTCCCG |  |
| PAM31_Top | AATTCGGGAGGGGCACGGGCAGCTTGCCGGTACCTTAAATGCGG | TACCTTAA |
| PAM31_Bottom | GATCCCGCATTTAAGGTACCGGCAAGCTGCCCGTGCCCCTCCCG |  |
| PAM32_Top | AATTCGGGAGGGGCACGGGCAGCTTGCCGGGGCACTAAATGCGG | GGCACTAA |
| PAM32_Bottom | GATCCCGCATTTAGTGCCCCGGCAAGCTGCCCGTGCCCCTCCCG |  |
| PAM33_Top | AATTCGGGAGGGGCACGGGCAGCTTGCCGGAGCGTAAAATGCGG | AGCGTAAA |
| PAM33_Bottom | GATCCCGCATTTTACGCTCCGGCAAGCTGCCCGTGCCCCTCCCG |  |
| PAM34_Top | AATTCGGGAGGGGCACGGGCAGCTTGCCGGTAGTAGAAATGCGG | TAGTAGAA |
| PAM34_Bottom | GATCCCGCATTTCTACTACCGGCAAGCTGCCCGTGCCCCTCCCG |  |

**Table S8. Adapters and PCR primers used to enrich cleaved and uncleaved dsDNA fragments in *in vitro* PAM characterization for deep sequencing**

| **Primer Name** | **Sequence (5’---3’)** | **Used for** |
| --- | --- | --- |
| TK-117 | CGGCATTCCTGCTGAACCGCTCTTCCGATCT | Generation of an adapter with a 3’ dT overhang for capturing cleaved dsDNA fragments, adapted from Karvelis et al. ^15^ |
| TK-111 | GATCGGAAGAGCGGTTCAGCAGGAATGCCG | Generation of an adapter with a 3’ dT overhang for capturing cleaved dsDNA fragments, adapted from Karvelis et al. ^15^ |
| Library_RTW554_Cut_F1 | GGCGTTTCACTTCTGAGTTCGGC | PCR amplification of cleaved dsDNA fragments |
| Library_RTW554_Cut_F2 | CAGACCGCTTCTGCGTTCTG | PCR amplification of cleaved dsDNA fragments |
| Library_RTW554_Cut_F3 | TCGCAACGTTCAAATCCGCTCC | PCR amplification of cleaved dsDNA fragments |
| Library_RTW554_UC_R1 | GCCTGACTCACTATAGGGAGACCG | PCR amplification of uncleaved dsDNA fragments |
| Library_RTW554_UC_R2 | CGGTATTTCACACCGCATACGTACG | PCR amplification of uncleaved dsDNA fragments |
| F1a_library | TCGTCGGCAGCGTCAGATGTGTATAAGAGACAGAAAACACACCGCATACGTACGATTTA | Addition of Illumina adapters to uncleaved samples, adapted from Pedrazzoli et al. ^16^ |
| F4a_invitro | TCGTCGGCAGCGTCAGATGTGTATAAGAGACAGCTGCTGAACCGCTCTTCCGATC | Addition of Illumina adapters to cleaved samples, adapted from Pedrazzoli et al. ^16^ |
| F5a_library | TCGTCGGCAGCGTCAGATGTGTATAAGAGACAGCGTACGATTTAAATAGGCCTGACT | Addition of Illumina adapters to uncleaved samples |
| R1_library | GTCTCGTGGGCTCGGAGATGTGTATAAGAGACAGCGTTCTGATTTAATCTGTATCAGGC | Addition of Illumina adapters to cleaved samples, adapted from Pedrazzoli et al. ^16^ |
| R2_library_Cut | GTCTCGTGGGCTCGGAGATGTGTATAAGAGACAGTCCTACTCAGGAGAGCGTTCAC | Addition of Illumina adapters to cleaved samples |
| R3_library_UC | GTCTCGTGGGCTCGGAGATGTGTATAAGAGACAGCAGTCTTTCGACTGAGCCTTTCG | Addition of Illumina adapters to uncleaved samples |

**Table S9. Human codon-optimized type II-C Cas9 coding sequences**

| **Name** | **Sequence (5’---3’)** |
| --- | --- |
| NsuCas9 | TCT**GGATCC**ATGGCCGCCTTCAAGCCCAATCCTATCAACTACATCATCGGCCTGGACATCGGCATTGCCTCCGTGGGATGGGCTATGGTGGAAATCGACGAGGAAGAAAACCCTATCCGGCTGATCGACCTGGGCGTGAGAGTGTTTGAGCGCGCCGAAGTGCCCAAGACCGGCGATAGCCTGGCCGCTGCCCGTAGGCTGGCTCGAAGCGTGCGACGGCTGACACGGCGGAGAGCCCACAGGCTGCTGAGAGCCAGACGGCTGCTGAAGAGAGAGGGCGTGCTGCAGGCCGCTGATTTCGACGAAAACGGTCTGATCAAGAGCCTGCCTAATACCCCCTGGCAGCTGAGAGCCGCTGCACTCGACAGGAAGCTGACCCCACTGGAATGGTCCGCCGTGTTGCTGCACCTGATCAAGCACCGAGGCTACCTGTCTCAGAGAAAGAACGAGGGCGAAACCGCTGATAAGGAACTGGGCGCCCTGCTGAAGGGCGTGGCCGATAACGCCCATGCCCTGCAAACCGGAGATTTCAGAACGCCTGCCGAGCTGGCTCTGAACAAGTTCGAGAAAGAATCTGGCCACATTAGAAATCAGCGGGGCGATTACAGCCACACATTCAGCAGAAAAGACCTGCAGGCCGAGCTGAATCTGCTGTTCGAAAAACAGAAAGAGTTCGGCAACCCCCACATCAGCGATGGACTGAAAGAGGGAATCGAGACACTGCTGATGACCCAGAGGGACTCTATCCCCGATGAGTCCGCCATCCTGAGAATGCAAGGCAAGTGCACATTCGAGAAGAACAAGAGCCGGGCCGCTAAGCACAGCTGGTCAGCCGAGAGATTCATCTGGCTGTCTAAGCTGGCCAACTTACGGATTACTGATGAGTTCGGCACCCGGAAGCTGACCGAGATCGAGAGACAGATCCTGATCGACCTGCCATACAAACTCAAGACCGTGAACTACCAGCAAGTGCGTAATGCCCTGACCACCCTGTCACCTAACGCCCTGTTTAACATCAGATACCACCAGAAGGACAAGAAGGGCGAGATCAAATCCAGCGACAAGATTGAAAAGGACACCAAGTTCATCGAGATGAAGTTCTGGCATAAGATCAAGGAAACCCTGGAGGGCAACGGAAGAAAGACCGAGTGGCAAAGCCTGTCCAGCAACCATAACCTGCTGGATGAAATCGGCACCGTGTTCACAATCTGCAAGAACGACGAGAGAAGAAAGGCTCTGCTGAGCGGCAAAATCAAGGAGGATCAGGACATCCTTAATATCCTGTGCAACCACGTGGACTTTACCTGTACCGTGAATTTGAGCCTGGAAGCTCTGAAAAACATCCTGCCTTTCATGGAACAGGGCGACGACTACGACACCGCCTGGCGGAAGGTGTACCCTCCTCAGTCTACAAAGAAAGAGAGCGTGCTGCCTCCAATCCCAGCCGACGAGATCAGAAACCCCGTCGTGCTCAGAGCCCTGTCACAGGCTCGGAAGGTGATCAACAGCGTGGTGCGGCGTTACGGCTCTCCCGCCAGAATCCACATCGAAACCGCCAGAGAAGTGGGCAAGAGCTTCAAGGATAGAAAGGAAATCGAGAAAAGACAGGAGGAAAACCGGAAAGACCGGGAAAAAGCCGCGGCCAAATTCCGGGAATATTTCCCTAACTTCGTGGGCGAGCCTAAGAGCAAGGATATCCTGAAGCTGAGACTGTACGAGCAACAACACGGCAAGTGCCTGTACAGCGGAAAAGAGATCAACTTAGGGAGACTGAATGAAAAAGGCTACGTTGAGATCGACCACGCCCTGCCCTTCAGTAGAACCTGGGACGACAGCTTCAACAATAAGGTCCTGGTCCTGGGCAGCGAGAACCAGAACAAGGGCAACCAGACCCCCTACGAGTACTTCAACGGCAAGGATAATAGCAGAGAGTGGCAGGAGTTCAAGGCAAGAGTGGAAACAAGCCGGTTTCCTAGATCTAAGAAGCAGCGGATTCTGTTGCAAAAATTCGACGAGGATGGCTTCAAAGAAAGAAACCTGAACGACACAAGATATGTGAACCGCTTTCTCTGTCAGTTCGTGGCCGATCACATGCTGCTGACCGGCAAGGGCAAGCGAAGAGTATTTGCCAGCAACGGGCAGATCACCAACTTGCTGCGGGGCTTTTGGGGTCTGCGGAAGGTGAGAACCGAGAACGACAGACACCACGCCCTGGACGCCGTGGTGGTGGCCTGCTCCACGGTGGCTATGCAGCAAAAGATCACCAGATTCGTGAGATACAAGGAGATGAACGCCTTCGATGGAAAGACAATTGACAAAGAGACAGGAGAGGTGCTGCACCAGAAGGCCCACTTCCCCCAGCCTTGGGAGTTTTTTGCCCAGGAGGTGATGATCAGAGTGTTCGGCAAGCCCGACGGCAAGCCTGAGTTCGAAGAGGCAGATACCCCTGAGAAGCTGCGGACACTGCTGGCCGAGAAGCTGTCCAGCAGACCTGAGGCCGTGCACGAGTACGTGACCCCACTCTTCGTGTCCAGGGCCCCTAACAGAAAGATGAGCGGCGCCCACAAGGACACACTGAGAAGCGCCAAGAGATTCGTTAAGCACAACGAGAAGATCAGCGTGAAGCGGGTGTGGCTGACAGAGATCAAGCTGGCCGACCTGGAAAACATGGTCAACTACAAAAACGGCAGAGAAATCGAACTGTACGAAGCTCTCAAGGCCAGACTGGGCGCTTATGGCGGAAACGCCAAGCAGGCCTTCGACCCTAAGGACAACCCCTTCTACAAAAAGGGCGGACAGCTGGTGAAGGCCGTGAGAGTGGAAAAGACACAGGAGAGCGGCGTCCTGCTGAACAAAAAGAATGCCTACACCATCGCCGACAACGGCGACATGGTGCGGGTGGATGTGTTCTGCAAGGTGGACAAAAAGGGCAAGAATCAGTACTTCATCGTGCCTATCTACGCCTGGCAGGTGGCCGAGAACATCCTGCCCGACATCGATTGCAAGGGCTATCGGATCGACGATTCTTATACCTTCTGCTTCAGCCTGCACAAGTACGACCTGATCGCCTTTCAGAAAGATGAAAAGAGCAAGGTGGAGTTCGCCTACTACATAAACTGCGACAGCAGCAACGGCAGATTCTACCTGGCCTGGCACGACAAGGGATCTAAGGAACAGCAGTTTAGAATCTCTACCCAAAATCTGGTGCTGATGCAGAAGTACCAGATCGACGAGCTGGGAAAGGAGATCCGGCCTTGTCGCCTGAAGAAGCGGCCTCCTGTGCGG**GGCGGCTCAGGCCCGCCCAAAAAAAAAAGAAAGGTG**TGA**ACCGGT**CAT |
| PsuCas9 | TCT**GGATCC**ATGAAGTACGCTATCGGCTTGGACATCGGCATTGCCAGCGTGGGATATGCGGTGCTGGCCCTGGATCACGAAGAGAACCCTTGGGGCATCATCAGACTGGGCAGCAGGATCTTTGACGTGGCAGAGAACCCAAAGGACGGTGCCTCTCTGGCCCTGCCAAGAAGAGAAGCGAGAAGCGTCAGACGGAGACTGCGGCGGCACCACCATAGACTGGAAAGGATCAAGAACCTGCTGATCAACACCGAGCTGATCACAAAGGACGAGCTGCTGCACCTGTACGACGGCAAGCTGAGCGATGTGTACGAGCTGAGAGTGAAAGCCCTGGACTACAGCGTGACCAACGCCGAGCTGACCAGAATCTTACTTCACCTGGCTCAGAGACGTGGATTTAAGAGCAACAGAAAGTCAGATGCCGGAGACAAGGAAGCCGGCCAGCTGCTGGAAGCCGTGTCTGCTAATGCCAAAAGAATGCAAGAGAACCACTACAGAACAGTCGGCGAGATGTTCTATAAAGATGACCTGTTCAGCAAGTACAAAAGGAACAAGGGGGGAACCTACCTGACAACGGTGCACCGGGATATGATCGCTGCTGAGGCACGCACAATCCTCGAAAAACAGTTCGCCCTGGGCAACGACATCTGTACACAGGACTTCATCGACAAGTACCTGAGCATACTGCTGTCACAGCGGCAGTTCGATGAAGGCCCTGGCGAGCCTAGCCCTTACGCCGGCAACCAGATCGCCAATATGATCGGAAAATGCACCTTCGAGCCCAACGAGTACCGGGCAGCTAAGGCCTCCTACAGCTTCGAGAGATTCAACCTGCTGCAGAAAGTGAATCACCTGAGACTGCTCCTTGAAGGCAAGTCTATTGCTCTGGATAACGAGCAAAGAAAAAAGATCATCGCCCTGGCCCACGACAAGGCCGACCTGCGGTACTCCCACATCAGAAAGGCTCTGGGCCTGGACGAAAAGGTGCTCTTTAACACCATCACCTACAACGACGATGTTTCTGTGATCGAGAAGAAAACCAAGTTCAATTTCCTGCAGGCCTACCACCAGATCAAGAAGGTCCTGGCTGAGGATATGCAGAAACTCACAACCGAGCAGCTGGATAACATCGGCCAGATTCTGTCCACATACAAGAGCGACAACAAGAGAACAGAAGAACTGTCTGCCCTGGGCCTGGAGAAGAAGATCATCGACGCCCTGCTGGGCATCAACGGCCTGTCCAAGTTCGCTCATCTGAGCCTGAAGGCCCTCAGAAAAATCAACCCTTACCTGGAAGAGGGCAAGATATACAACGAGGCCTGTGCCGCTGCCGGATACGACTTCAAGGCCCACGCCAATACCCAGAAAACCGAGCTGCTGCCTGCTTATAAGGAAGAGATGGACGACATCACCAGCCCCGTGGCCAGACGGGCCATCGCCCAGTCCATCAAGGTGATCAACGCCATCATCAGAGAACAAAAATGCAGCCCCGTGTACATCAACATCGAGCTGGCCAGAGAGATGGCCAAGGGCTTTGACGAGCGGACCCAAATCGACAAGGCCAATAAGGAAAACCAGGCCAAGAACGAGAGAATTATGGAGAGAATCAGAACCGAGTTCCACAAGAGCAATCCTACCGGAATGGACCTGATCAAACTGAAACTGTGGGAGGAGCAGGACGGCAGATCTCCTTATAGCCAGAAGGCCATCTCTATTAATCGGCTGTTTGAACCTGGATATGTGGATATTGACCATATCGTGCCTTATAGCATCAGCTTTGATGATAGCTTCAAAAACAAGGTGCTGGTGTTCTCTGATGAGAACCGGGACAAGGGCAATCGGCTGCCTATCGCCTACCTGCAGAGCAAGTTCGGCGCCAAGGCCGCTGAAAATTTCATCATCTGGGTCCAGAGCAACATCAAGGACTACAAGAAACGGCAAAAATTACTTAAGCGAGAAATCACCGAGGAGGACATCAACAAGTTCAAGGAGAGGAACCTGCAAGACACTAAGACAATCAGCCGCTTCCTGTACAACTACATTAATGACTACCTGCTGTTTGCCCCTAGCGATACCGGCAAGAAAAAGCGGGTGACCGCCGTGAACGGCACCATCACAGCCTACCTGAGAAAGAGGTGGGGCATCAATAAGATCCGGGCCAATGGCGATAAGCACCACGCCGTGGACGCTGTTGTGATCGCCTGCACCACAGACAAGATGATCAAGGACCTGAGCAGCTTCAGCCAGTACCACGAGCTGGAATACACACACACAGACACAGAGAGCCTGCTCGTGAACAGCCTGACCGGCGAAATCCTGAAGCGGTTCCCCTATCCTTGGGAAGATTTTAGACCTGAGCTGATGGCCAGACTGTCTGACAACCCAGCTGACGCCCTGAGAAAGCTCAACCTGATCTTCTACCACGGCACCGACCTGAGCACCATCAAACCAATCTTCGTGTCCAGAATGCCCAGACACAAAGTGACCGGCGCCGCTCATAAGGCTACAATCAAGTCTGCCCGGTGCCTGAACAACGGCATCGTGATCTGCAAGACCTCCCTGCAGAACCTGAAGCTGGATAAGGAAGGCGAGATCGCGAACTACTACGCCCCTGAGTCCGACACAATTCTGTACAACGCCCTTAAGGAACGCCTGAGAGCCTATGATAATAAACCCGCTAAGGCCTTCGCCGAGCCTTTCTACAAGCCCAAGGCTGACGGCACCCCTGGACCTCTGGTGAAGAAAGTGAAAGTGTACGAGAAATCCACCCTGAACGTGGCCGTGCAGCAGAACACCGCCGTGGCCGACAACGACTCTATGGTGCGGGTCGACGTGTTCCACGTGAAGGGCGACGGATACTACCTGGTGCCCATCTACATAGCCGACACCCTGAAGAAGGAGCTGCCCAACAAGGCCATCGTGGCCTACAAGCCTTACGCCGATTGGCCTGTGATGGACGATAGCAACTTCATCTTCAGCCTGTACCCTAATGACCTCATCAAGTTCGAGCACAGAAACGGCGTGAAGTTTGCCAAAGTGAATAAAGAGAGCAGCCTGGCCGAAACCTACCTGACCAAGAGTGAACTGGTTTACTACAAGGGCACAAACATCACCGTGGGCAGTACAAGCATAATCACCAACGATAACAGCTACACCGTGAAGAGCCTGGGCGTGAAAACACTGTCTAATCTGGAAAAGTACCAGGTGGACGTGCTCGGAAACTATACCAAGGTGAAGAAGGAAATCAGACGGCCCTTCAGA**GGCGGCTCAGGCCCGCCCAAAAAAAAAAGAAAGGTG**TGA**ACCGGT**CAT |
| GfoCas9 | TCT**GGATCC**ATGAAGCACCCTTACGGAATCGGCCTGGATATCGGCATCGCCAGCGTGGGCTGGGCTGTTGTGGCCCTGAACGAGAACGCCGAGCCTTACGGCCTTATCAGGTGCGGCTCTAGAATCTTCGACAAGGCTGAGCAGCCTAAGACGGGCGACTCACTGGCCGCCCCTAGAAGAGAGGCCCGCAGCGCCAGAAGGCGGCTGCGGAGAAGAAGCCTGAGAAAGGCCGACCTCTACGAGCTGATGGAAAAGAACGGCCTGCCTGGTAAGGCCGAGATCGAGCAGGCCGTGCAGGCCGGACACCTACCCGACGTGTATGCCCTGAGAGTGCAAGCCCTGGATGGACCTGTGACAGCTCTGGATTTCGCCAGAATCCTGCTGCACCTGATGCAGAGAAGGGGCTTCAGAAGCAACCGGAAGGCTGATGACGCCCAGAAAGACGGCAAGCTGCTGCAGGCCATTGACGCCAACACCAGACGGATGGAAGCCAATCGGTACAGAACTGTGGGAGAGATGATGTACCGCGACCCCGTGTTTGCCGAGCACAAGAGAAACAAGGCTGAGAATTACCTGAGCACCGTGAAGCGGGATCAGATTATCGATGAGGCCAGACTGGTGTTCGCCGCCCAGCGACAGTATGGCGCTACCTGGGCCTCTCCTGAGATGGAAGCCGAGTACCTGTGCATCCTGACAAGACAGAGAAGCTTCGCCGAGGGCCCCGGAAAGGGCAGCCCTTATAGTGGCAGCAACCGGGTGGGAACCTGTACCCTGGAAGGCAAGAGCGAGCAGCGGGCCGCCAAGGCCGCTTTCAGCTTCGAGTACTTCACCCTGCTCCAGAAGATCAACCACATCAGAATCGCTGAGAACGGCACCTCCAGAACCCTGACACCTGCTGAAAGACAGGTGCTGCTGAGCGCCTGCTACCAAACCGACAAGCTGGACTTCGCCAGAATCAGAAAAGCTCTGGCCCTGCCTGAGGAAGCTCGGTTCAATATGGTGCGGTACCGGGGCGAGCAAACCGCCGAGGCCTGTGAAAAGAAGGAAAAAATCACCGCCCTGCCGTGCTACCACAAGATGCGCAAGGCTCTGAATACCCTGCGGAAGGATCACATCAGAAACATCAGCAGGGAGCAGCTGGACGCCGCTGGCGCTGCCCTGACCAATCCTGAAAACGAGGACAAGCTGAGAGAGGCCCTGAAACAGGCCGAGTTCGAGCCACTGGAAATCGAGGCTCTGCTGACGCTGCCTAGCTTCGCCGGCTACGGCCATATCTCCGTGAAAGCCTGCAGAAAGCTCATCCCTTACCTGGAACAAGGGATGAACTACAACGACGCCTGCCAGGCTGCCGGCTACGACTTTCAGGGAAGCCAGAATGGCGAGAAGGCACAATTTCTGCCTGCCTCTACAGAGGAAATGGAAGATATCACAAGCCCCGTGGTGCGAAGAGCCGTGGCCCAGACCATCAAAGTGGTGAACGCTATCATCCGGGAACAGGGCGAGTCTCCCGTGTCCATCCACCTGGAACTGGCCCGTGAAATGAACAAAAACTTCCAGCAGAGATCTGAGCTGGACAAAGCTATGAGAGACAACAGCGCCGAGAACGAGAGACTGATGAAGGAACTCAATGAGCTTTTTCCAGGACGGACAGTGACCGGACAGGATCTCGTGAAGTACAGACTGTGGAAGGAGCAGGACGGCAGATGCGCCTACAGCATCCAACCTCTGGAACTGGACAAGGTGATCACCGTGTCTGGCTACGCCGAGGTGGACCACATCGTGCCTTACAGCATCTCCTTCGATGATAGAAGAACCAACAAGGTGCTGGTGCTGGCTTCTGAAAATAGACAAAAGGGCAATAGACTGCCACTGCAGTACCTGCAAGGCAAGCGGAGAGATGACTTCATCGTGTACACCAAGGCCAACGTGAAAAACTTCAGAAAGCGGCAGAACCTGCTGAAGGAAAGACTGAGCGAAGAGGACGGCAAAGGCTACATCCAGCGGAACCTGCAGGACACCCAGTACATTGCCGCCTTTATGCTGAACTACATTAGAAATCACCTGGCCTTCGCCGACTGCAGCGGCGCTGGCAAAAGACGGGTGGTGGCTGTCAACGGCGCCGTGACCGCCTTCCTGAGAAAGAGATGGGGCCTGAGCAAGGTTAGAGCCGATGGGGATCTGCACCACGCCGCTGATGCCGCCGTGATCGCTTGCACAACCCAGGGAATGATCAAGAGAGTCAGCGACTTCTGCAAGAGAGCCGAAACCACAGCCGTGCGGAACGAGCATTTCCCTGAGCCTTGGCCTAGATTTCGGGACGAGCTGACCCAGAGACTGAGCGCCTGTCCTCAGGAGAACCTGATGCAGATCAACCCCGTTTACTACGCCACAGTGGACATCTCCTCTATCCAGCCCGTGTTCGCGTCCAGAATGCCACGGCACAAGGTGACCGGCGCCGCCCACAAAGAGACAATCAAGAGCCGGCTGGACGATACCCACGTGGTGCAGAGAAGAAACATCACCGAGCTGAAGCTGGACAAGGACGGCGAAATCGCTGGATACTTCAACAGAAGTTCCGATACACTGCTGTACAACGCCCTTAAGGCAAGACTGCTGGCTTTTGGCGGCGACGGCAAGAAAGCCTTCGCTGAACCCTTCTACAAGCCCAGGGCCGACGGCACACCTGGCGCACGGGTGCAAAAGGTGAAGATTTGCGACAAAGTGACATCTACCGTGCCCGTGCACGGCGGCAAGGGCGTGGCCGACAATGACACAATGGTGCGAATGGACGTGTACTACGTGCCTGGCGACGGCTACTATTGGATTCCTGTGTACGTTGCCGACACCGTGAAACCTGAGCTGCCCAGCAAGGCCGTGGTGGCTTACAAGTCCAGCGCTGAATGGAAGGAAATGAAAGACGAGGATTTTCTGTTCAGCCTGTGTCAGCACGACCTGGTCAGAATCGAGAGCAAGAGACTGATGAAATTCAAGGTGCAGAACCGCGATAGCACCCTGGAAAAAGAGATGCCTGTGAACAAGATCCTGGCCTATTTCGAGGGCGGGAATATCAGCACCGGCGCCATAACAGTGACCACCCACGATGACGCCTACATCGCTGATGGACTGGGATTTAAGACCCTGCAGAAAGTCCAGAAGTACCAGGTGGATGTGCTGGGCAACTACACACCCGTCAAGAAGGAAAAGAGACAGACATTCCCTGCTCAGAGGCGG**GGCGGCTCAGGCCCGCCCAAAAAAAAAAGAAAGGTG**TGA**ACCGGT**CAT |

Green highlighted: short GS linker

Red highlighted: SV40 NLS

Yellow highlighted: BamHI restriction site

Pink highlighted: AgeI restriction site

**Table S10.** **Complementary oligonucleotide pairs used for guide RNA cloning into BbsI-digested sgRNA backbone vectors**

| **Oligonucleotide Name** | **Sequence (5’---3’)** |
| --- | --- |
| Nme2Cas9/NsuCas9-*Casp-3* target-1-N_4_CC_Top | CACCGAACTGCCTAGTTATGGATGAAT |
| Nme2Cas9/NsuCas9-*Casp-3* target-1-N_4_CC_Bot | CAACATTCATCCATAACTAGGCAGTTC |
| Nme2Cas9/NsuCas9-*HBB* target-1-N_4_CC_Top | CACCGGGTGAGCCAGGCCATCACTAAA |
| Nme2Cas9/NsuCas9-*HBB* target-1-N_4_CC_Bot | CAACTTTAGTGATGGCCTGGCTCACCC |
| Nme2Cas9/NsuCas9-*HBB* target-2-N_4_CC_Top | CACCGCTCAGAATAATCCAGCCTTATC |
| Nme2Cas9/NsuCas9-*HBB* target-2-N_4_CC_Bot | CAACGATAAGGCTGGATTATTCTGAGC |
| PsuCas9_*Casp-3* target-1-N_4_ATAA_Top | CACCGCCATCTGCTAGTTATTGGAAGC |
| PsuCas9_*Casp-3* target-1-N_4_ATAA_Bot | CAACGCTTCCAATAACTAGCAGATGGC |
| PsuCas9_*Casp-3* target-1-N_4_GAAA_Top | CACCGTTCCAATAACTAGCAGATGGAA |
| PsuCas9_*Casp-3* target-1-N_4_GAAA_Bot | CAACTTCCATCTGCTAGTTATTGGAAC |
| PsuCas9_*Casp-3* target-1-N_4_ACAA_Top | CACCGATGCAATGCCAGTTTTCAGTCC |
| PsuCas9_*Casp-3* target-1-N_4_ACAA_Bot | CAACGGACTGAAAACTGGCATTGCATC |
| PsuCas9_*HBB* target-1-N_4_ATAA_Top | CACCGCATGAGCCTTCACCTTAGGGTT |
| PsuCas9_*HBB* target-1-N_4_ATAA_Bot | CAACAACCCTAAGGTGAAGGCTCATGC |
| PsuCas9_*HBB* target-1-N_4_GAAA_Top | CACCGAACCCTAAGGTGAAGGCTCATG |
| PsuCas9_*HBB* target-1-N_4_GAAA_Bot | CAACCATGAGCCTTCACCTTAGGGTTC |
| PsuCas9_*HBB* target-1-N_4_ACAA_Top | CACCGTGCCACACTGAGTGAGCTGCAC |
| PsuCas9_*HBB* target-1-N_4_ACAA_Bot | CAACGTGCAGCTCACTCAGTGTGGCAC |
| PsuCas9_*HBB* target-2-N_4_ATAA_Top | CACCGCAGAATAATCCAGCCTTATCCC |
| PsuCas9_*HBB* target-2-N_4_ATAA_Bot | CAACGGGATAAGGCTGGATTATTCTGC |
| PsuCas9_*HBB* target-2-N_4_GAAA_Top | CACCGTGGATTGTAGCTGCTATTAGCA |
| PsuCas9_*HBB* target-2-N_4_GAAA_Bot | CAACTGCTAATAGCAGCTACAATCCAC |
| PsuCas9_*HBB* target-2-N_4_ACAA_Top | CACCGGAGGTTTCATATTGCTAATAGC |
| PsuCas9_*HBB* target-2-N_4_ACAA_Bot | CAACGCTATTAGCAATATGAAACCTCC |
| GfoCas9_*Casp-3* target-1-N_4_ATAA_Top | CACCGCCATCTGCTAGTTATTGGAAGC |
| GfoCas9_*Casp-3* target-1-N_4_ATAA_Bot | TGACGCTTCCAATAACTAGCAGATGGC |
| GfoCas9_*Casp-3* target-1-N_4_GAAA_Top | CACCGTTCCAATAACTAGCAGATGGAA |
| GfoCas9_*Casp-3* target-1-N_4_GAAA_Bot | TGACTTCCATCTGCTAGTTATTGGAAC |
| GfoCas9_*Casp-3* target-1-N_4_ACAA_Top | CACCGATGCAATGCCAGTTTTCAGTCC |
| GfoCas9_*Casp-3* target-1-N_4_ACAA_Bot | TGACGGACTGAAAACTGGCATTGCATC |
| GfoCas9_*HBB* target-1-N_4_ATAA_Top | CACCGCATGAGCCTTCACCTTAGGGTT |
| GfoCas9_*HBB* target-1-N_4_ATAA_Bot | TGACAACCCTAAGGTGAAGGCTCATGC |
| GfoCas9_*HBB* target-1-N_4_GAAA_Top | CACCGAACCCTAAGGTGAAGGCTCATG |
| GfoCas9_*HBB* target-1-N_4_GAAA_Bot | TGACCATGAGCCTTCACCTTAGGGTTC |
| GfoCas9_*HBB* target-1-N_4_ACAA_Top | CACCGTGCCACACTGAGTGAGCTGCAC |
| GfoCas9_*HBB* target-1-N_4_ACAA_Bot | TGACGTGCAGCTCACTCAGTGTGGCAC |
| GfoCas9_*HBB* target-2-N_4_ATAA_Top | CACCGCAGAATAATCCAGCCTTATCCC |
| GfoCas9_*HBB* target-2-N_4_ATAA_Bot | TGACGGGATAAGGCTGGATTATTCTGC |
| GfoCas9_*HBB* target-2-N_4_GAAA_Top | CACCGTGGATTGTAGCTGCTATTAGCA |
| GfoCas9_*HBB* target-2-N_4_GAAA_Bot | TGACTGCTAATAGCAGCTACAATCCAC |
| GfoCas9_*HBB* target-2-N_4_ACAA_Top | CACCGGAGGTTTCATATTGCTAATAGC |
| GfoCas9_*HBB* target-2-N_4_ACAA_Bot | TGACGCTATTAGCAATATGAAACCTCC |
| SpCas9-*Casp-3* target-1-NGG_Top | CACCGGCAATGCCAGTTTTCAGTCC |
| SpCas9-*Casp-3* target-1-NGG_Bot | AAACGGACTGAAAACTGGCATTGCC |
| SpCas9-*HBB* target-1-NGG_Top | CACCGTTGCCATGAGCCTTCACCTT |
| SpCas9-*HBB* target-1-NGG_Bot | AAACAAGGTGAAGGCTCATGGCAAC |
| SpCas9-*HBB* target-2-NGG_Top | CACCGGATTATTCTGAGTCCAAGCT |
| SpCas9-*HBB* target-2-NGG_Bot | AAACAGCTTGGACTCAGAATAATCC |
| Nme2Cas9/NsuCas9_*AIFM* target-1_N_4_CC_Top | CACCGGTTGTCTCTCACATCCAGCTGT |
| Nme2Cas9/NsuCas9_*AIFM* target-1_N_4_CC_Bot | CAACACAGCTGGATGTGAGAGACAACC |
| Nme2Cas9/NsuCas9_*CLTA* target-1_N_4_CC_Top | CACCGTGGCAGATGAAGCTTTCTACAA |
| Nme2Cas9/NsuCas9_*CLTA* target-1_N_4_CC_Bot | CAACTTGTAGAAAGCTTCATCTGCCAC |
| Nme2Cas9/NsuCas9_*CLTA* target-2_N_4_CC_Top | CACCGCTGTTTCTCTTCCAATCTGACG |
| Nme2Cas9/NsuCas9_*CLTA* target-2_N_4_CC_Bot | CAACCGTCAGATTGGAAGAGAAACAGC |
| PsuCas9_*AIFM* target-1_N_4_ATAA_Top | CACCGTGGTGAAACTTAATGATGGCTC |
| PsuCas9_*AIFM* target-1_N_4_ATAA_Bot | CAACGAGCCATCATTAAGTTTCACCAC |
| PsuCas9_*AIFM* target-1_N_4_GAAA_Top | CACCGCAGCTGGATGTGAGAGACAACA |
| PsuCas9_*AIFM* target-1_N_4_GAAA_Bot | CAACTGTTGTCTCTCACATCCAGCTGC |
| PsuCas9_*AIFM* target-1_N_4_ACAA_Top | CACCGCCAGGTAGTACAGCTGGATGTG |
| PsuCas9_*AIFM* target-1_N_4_ACAA_Bot | CAACCACATCCAGCTGTACTACCTGGC |
| PsuCas9_*CLTA* target-1_N_4_ATAA_Top | CACCGTACAGCCAGCATGGGCAACCAA |
| PsuCas9_*CLTA* target-1_N_4_ATAA_Bot | CAACTTGGTTGCCCATGCTGGCTGTAC |
| PsuCas9_*CLTA* target-1_N_4_GAAA_Top | CACCGATCACGTCAGCGAAGGGTTGTT |
| PsuCas9_*CLTA* target-1_N_4_GAAA_Bot | CAACAACAACCCTTCGCTGACGTGATC |
| PsuCas9_*CLTA* target-1_N_4_ACAA_Top | CACCGAGGGTGGCAGATGAAGCTTTCT |
| PsuCas9_*CLTA* target-1_N_4_ACAA_Bot | CAACAGAAAGCTTCATCTGCCACCCTC |
| PsuCas9_*CLTA* target-2_N_4_ATAA_Top | CACCGGCTCAGTACTGATATATGCATA |
| PsuCas9_*CLTA* target-2_N_4_ATAA_Bot | CAACTATGCATATATCAGTACTGAGCC |
| PsuCas9_*CLTA* target-2_N_4_GAAA_Top | CACCGACCTGGGAACACGTCAGATTGG |
| PsuCas9_*CLTA* target-2_N_4_GAAA_Bot | CAACCCAATCTGACGTGTTCCCAGGTC |
| PsuCas9_*CLTA* target-2_N_4_ACAA_Top | CACCGCAGTACTGATATATGCATACCT |
| PsuCas9_*CLTA* target-2_N_4_ACAA_Bot | CAACAGGTATGCATATATCAGTACTGC |
| GfoCas9_*AIFM* target-1_N_4_ATAA_Top | CACCGTGGTGAAACTTAATGATGGCTC |
| GfoCas9_*AIFM* target-1_N_4_ATAA_Bot | TGACGAGCCATCATTAAGTTTCACCAC |
| GfoCas9_*AIFM* target-1_N_4_GAAA_Top | CACCGCAGCTGGATGTGAGAGACAACA |
| GfoCas9_*AIFM* target-1_N_4_GAAA_Bot | TGACTGTTGTCTCTCACATCCAGCTGC |
| GfoCas9_*AIFM* target-1_N_4_ACAA_Top | CACCGCCAGGTAGTACAGCTGGATGTG |
| GfoCas9_*AIFM* target-1_N_4_ACAA_Bot | TGACCACATCCAGCTGTACTACCTGGC |
| GfoCas9_*CLTA* target-1_N_4_ATAA_Top | CACCGTACAGCCAGCATGGGCAACCAA |
| GfoCas9_*CLTA* target-1_N_4_ATAA_Bot | TGACTTGGTTGCCCATGCTGGCTGTAC |
| GfoCas9_*CLTA* target-1_N_4_GAAA_Top | CACCGATCACGTCAGCGAAGGGTTGTT |
| GfoCas9_*CLTA* target-1_N_4_GAAA_Bot | TGACAACAACCCTTCGCTGACGTGATC |
| GfoCas9_*CLTA* target-1_N_4_ACAA_Top | CACCGAGGGTGGCAGATGAAGCTTTCT |
| GfoCas9_*CLTA* target-1_N_4_ACAA_Bot | TGACAGAAAGCTTCATCTGCCACCCTC |
| GfoCas9_*CLTA* target-2_N_4_ATAA_Top | CACCGGCTCAGTACTGATATATGCATA |
| GfoCas9_*CLTA* target-2_N_4_ATAA_Bot | TGACTATGCATATATCAGTACTGAGCC |
| GfoCas9_*CLTA* target-2_N_4_GAAA_Top | CACCGACCTGGGAACACGTCAGATTGG |
| GfoCas9_*CLTA* target-2_N_4_GAAA_Bot | TGACCCAATCTGACGTGTTCCCAGGTC |
| GfoCas9_*CLTA* target-2_N_4_ACAA_Top | CACCGCAGTACTGATATATGCATACCT |
| GfoCas9_*CLTA* target-2_N_4_ACAA_Bot | TGACAGGTATGCATATATCAGTACTGC |
| SpCas9_*AIFM* target-1_NGG_Top | CACCGGCTGGATGTGAGAGACAACA |
| SpCas9_*AIFM* target-1_NGG_Bot | AAACTGTTGTCTCTCACATCCAGCC |
| SpCas9_*CLTA* target-1_NGG_Top | CACCGTAACCAATCACGTCAGCGAA |
| SpCas9_*CLTA* target-1_NGG_Bot | AAACTTCGCTGACGTGATTGGTTAC |
| SpCas9_*CLTA* target-2_NGG_Top | CACCGTTCCAATCTGACGTGTTCCC |
| SpCas9_*CLTA* target-2_NGG_Bot | AAACGGGAACACGTCAGATTGGAAC |
| NsuCas9/Nme2Cas9_*CLTA* target-3_N_4_CC_Top | CACCGCAGTGTTCTTCCCACCAGACCA |
| NsuCas9/Nme2Cas9_*CLTA* target-3_N_4_CC_Bot | CAACTGGTCTGGTGGGAAGAACACTGC |
| PsuCas9_*CLTA* target-3_N_4_ATAA_Top | CACCGGGGAAGCAGTAGTTAGATAATC |
| PsuCas9_*CLTA* target-3_N_4_ATAA_Bot | CAACGATTATCTAACTACTGCTTCCCC |
| PsuCas9_*CLTA* target-3_N_4_ACAA_Top | CACCGGAGTAGAGAGAGAGCCATATTC |
| PsuCas9_*CLTA* target-3_N_4_ACAA_Bot | CAACGAATATGGCTCTCTCTCTACTCC |
| GfoCas9_*CLTA* target-1_N_4_GAAA_Top | CACCGAGGAGCATTAACTCTAAACAAA |
| GfoCas9_*CLTA* target-1_N_4_GAAA_Bot | TGACTTTGTTTAGAGTTAATGCTCCTC |
| NsuCas9/Nme2Cas9_*CLTA* target-4_N_4_CC_Top | CACCGCTAGGAGACAGGACATCAGAAGA |
| NsuCas9/Nme2Cas9_*CLTA* target-4_N_4_CC_Bot | CAACTCTTCTGATGTCCTGTCTCCTAGC |
| PsuCas9_*CLTA* target-2_N_4_ATAA-3_Top | CACCGTAGGCTTTATACATACGTGAGT |
| PsuCas9_*CLTA* target-2_N_4_ATAA-3_Bot | CAACACTCACGTATGTATAAAGCCTAC |
| PsuCas9_*CLTA* target-4_N_4_ACAA_Top | CACCGTTTCTGACATGCAAGACCACAG |
| PsuCas9_*CLTA* target-4_N_4_ACAA_Bot | CAACCTGTGGTCTTGCATGTCAGAAAC |
| GfoCas9_*CLTA* target-4_N_4_GAAA_Top | CACCGTTGTTCCTCTGTGGTCTTGCAT |
| GfoCas9_*CLTA* target-4_N_4_GAAA_Bot | TGACATGCAAGACCACAGAGGAACAAC |
| NsuCas9/Nme2Cas9_*AIFM* target-2_N_4_CC_Top | CACCGGGTGTTATCTAAGTTGGTTCAC |
| NsuCas9/Nme2Cas9_*AIFM* target-2_N_4_CC_Bot | CAACGTGAACCAACTTAGATAACACCC |
| PsuCas9_*AIFM* target-2_N_4_ATAA_Top | CACCGTTTAAGCTAGCAACCAAGGAAC |
| PsuCas9_*AIFM* target-2_N_4_ATAA_Bot | CAACGTTCCTTGGTTGCTAGCTTAAAC |
| PsuCas9_*AIFM* target-2_N_4_ACAA_Top | CACCGCCTAAACTTGTAATGGCTTAAG |
| PsuCas9_*AIFM* target-2_N_4_ACAA_Bot | CAACCTTAAGCCATTACAAGTTTAGGC |
| GfoCas9_*AIFM* target-2_N_4_GAAA_Top | CACCGCTTTGTTCCTTGGTTGCTAGCT |
| GfoCas9_*AIFM* target-2_N_4_GAAA_Bot | TGACAGCTAGCAACCAAGGAACAAAGC |
| NsuCas9/Nme2Cas9_*HBB* target-3_N_4_CC_Top | CACCGGGCCCATCACTTTGGCAAAGAA |
| NsuCas9/Nme2Cas9_*HBB* target-3_N_4_CC_Bot | CAACTTCTTTGCCAAAGTGATGGGCCC |
| PsuCas9_*HBB* target-3_N_4_ATAA_Top | CACCGCAGAATCCAGATGCTCAAGGCC |
| PsuCas9_*HBB* target-3_N_4_ATAA_Bot | CAACGGCCTTGAGCATCTGGATTCTGC |
| PsuCas9_*HBB* target-3_N_4_ACAA_Top | CACCGCCCAGTTTAGTAGTTGGACTTA |
| PsuCas9_*HBB* target-3_N_4_ACAA_Bot | CAACTAAGTCCAACTACTAAACTGGGC |
| GfoCas9_*HBB* target-3_N_4_GAAA_Top | CACCGCTTAGGGAACAAAGGAACCTTT |
| GfoCas9_*HBB* target-3_N_4_GAAA_Bot | TGACAAAGGTTCCTTTGTTCCCTAAGC |
| NsuCas9/Nme2Cas9_*HBB* target-4_N_4_CC_Top | CACCGGATACAATGTATCATGCCTCTT |
| NsuCas9/Nme2Cas9_*HBB* target-4_N_4_CC_Bot | CAACAAGAGGCATGATACATTGTATCC |
| PsuCas9_*HBB* target-2_N_4_ATAA_Top | CACCGTTGCACCATTCTAAAGAATAAC |
| PsuCas9_*HBB* target-2_N_4_ATAA_Bot | CAACGTTATTCTTTAGAATGGTGCAAC |
| PsuCas9_*HBB* target-4_N_4_ACAA_Top | CACCGTAGGGAAAGTATTAGAAATAAG |
| PsuCas9_*HBB* target-4_N_4_ACAA_Bot | CAACCTTATTTCTAATACTTTCCCTAC |
| GfoCas9_*HBB* target-4_N_4_GAAA_Top | CACCGTACATTGTATCATTATTGCCCT |
| GfoCas9_*HBB* target-4_N_4_GAAA_Bot | TGACAGGGCAATAATGATACAATGTAC |
| NsuCas9/Nme2Cas9_*EMX* target-1_N_4_CC_Top | CACCGAGCTAGGATGCACAGCAGCTCT |
| NsuCas9/Nme2Cas9_*EMX* target-1_N_4_CC_Bot | CAACAGAGCTGCTGTGCATCCTAGCTC |
| PsuCas9_*EMX* target-1_N_4_ATAA_Top | CACCGCTCTACGAGTTTCTAGAGGAGA |
| PsuCas9_*EMX* target-1_N_4_ATAA_Bot | CAACTCTCCTCTAGAAACTCGTAGAGC |
| PsuCas9_*EMX* target-1_N_4_ACAA_Top | CACCGAAGGGTGGTTTTCCTGTTCCTC |
| PsuCas9_*EMX* target-1_N_4_ACAA_Bot | CAACGAGGAACAGGAAAACCACCCTTC |
| GfoCas9_*EMX* target-1_N_4_GAAA_Top | CACCGAAGAAGCGATTATGATCTCTCC |
| GfoCas9_*EMX* target-1_N_4_GAAA_Bot | TGACGGAGAGATCATAATCGCTTCTTC |
| SpCas9_*EMX* target-1_NGG_Top | CACCGTCGTAGAGTCCCATGTCTGC |
| SpCas9_*EMX* target-1_NGG_Bot | AAACGCAGACATGGGACTCTACGAC |
| NsuCas9/Nme2Cas9_Casp3 target-2_N_4_CC_Top | CACCGGAATTGTGGAATTATACTGTAC |
| NsuCas9/Nme2Cas9_Casp3 target-2_N_4_CC_Bot | CAACGTACAGTATAATTCCACAATTCC |
| PsuCas9_Casp3 target-2_N_4_ATAA_Top | CACCGACTATGGTCTAGCAATAATTAT |
| PsuCas9_Casp3 target-2_N_4_ATAA_Bot | CAACATAATTATTGCTAGACCATAGTC |
| PsuCas9_Casp3 target-2_N_4_ACAA_Top | CACCGGAATTGATGCGTGATGGTAAGA |
| PsuCas9_Casp3 target-2_N_4_ACAA_Bot | CAACTCTTACCATCACGCATCAATTCC |
| GfoCas9_Casp3 target-2_N_4_GAAA_Top | CACCGGTGGAATTGATGCGTGATGGTA |
| GfoCas9_Casp3 target-2_N_4_GAAA_Bot | TGACTACCATCACGCATCAATTCCACC |
| SpCas9_*CLTA* target-3_NGG_Top | CACCGAGAACACTGGACTAGGATCC |
| SpCas9_*CLTA* target-3_NGG_Bot | AAACGGATCCTAGTCCAGTGTTCTC |
| SpCas9_*CLTA* target-4_NGG_Top | CACCGTCTGACATGCAAGACCACAG |
| SpCas9_*CLTA* target-4_NGG_Bot | AAACCTGTGGTCTTGCATGTCAGAC |
| SpCas9_*HBB* target-3_NGG_Top | CACCGATATCCCCCAGTTTAGTAGT |
| SpCas9_*HBB* target-3_NGG_Bot | AAACACTACTAAACTGGGGGATATC |
| SpCas9_*HBB* target-4_NGG_Top | CACCGTTCTTTAGAATGGTGCAAAG |
| SpCas9_*HBB* target-4_NGG_Bot | AAACCTTTGCACCATTCTAAAGAAC |
| SpCas9_Casp3 target-2_NGG_Top | CACCGTCTTACACGTGAAGAAATTG |
| SpCas9_Casp3 target-2_NGG_Bot | AAACCAATTTCTTCACGTGTAAGAC |
| SpCas9_*AIFM* target-2_NGG_Top | CACCGCTAGTAACACCCTTGATTGA |
| SpCas9_*AIFM* target-2_NGG_Bot | AAACTCAATCAAGGGTGTTACTAGC |

**Table S11. Backbone plasmid sequences for cloning guides**

| **Name** | **Sequence (5’---3’)** |
| --- | --- |
| NsuCas9 sgRNA backbone | ATTGTTGCCGGGAAGCTAGAGTAAGTAGTTCGCCAGTTAATAGTTTGCGCAACGTTGTTGCCATTGCTACAGGCATCGTGGTGTCACGCTCGTCGTTTGGTATGGCTTCATTCAGCTCCGGTTCCCAACGATCAAGGCGAGTTACATGATCCCCCATGTTGTGCAAAAAAGCGGTTAGCTCCTTCGGTCCTCCGATCGTTGTCAGAAGTAAGTTGGCCGCAGTGTTATCACTCATGGTTATGGCAGCACTGCATAATTCTCTTACTGTCATGCCATCCGTAAGATGCTTTTCTGTGACTGGTGAGTACTCAACCAAGTCATTCTGAGAATAGTGTATGCGGCGACCGAGTTGCTCTTGCCCGGCGTCAATACGGGATAATACCGCGCCACATAGCAGAACTTTAAAAGTGCTCATCATTGGAAAACGTTCTTCGGGGCGAAAACTCTCAAGGATCTTACCGCTGTTGAGATCCAGTTCGATGTAACCCACTCGTGCACCCAACTGATCTTCAGCATCTTTTACTTTCACCAGCGTTTCTGGGTGAGCAAAAACAGGAAGGCAAAATGCCGCAAAAAAGGGAATAAGGGCGACACGGAAATGTTGAATACTCATACTCTTCCTTTTTCAATATTATTGAAGCATTTATCAGGGTTATTGTCTCATGAGCGGATACATATTTGAATGTATTTAGAAAAATAAACAAATAGGGGTTCCGCGCACATTTCCCCGAAAAGTGCCACCTGACGTCGCTAGCTGTACAAAAAAGCAGGCTTTAAAGGAACCAATTCAGTCGACTGGATCCGGTACCAAGGTCGGGCAGGAAGAGGGCCTATTTCCCATGATTCCTTCATATTTGCATATACGATACAAGGCTGTTAGAGAGATAATTGGAATTAATTTGACTGTAAACACAAAGATATTAGTACAAAATACGTGACGTAGAAAGTAATAATTTCTTGGGTAGTTTGCAGTTTTAAAATTATGTTTTAAAATGGACTATCATATGCTTACCGTAACTTGAAAGTATTTCGATTTCTTGGCTTTATATATCTTGTGGAAAGGACGAAACACCGGGTCTTCGAGAAGACCTGTTGTAGCTCCCTTTCTCATTTCGGAAACGAAATGAGAAACGTTGCTACAATAAGGCCGTCTGAAAAGATGTGCCGCAACGCTCTGCCCCTTAAGGCTTCTGCTTTAAGGGGCATCGTTTTTGGCCGGCATGGTCCCAGCCTCCTCGCTGGCGCCGGCTGGGCAACATGCTTCGGCATGGCGAATGGGACTTTTTTTTAAGCTTGGGCCGCTCGAGGTACCTCTCTACATATGACATGTGAGCAAAAGGCCAGCAAAAGGCCAGGAACCGTAAAAAGGCCGCGTTGCTGGCGTTTTTCCATAGGCTCCGCCCCCCTGACGAGCATCACAAAAATCGACGCTCAAGTCAGAGGTGGCGAAACCCGACAGGACTATAAAGATACCAGGCGTTTCCCCCTGGAAGCTCCCTCGTGCGCTCTCCTGTTCCGACCCTGCCGCTTACCGGATACCTGTCCGCCTTTCTCCCTTCGGGAAGCGTGGCGCTTTCTCATAGCTCACGCTGTAGGTATCTCAGTTCGGTGTAGGTCGTTCGCTCCAAGCTGGGCTGTGTGCACGAACCCCCCGTTCAGCCCGACCGCTGCGCCTTATCCGGTAACTATCGTCTTGAGTCCAACCCGGTAAGACACGACTTATCGCCACTGGCAGCAGCCACTGGTAACAGGATTAGCAGAGCGAGGTATGTAGGCGGTGCTACAGAGTTCTTGAAGTGGTGGCCTAACTACGGCTACACTAGAAGAACAGTATTTGGTATCTGCGCTCTGCTGAAGCCAGTTACCTTCGGAAAAAGAGTTGGTAGCTCTTGATCCGGCAAACAAACCACCGCTGGTAGCGGTGGTTTTTTTGTTTGCAAGCAGCAGATTACGCGCAGAAAAAAAGGATCTCAAGAAGATCCTTTGATCTTTTCTACGGGGTCTGACGCTCAGTGGAACGAAAACTCACGTTAAGGGATTTTGGTCATGAGATTATCAAAAAGGATCTTCACCTAGATCCTTTTAAATTAAAAATGAAGTTTTAAATCAATCTAAAGTATATATGAGTAAACTTGGTCTGACAGTTACCAATGCTTAATCAGTGAGGCACCTATCTCAGCGATCTGTCTATTTCGTTCATCCATAGTTGCCTGACTCCCCGTCGTGTAGATAACTACGATACGGGAGGGCTTACCATCTGGCCCCAGTGCTGCAATGATACCGCGAGACCCACGCTCACCGGCTCCAGATTTATCAGCAATAAACCAGCCAGCCGGAAGGGCCGAGCGCAGAAGTGGTCCTGCAACTTTATCCGCCTCCATCCAGTCTATTA |
| PsuCas9 sgRNA backbone | ATTGTTGCCGGGAAGCTAGAGTAAGTAGTTCGCCAGTTAATAGTTTGCGCAACGTTGTTGCCATTGCTACAGGCATCGTGGTGTCACGCTCGTCGTTTGGTATGGCTTCATTCAGCTCCGGTTCCCAACGATCAAGGCGAGTTACATGATCCCCCATGTTGTGCAAAAAAGCGGTTAGCTCCTTCGGTCCTCCGATCGTTGTCAGAAGTAAGTTGGCCGCAGTGTTATCACTCATGGTTATGGCAGCACTGCATAATTCTCTTACTGTCATGCCATCCGTAAGATGCTTTTCTGTGACTGGTGAGTACTCAACCAAGTCATTCTGAGAATAGTGTATGCGGCGACCGAGTTGCTCTTGCCCGGCGTCAATACGGGATAATACCGCGCCACATAGCAGAACTTTAAAAGTGCTCATCATTGGAAAACGTTCTTCGGGGCGAAAACTCTCAAGGATCTTACCGCTGTTGAGATCCAGTTCGATGTAACCCACTCGTGCACCCAACTGATCTTCAGCATCTTTTACTTTCACCAGCGTTTCTGGGTGAGCAAAAACAGGAAGGCAAAATGCCGCAAAAAAGGGAATAAGGGCGACACGGAAATGTTGAATACTCATACTCTTCCTTTTTCAATATTATTGAAGCATTTATCAGGGTTATTGTCTCATGAGCGGATACATATTTGAATGTATTTAGAAAAATAAACAAATAGGGGTTCCGCGCACATTTCCCCGAAAAGTGCCACCTGACGTCGCTAGCTGTACAAAAAAGCAGGCTTTAAAGGAACCAATTCAGTCGACTGGATCCGGTACCAAGGTCGGGCAGGAAGAGGGCCTATTTCCCATGATTCCTTCATATTTGCATATACGATACAAGGCTGTTAGAGAGATAATTGGAATTAATTTGACTGTAAACACAAAGATATTAGTACAAAATACGTGACGTAGAAAGTAATAATTTCTTGGGTAGTTTGCAGTTTTAAAATTATGTTTTAAAATGGACTATCATATGCTTACCGTAACTTGAAAGTATTTCGATTTCTTGGCTTTATATATCTTGTGGAAAGGACGAAACACCGGGTCTTCGAGAAGACCTGTTGTAGTTCCCCGGTGGTTCTTGGAAACAAGGATTATCGGGTTACTATGATAAGGTAACACACCGAAAAGCTCTAACGCCCTGCCATCCGGCAGGGCGTTATCTTTTTGGCCGGCATGGTCCCAGCCTCCTCGCTGGCGCCGGCTGGGCAACATGCTTCGGCATGGCGAATGGGACTTTTTTTTAAGCTTGGGCCGCTCGAGGTACCTCTCTACATATGACATGTGAGCAAAAGGCCAGCAAAAGGCCAGGAACCGTAAAAAGGCCGCGTTGCTGGCGTTTTTCCATAGGCTCCGCCCCCCTGACGAGCATCACAAAAATCGACGCTCAAGTCAGAGGTGGCGAAACCCGACAGGACTATAAAGATACCAGGCGTTTCCCCCTGGAAGCTCCCTCGTGCGCTCTCCTGTTCCGACCCTGCCGCTTACCGGATACCTGTCCGCCTTTCTCCCTTCGGGAAGCGTGGCGCTTTCTCATAGCTCACGCTGTAGGTATCTCAGTTCGGTGTAGGTCGTTCGCTCCAAGCTGGGCTGTGTGCACGAACCCCCCGTTCAGCCCGACCGCTGCGCCTTATCCGGTAACTATCGTCTTGAGTCCAACCCGGTAAGACACGACTTATCGCCACTGGCAGCAGCCACTGGTAACAGGATTAGCAGAGCGAGGTATGTAGGCGGTGCTACAGAGTTCTTGAAGTGGTGGCCTAACTACGGCTACACTAGAAGAACAGTATTTGGTATCTGCGCTCTGCTGAAGCCAGTTACCTTCGGAAAAAGAGTTGGTAGCTCTTGATCCGGCAAACAAACCACCGCTGGTAGCGGTGGTTTTTTTGTTTGCAAGCAGCAGATTACGCGCAGAAAAAAAGGATCTCAAGAAGATCCTTTGATCTTTTCTACGGGGTCTGACGCTCAGTGGAACGAAAACTCACGTTAAGGGATTTTGGTCATGAGATTATCAAAAAGGATCTTCACCTAGATCCTTTTAAATTAAAAATGAAGTTTTAAATCAATCTAAAGTATATATGAGTAAACTTGGTCTGACAGTTACCAATGCTTAATCAGTGAGGCACCTATCTCAGCGATCTGTCTATTTCGTTCATCCATAGTTGCCTGACTCCCCGTCGTGTAGATAACTACGATACGGGAGGGCTTACCATCTGGCCCCAGTGCTGCAATGATACCGCGAGACCCACGCTCACCGGCTCCAGATTTATCAGCAATAAACCAGCCAGCCGGAAGGGCCGAGCGCAGAAGTGGTCCTGCAACTTTATCCGCCTCCATCCAGTCTATTA |
| GfoCas9 sgRNA backbone | ATTGTTGCCGGGAAGCTAGAGTAAGTAGTTCGCCAGTTAATAGTTTGCGCAACGTTGTTGCCATTGCTACAGGCATCGTGGTGTCACGCTCGTCGTTTGGTATGGCTTCATTCAGCTCCGGTTCCCAACGATCAAGGCGAGTTACATGATCCCCCATGTTGTGCAAAAAAGCGGTTAGCTCCTTCGGTCCTCCGATCGTTGTCAGAAGTAAGTTGGCCGCAGTGTTATCACTCATGGTTATGGCAGCACTGCATAATTCTCTTACTGTCATGCCATCCGTAAGATGCTTTTCTGTGACTGGTGAGTACTCAACCAAGTCATTCTGAGAATAGTGTATGCGGCGACCGAGTTGCTCTTGCCCGGCGTCAATACGGGATAATACCGCGCCACATAGCAGAACTTTAAAAGTGCTCATCATTGGAAAACGTTCTTCGGGGCGAAAACTCTCAAGGATCTTACCGCTGTTGAGATCCAGTTCGATGTAACCCACTCGTGCACCCAACTGATCTTCAGCATCTTTTACTTTCACCAGCGTTTCTGGGTGAGCAAAAACAGGAAGGCAAAATGCCGCAAAAAAGGGAATAAGGGCGACACGGAAATGTTGAATACTCATACTCTTCCTTTTTCAATATTATTGAAGCATTTATCAGGGTTATTGTCTCATGAGCGGATACATATTTGAATGTATTTAGAAAAATAAACAAATAGGGGTTCCGCGCACATTTCCCCGAAAAGTGCCACCTGACGTCGCTAGCTGTACAAAAAAGCAGGCTTTAAAGGAACCAATTCAGTCGACTGGATCCGGTACCAAGGTCGGGCAGGAAGAGGGCCTATTTCCCATGATTCCTTCATATTTGCATATACGATACAAGGCTGTTAGAGAGATAATTGGAATTAATTTGACTGTAAACACAAAGATATTAGTACAAAATACGTGACGTAGAAAGTAATAATTTCTTGGGTAGTTTGCAGTTTTAAAATTATGTTTTAAAATGGACTATCATATGCTTACCGTAACTTGAAAGTATTTCGATTTCTTGGCTTTATATATCTTGTGGAAAGGACGAAACACCGGGTCTTCGAGAAGACCTGTCATAGTTCCCTAATAGCTCTTGGAAACAAGAACTATTATGGTTGCTATGATAAGGTCATAGGACCGTAAAGCTCTGACGCCCTGCTATTTGGCAGGGCGTCATCTTTTTTGGCCGGCATGGTCCCAGCCTCCTCGCTGGCGCCGGCTGGGCAACATGCTTCGGCATGGCGAATGGGACTTTTTTTTAAGCTTGGGCCGCTCGAGGTACCTCTCTACATATGACATGTGAGCAAAAGGCCAGCAAAAGGCCAGGAACCGTAAAAAGGCCGCGTTGCTGGCGTTTTTCCATAGGCTCCGCCCCCCTGACGAGCATCACAAAAATCGACGCTCAAGTCAGAGGTGGCGAAACCCGACAGGACTATAAAGATACCAGGCGTTTCCCCCTGGAAGCTCCCTCGTGCGCTCTCCTGTTCCGACCCTGCCGCTTACCGGATACCTGTCCGCCTTTCTCCCTTCGGGAAGCGTGGCGCTTTCTCATAGCTCACGCTGTAGGTATCTCAGTTCGGTGTAGGTCGTTCGCTCCAAGCTGGGCTGTGTGCACGAACCCCCCGTTCAGCCCGACCGCTGCGCCTTATCCGGTAACTATCGTCTTGAGTCCAACCCGGTAAGACACGACTTATCGCCACTGGCAGCAGCCACTGGTAACAGGATTAGCAGAGCGAGGTATGTAGGCGGTGCTACAGAGTTCTTGAAGTGGTGGCCTAACTACGGCTACACTAGAAGAACAGTATTTGGTATCTGCGCTCTGCTGAAGCCAGTTACCTTCGGAAAAAGAGTTGGTAGCTCTTGATCCGGCAAACAAACCACCGCTGGTAGCGGTGGTTTTTTTGTTTGCAAGCAGCAGATTACGCGCAGAAAAAAAGGATCTCAAGAAGATCCTTTGATCTTTTCTACGGGGTCTGACGCTCAGTGGAACGAAAACTCACGTTAAGGGATTTTGGTCATGAGATTATCAAAAAGGATCTTCACCTAGATCCTTTTAAATTAAAAATGAAGTTTTAAATCAATCTAAAGTATATATGAGTAAACTTGGTCTGACAGTTACCAATGCTTAATCAGTGAGGCACCTATCTCAGCGATCTGTCTATTTCGTTCATCCATAGTTGCCTGACTCCCCGTCGTGTAGATAACTACGATACGGGAGGGCTTACCATCTGGCCCCAGTGCTGCAATGATACCGCGAGACCCACGCTCACCGGCTCCAGATTTATCAGCAATAAACCAGCCAGCCGGAAGGGCCGAGCGCAGAAGTGGTCCTGCAACTTTATCCGCCTCCATCCAGTCTATTA |
| Nme2Cas9 sgRNA backbone | ATTGTTGCCGGGAAGCTAGAGTAAGTAGTTCGCCAGTTAATAGTTTGCGCAACGTTGTTGCCATTGCTACAGGCATCGTGGTGTCACGCTCGTCGTTTGGTATGGCTTCATTCAGCTCCGGTTCCCAACGATCAAGGCGAGTTACATGATCCCCCATGTTGTGCAAAAAAGCGGTTAGCTCCTTCGGTCCTCCGATCGTTGTCAGAAGTAAGTTGGCCGCAGTGTTATCACTCATGGTTATGGCAGCACTGCATAATTCTCTTACTGTCATGCCATCCGTAAGATGCTTTTCTGTGACTGGTGAGTACTCAACCAAGTCATTCTGAGAATAGTGTATGCGGCGACCGAGTTGCTCTTGCCCGGCGTCAATACGGGATAATACCGCGCCACATAGCAGAACTTTAAAAGTGCTCATCATTGGAAAACGTTCTTCGGGGCGAAAACTCTCAAGGATCTTACCGCTGTTGAGATCCAGTTCGATGTAACCCACTCGTGCACCCAACTGATCTTCAGCATCTTTTACTTTCACCAGCGTTTCTGGGTGAGCAAAAACAGGAAGGCAAAATGCCGCAAAAAAGGGAATAAGGGCGACACGGAAATGTTGAATACTCATACTCTTCCTTTTTCAATATTATTGAAGCATTTATCAGGGTTATTGTCTCATGAGCGGATACATATTTGAATGTATTTAGAAAAATAAACAAATAGGGGTTCCGCGCACATTTCCCCGAAAAGTGCCACCTGACGTCGCTAGCTGTACAAAAAAGCAGGCTTTAAAGGAACCAATTCAGTCGACTGGATCCGGTACCAAGGTCGGGCAGGAAGAGGGCCTATTTCCCATGATTCCTTCATATTTGCATATACGATACAAGGCTGTTAGAGAGATAATTGGAATTAATTTGACTGTAAACACAAAGATATTAGTACAAAATACGTGACGTAGAAAGTAATAATTTCTTGGGTAGTTTGCAGTTTTAAAATTATGTTTTAAAATGGACTATCATATGCTTACCGTAACTTGAAAGTATTTCGATTTCTTGGCTTTATATATCTTGTGGAAAGGACGAAACACCGGGTCTTCGAGAAGACCTGTTGTAGCTCCCTTTCTCATTTCGGAAACGAAATGAGAACCGTTGCTACAATAAGGCCGTCTGAAAAGATGTGCCGCAACGCTCTGCCCCTTAAAGCTTCTGCTTTAAGGGGCATCGTTTTTGGCCGGCATGGTCCCAGCCTCCTCGCTGGCGCCGGCTGGGCAACATGCTTCGGCATGGCGAATGGGACTTTTTTTTAAGCTTGGGCCGCTCGAGGTACCTCTCTACATATGACATGTGAGCAAAAGGCCAGCAAAAGGCCAGGAACCGTAAAAAGGCCGCGTTGCTGGCGTTTTTCCATAGGCTCCGCCCCCCTGACGAGCATCACAAAAATCGACGCTCAAGTCAGAGGTGGCGAAACCCGACAGGACTATAAAGATACCAGGCGTTTCCCCCTGGAAGCTCCCTCGTGCGCTCTCCTGTTCCGACCCTGCCGCTTACCGGATACCTGTCCGCCTTTCTCCCTTCGGGAAGCGTGGCGCTTTCTCATAGCTCACGCTGTAGGTATCTCAGTTCGGTGTAGGTCGTTCGCTCCAAGCTGGGCTGTGTGCACGAACCCCCCGTTCAGCCCGACCGCTGCGCCTTATCCGGTAACTATCGTCTTGAGTCCAACCCGGTAAGACACGACTTATCGCCACTGGCAGCAGCCACTGGTAACAGGATTAGCAGAGCGAGGTATGTAGGCGGTGCTACAGAGTTCTTGAAGTGGTGGCCTAACTACGGCTACACTAGAAGAACAGTATTTGGTATCTGCGCTCTGCTGAAGCCAGTTACCTTCGGAAAAAGAGTTGGTAGCTCTTGATCCGGCAAACAAACCACCGCTGGTAGCGGTGGTTTTTTTGTTTGCAAGCAGCAGATTACGCGCAGAAAAAAAGGATCTCAAGAAGATCCTTTGATCTTTTCTACGGGGTCTGACGCTCAGTGGAACGAAAACTCACGTTAAGGGATTTTGGTCATGAGATTATCAAAAAGGATCTTCACCTAGATCCTTTTAAATTAAAAATGAAGTTTTAAATCAATCTAAAGTATATATGAGTAAACTTGGTCTGACAGTTACCAATGCTTAATCAGTGAGGCACCTATCTCAGCGATCTGTCTATTTCGTTCATCCATAGTTGCCTGACTCCCCGTCGTGTAGATAACTACGATACGGGAGGGCTTACCATCTGGCCCCAGTGCTGCAATGATACCGCGAGACCCACGCTCACCGGCTCCAGATTTATCAGCAATAAACCAGCCAGCCGGAAGGGCCGAGCGCAGAAGTGGTCCTGCAACTTTATCCGCCTCCATCCAGTCTATTA |
| SpCas9 sgRNA backbone | ATTGTTGCCGGGAAGCTAGAGTAAGTAGTTCGCCAGTTAATAGTTTGCGCAACGTTGTTGCCATTGCTACAGGCATCGTGGTGTCACGCTCGTCGTTTGGTATGGCTTCATTCAGCTCCGGTTCCCAACGATCAAGGCGAGTTACATGATCCCCCATGTTGTGCAAAAAAGCGGTTAGCTCCTTCGGTCCTCCGATCGTTGTCAGAAGTAAGTTGGCCGCAGTGTTATCACTCATGGTTATGGCAGCACTGCATAATTCTCTTACTGTCATGCCATCCGTAAGATGCTTTTCTGTGACTGGTGAGTACTCAACCAAGTCATTCTGAGAATAGTGTATGCGGCGACCGAGTTGCTCTTGCCCGGCGTCAATACGGGATAATACCGCGCCACATAGCAGAACTTTAAAAGTGCTCATCATTGGAAAACGTTCTTCGGGGCGAAAACTCTCAAGGATCTTACCGCTGTTGAGATCCAGTTCGATGTAACCCACTCGTGCACCCAACTGATCTTCAGCATCTTTTACTTTCACCAGCGTTTCTGGGTGAGCAAAAACAGGAAGGCAAAATGCCGCAAAAAAGGGAATAAGGGCGACACGGAAATGTTGAATACTCATACTCTTCCTTTTTCAATATTATTGAAGCATTTATCAGGGTTATTGTCTCATGAGCGGATACATATTTGAATGTATTTAGAAAAATAAACAAATAGGGGTTCCGCGCACATTTCCCCGAAAAGTGCCACCTGACGTCGCTAGCTGTACAAAAAAGCAGGCTTTAAAGGAACCAATTCAGTCGACTGGATCCGGTACCAAGGTCGGGCAGGAAGAGGGCCTATTTCCCATGATTCCTTCATATTTGCATATACGATACAAGGCTGTTAGAGAGATAATTGGAATTAATTTGACTGTAAACACAAAGATATTAGTACAAAATACGTGACGTAGAAAGTAATAATTTCTTGGGTAGTTTGCAGTTTTAAAATTATGTTTTAAAATGGACTATCATATGCTTACCGTAACTTGAAAGTATTTCGATTTCTTGGCTTTATATATCTTGTGGAAAGGACGAAACACCGGGTCTTCGAGAAGACCTGTTTTAGAGCTAGAAATAGCAAGTTAAAATAAGGCTAGTCCGTTATCAACTTGAAAAAGTGGCACCGAGTCGGTGCTTTTTTGGCCGGCATGGTCCCAGCCTCCTCGCTGGCGCCGGCTGGGCAACATGCTTCGGCATGGCGAATGGGACTTTTTTTTAAGCTTGGGCCGCTCGAGGTACCTCTCTACATATGACATGTGAGCAAAAGGCCAGCAAAAGGCCAGGAACCGTAAAAAGGCCGCGTTGCTGGCGTTTTTCCATAGGCTCCGCCCCCCTGACGAGCATCACAAAAATCGACGCTCAAGTCAGAGGTGGCGAAACCCGACAGGACTATAAAGATACCAGGCGTTTCCCCCTGGAAGCTCCCTCGTGCGCTCTCCTGTTCCGACCCTGCCGCTTACCGGATACCTGTCCGCCTTTCTCCCTTCGGGAAGCGTGGCGCTTTCTCATAGCTCACGCTGTAGGTATCTCAGTTCGGTGTAGGTCGTTCGCTCCAAGCTGGGCTGTGTGCACGAACCCCCCGTTCAGCCCGACCGCTGCGCCTTATCCGGTAACTATCGTCTTGAGTCCAACCCGGTAAGACACGACTTATCGCCACTGGCAGCAGCCACTGGTAACAGGATTAGCAGAGCGAGGTATGTAGGCGGTGCTACAGAGTTCTTGAAGTGGTGGCCTAACTACGGCTACACTAGAAGAACAGTATTTGGTATCTGCGCTCTGCTGAAGCCAGTTACCTTCGGAAAAAGAGTTGGTAGCTCTTGATCCGGCAAACAAACCACCGCTGGTAGCGGTGGTTTTTTTGTTTGCAAGCAGCAGATTACGCGCAGAAAAAAAGGATCTCAAGAAGATCCTTTGATCTTTTCTACGGGGTCTGACGCTCAGTGGAACGAAAACTCACGTTAAGGGATTTTGGTCATGAGATTATCAAAAAGGATCTTCACCTAGATCCTTTTAAATTAAAAATGAAGTTTTAAATCAATCTAAAGTATATATGAGTAAACTTGGTCTGACAGTTACCAATGCTTAATCAGTGAGGCACCTATCTCAGCGATCTGTCTATTTCGTTCATCCATAGTTGCCTGACTCCCCGTCGTGTAGATAACTACGATACGGGAGGGCTTACCATCTGGCCCCAGTGCTGCAATGATACCGCGAGACCCACGCTCACCGGCTCCAGATTTATCAGCAATAAACCAGCCAGCCGGAAGGGCCGAGCGCAGAAGTGGTCCTGCAACTTTATCCGCCTCCATCCAGTCTATTA |

Green highlighted: Human U6 promoter

Yellow highlighted: gRNA Scaffold

Red highlighted: BbsI restriction sites

**Table S12. Primers used for PCR amplification of U6-sgRNA cassettes from positive clones and for the Sanger sequencing of expression constructs**

| **Primer Name** | **Sequence (5’---3’)** | **Used for** |
| --- | --- | --- |
| U6 gRNA-seq-F | CGGTACCAAGGTCGGGCAGG | PCR amplification of U6-sgRNA cassettes, Sanger sequencing |
| HDV gRNA-seq-R | GAGGTACCTCGAGCGGCCC | PCR amplification of U6-sgRNA cassettes, Sanger sequencing |
| T7-F | TAATACGACTCACTATAGGG | Sanger sequencing |
| CMV-F | CGCAAATGGGCGGTAGGCGTG | Sanger sequencing |
| BGH-rev | TAGAAGGCACAGTCGAGG | Sanger sequencing |

**Table S13.** **Genomic on-target sequences**

| **Name** | **Ampicon sequence (5’---3’)** |
| --- | --- |
| *CLTA* target-1 | GCTCCTCAGAGCACAGTTGTACCTCAATTGTGGATTTTAGATGTTTCTGCTTCTCAATGTTCTCTCTTTTTTCCTGCCTGCTTGCCTGCCTTTTGGACCTCTTGCTGTCTAGGGTGGCAGATGAAGCTTTCTACAAACAACCCTTCGCTGACGTGATTGGTTATGTGTATGTTGCAAAACTAATTCCTTTTTCTACATCTGATTCTTTCTACTTTTGTTTAGAGTTAATGCTCCTTTTATGTCACAAGTTGCTTTGCTTTTTAAATTATTTCAAATTGGCACTTTGGGGGCTGCCTAAGAATTGATAAGCGGGGTATGATCTGTTGATGAATCTTCCAGATTTGTGACTCCCTGCATCC |
| *CLTA* target-2 | TTTGGTAATCACTGTGTGCTATTTATCGACTATAGATTTTAGGCTTCAACTGGTTATTAACCTCATTCACTCACTTGTTCACAAAATATATGCTCAGTACTGATATATGCATACCTTATAACAACTCACGTATGTATAAAGCCTAGTCAGTTGAAGTATGAGCAAAGGTAAATACCTGTTTCTCTTCCAATCTGACGTGTTCCCAGGTTGATCACAGGTGTTCCAAAAGCTCCAACTTCTCGAACCCCACAGCTGCTGTTCCCAATCATACAGCCAGCATGGGCAACCAACTGTATAAACTGGTCAAATGGGACGTGTTTAACTGCACGAAAGTTGGGATGATGCTCAATGCCCTTCTTCCGCATCACTCGAACCAT |
| *CLTA* target-3 | TGAAGTTCTGGATTCTGGGCTGCACCTTATCAAAGCCAATTGAGTCTGGTGTAAAAATATTCAGAAACTGTACCCGCTTCTTGAGTAGAGAGAGAGCCATATTCAGGAACAACAGAAGCCAGGGCCAGGATCCTAGTCCAGTGTTCTTCCCACCAGACCACTCACCTCCAGCCGTTTTTTCCTGTAAGGTTGAGATTTTAAAATGACGATATGTGTTCCCAGGATTCTGGAAGTGTATCCCACCCAGACAGTGTGGGGAAGCAGTAGTTAGATAATCCAAGATAAAACTGGCTTTGGGTCCTAGCGCATCTGGTGTCTGGCAGCATTCCTGCCCAGTGCATTAACTGAAACAGGAAGGTGGCGTGGCTTTCTGGGCACATACAGCAGGGAAGAAGCATGAC |
| *CLTA* target-4 | AGCACCTTGCATTGCCAAAATCAGAGCTGTGGGGCAGCATTTTCCATACTTGGTTCATTCAAACAATGTGGGGTCGTTTGCCTCTCCCTCAGGCAGAAGAGTCCAGAGCAGAGGGGGTTGTCTCCAGTGGGTGTCTATAATTGTTCCTCTGTGGTCTTGCATGTCAGAAATTGCTATTTAGTGGTCAGCTTCTGATGTCCTGTCTCCTAGCACAGGGAAGCCCCCAGCTTTGAGATCAGGATAGTCAAAGTGTCTTGATACGGGCCACTTGGGCTTCTCCCCCTGACCCCCCCAAAAGAAACCTGCATCTTCTCTGGAGA |
| *HBB* target-1 | GCAATCATTCGTCTGTTTCCCATTCTAAACTGTACCCTGTTACTTCTCCCCTTCCTATGACATGAACTTAACCATAGAAAAGAAGGGGAAAGAAAACATCAAGGGTCCCATAGACTCACCCTGAAGTTCTCAGGATCCACGTGCAGCTTGTCACAGTGCAGCTCACTCAGTGTGGCAAAGGTGCCCTTGAGGTTGTCCAGGTGAGCCAGGCCATCACTAAAGGCACCGAGCACTTTCTTGCCATGAGCCTTCACCTTAGGGTTGCCCATAACAGCATCAGGAGTGGACAGATCCCCAAAGGACTCAAAGAACCTCTGGGTCCAAGGGTAGACCACCAGCAG |
| *HBB* target-2 | GCCACCACTTTCTGATAGGCAGCCTGCACTGGTGGGGTGAATTCTTTGCCAAAGTGATGGGCCAGCACACAGACCAGCACGTTGCCCAGGAGCTGTGGGAGGAAGATAAGAGGTATGAACATGATTAGCAAAAGGGCCTAGCTTGGACTCAGAATAATCCAGCCTTATCCCAACCATAAAATAAAAGCAGAATGGTAGCTGGATTGTAGCTGCTATTAGCAATATGAAACCTCTTACATCAGTTACAATTTATATGCAGAAATATTTATATGCAGAAATATTGCTATTGCCTTAACCCAGAAATTATCACTGTTATTCTTTAGAATGGTGCAAAGAGGCATGATAC |
| *HBB* target-3 | CTGACCTCCCACATTCCCTTTTTAGTAAAATATTCAGAAATAATTTAAATACATCATTGCAATGAAAATAAATGTTTTAGGCAGAATCCAGATGCTCAAGGCCCTTCATAATATCCCCCAGTTTAGTAGTTGGACTTAGGGAACAAAGGAACCTTTAATAGAAATTGGACAGCAAGAAAGCGAGCTTAGTGATACTTGTGGGCCAGGGCATTAGCCACACCAGCCACCACTTTCTGATAGGCAGCCTGCACTGGTGGGGTGAATTCTTTGCCAAAGTGATGGGCCAGCACACAGACCAGCACGTTGCCCAGGAGCTGTGGGAGGAAGATAAGAGGTATGAACATGATTAGCAAAAGGGCCTAGCTTGGACTCAGAATAATCCAGCCTTATCCCAACCAT |
| *HBB* target-4 | TAGCTGCTATTAGCAATATGAAACCTCTTACATCAGTTACAATTTATATGCAGAAATATTTATATGCAGAAATATTGCTATTGCCTTAACCCAGAAATTATCACTGTTATTCTTTAGAATGGTGCAAAGAGGCATGATACATTGTATCATTATTGCCCTGAAAGAAAGAGATTAGGGAAAGTATTAGAAATAAGATAAACAAAAAAGTATATTAAAAGAAGAAAGCATTTTTTAAAATTACAAATGCAAAATTACCCTGATTTGGTCAATATGTGTACACATATTAAAACATTACACTTTAACCCATAAATATGTATAATGATTATGTATCAATTAAAAATAAAAGAAAATAAAGTAGGGAGATTATGAATATGCAAATAAGCACACATATATTCCAAATAGTAATGTACTAGGCAGACTGTGT |
| *AIFM* target-1 | GGCTGACAACTTCTTATTGACCTGGAGTTTTGGGGTGGTGATGGAAATGTTCTGAAATCAGATAGTGGTGATGGTGGCACAACCTTGTGAATATACTGAAATCCACAGAATTCCACACTTTAAAATGGTGAATTTGATGTGAATTATATCTCAATAAAACTTTTTCAATGCTAATTCATCTCTACCTCTTTTGTGTATGTGTGTGACTGTACCAGGTAGTACAGCTGGATGTGAGAGACAACATGGTGAAACTTAATGATGGCTCTCAAATAACCTATGAAAAGTGCTTGATTGCAACAGGTGAGCATTTCTGGAGATGGCTTTCTTCCCTGATACAGCTCATTCAAGTTTTAGAGTCCAGCCTTCTCTATGGAAGG |
| *AIFM* target-2 | CGATCAGCTTAGCAGGTCACAGAGGAATTATTTTGTGGGTGTTTTTTTCCTAACTTGGCAGGGTTTACTGTGACTAAGCCGATCAGTCTAAAGTTGGATCCTGTAAGGTTGAGTGAACCAACTTAGATAACACCTAGTAACACCCTTGATTGAAGGAAGCTGTTTGTTTTACTTAAGCCATTACAAGTTTAGGGGTTTTTTTCTACATTGGTAAGACAATGAATTTATCTTTGTTCCTTGGTTGCTAGCTTAAAGAAAGGGACATGAAAATTATTTGAAGCTACCTATATCCTCATCCCCTGATGTGACTCCCAAACGGACA |
| *Casp-3* target-1 | AGGCCTAGTAGGGTGTGTGACCCATAGTTGGAAACCTAGTCCTGGAACCAAAAGGACACCACAGGAGAGGCAGGACTCCCAAGCGCCGGTGCTTTCTACCCTCCCGAATCGCTCTATAGGATTAGAGGGTTTAGGTTTGGGAGGAGTTTCTTGACCTCTTGTTTATGAGTTTTTACGACTAAAATGCAATGCCAGTTTTCAGTCCGGGGACAAACTGCCTAGTTATGGATGAATTTTACCCTTTTTTTCCCCGTGTCTTTTTGGAATATTCTGCTTATTAATGCTTCCAATAACTAGCAGATGGAAAAAGGAAAAAGATAAAACTCAAATTCACTTTTTAATTCTTGTCTGTTATTATTTATTTGGGATCTCTCGTAATACTTTAAATTCTGAAAGTTTATTTCTGAAACCCAGTATTGTTATTCTCGGCTGCCCTGA |
| *Casp-3* target-2 | AAGGAATGACATCTCGGTCTGGTACAGATGTCGATGCAGCAAACCTCAGGGAAACATTCAGAAACTTGAAATATGAAGTCAGGAATAAAAATGATCTTACACGTGAAGAAATTGTGGAATTGATGCGTGATGGTAAGAAGAAACAATTAGAATGAACTTCATTGTACAATTAATTTTATACTATGGTCTAGCAATAATTATATGTATAATAAAATATGAGAATTGTGGAATTATACTGTACATAACCCTAATCTTACATACTTTAGTAAGAAGTTTAAAAAAAGTTATTTATGCCAACTTCCTAAAATGGTTTGAGATGTGTTGCCG |
| *EMX* target-1 | TATGGAAAAGAGCATGGGGCTGGCCCGTGGGGTGGTGTCCACTTTAGGCCCTGTGGGAGATCATGGGAACCCACGCAGTGGGTCATAGGCTCTCTCATTTACTACTCACATCCACTCTGTGAAGAAGCGATTATGATCTCTCCTCTAGAAACTCGTAGAGTCCCATGTCTGCCGGCTTCCAGAGCCTGCACTCCTCCACCTTGGCTTGGCTTTGCTGGGGCTAGAGGAGCTAGGATGCACAGCAGCTCTGTGACCCTTTGTTTGAGAGGAACAGGAAAACCACCCTTCTCTCTGGCCCACTGTGTCCTCTTCCTGCCCTGCCATCCCCTTCTGTGAATGTTAGACCCATGGGAGCAG |

**Table S14.** **Information of all guide RNA, PAM types along with actual PAM sequences for each Cas9 orthologs across all target sites**

| **Target** | **Cas9 Protein** | **PAM (5’---3’)** | **Actual PAM** | **Spacer (5’---3’)** | **Orientation on genomic target** |
| --- | --- | --- | --- | --- | --- |
| *CLTA* target-1 | SpCas9 | NGG | GGG | TAACCAATCACGTCAGCGAA | Antisense |
| *CLTA* target-1 | Nme2Cas9 and NsuCas9 | N_4_CC | ACAACC | TGGCAGATGAAGCTTTCTACAA | Sense |
| *CLTA* target-1 | PsuCas9 and GfoCas9 | N_4_ATAA | CTGTATAA | TACAGCCAGCATGGGCAACCAA | Sense |
| *CLTA* target-1 | PsuCas9 and GfoCas9 | N_4_GAAA | TGTAGAAA | ATCACGTCAGCGAAGGGTTGTT | Antisense |
| *CLTA* target-1 | PsuCas9 and GfoCas9 | N_4_ACAA | ACAAACAA | AGGGTGGCAGATGAAGCTTTCT | Sense |
| *CLTA* target-2 | SpCas9 | NGG | AGG | TTCCAATCTGACGTGTTCCC | Sense |
| *CLTA* target-2 | Nme2Cas9 and NsuCas9 | N_4_CC | TGTTCC | CTGTTTCTCTTCCAATCTGACG | Sense |
| *CLTA* target-2 | PsuCas9 and GfoCas9 | N_4_ATAA | CCTTATAA | GCTCAGTACTGATATATGCATA | Sense |
| *CLTA* target-2 | PsuCas9 and GfoCas9 | N_4_GAAA | AAGAGAAA | ACCTGGGAACACGTCAGATTGG | Antisense |
| *CLTA* target-2 | PsuCas9 and GfoCas9 | N_4_ACAA | TATAACAA | CAGTACTGATATATGCATACCT | Sense |
| *CLTA* target-3 | SpCas9 | NGG | TGG | AGAACACTGGACTAGGATCC | Antisense |
| *CLTA* target-3 | Nme2Cas9 and NsuCas9 | N_4_CC | CTCACC | CAGTGTTCTTCCCACCAGACCA | Sense |
| *CLTA* target-3 | PsuCas9 | N_4_ATAA | CAAGATAA | GGGAAGCAGTAGTTAGATAATC | Sense |
| *CLTA* target-3 | GfoCas9 | N_4_GAAA | AGTAGAAA | AGGAGCATTAACTCTAAACAAA | Antisense |
| *CLTA* target-3 | PsuCas9 | N_4_ACAA | AGGAACAA | GAGTAGAGAGAGAGCCATATTC | Sense |
| *CLTA* target-4 | SpCas9 | NGG | AGG | TCTGACATGCAAGACCACAG | Antisense |
| *CLTA* target-4 | Nme2Cas9 and NsuCas9 | N_4_CC | CTGACC | CTAGGAGACAGGACATCAGAAG | Antisense |
| *CLTA* target-4 | PsuCas9 | N_4_ATAA | TGTTATAA | TAGGCTTTATACATACGTGAGT | Antisense |
| *CLTA* target-4 | GfoCas9 | N_4_GAAA | GTCAGAAA | TTGTTCCTCTGTGGTCTTGCAT | Sense |
| *CLTA* target-4 | PsuCas9 | N_4_ACAA | AGGAACAA | TTTCTGACATGCAAGACCACAG | Antisense |
| *HBB* target-1 | SpCas9 | NGG | AGG | TTGCCATGAGCCTTCACCTT | Sense |
| *HBB* target-1 | Nme2Cas9 and NsuCas9 | N_4_CC | GGCACC | GGTGAGCCAGGCCATCACTAAA | Sense |
| *HBB* target-1 | PsuCas9 and GfoCas9 | N_4_ATAA | GCCCATAA | CATGAGCCTTCACCTTAGGGTT | Sense |
| *HBB* target-1 | PsuCas9 and GfoCas9 | N_4_GAAA | GCAAGAAA | AACCCTAAGGTGAAGGCTCATG | Antisense |
| *HBB* target-1 | PsuCas9 and GfoCas9 | N_4_ACAA | TGTGACAA | TGCCACACTGAGTGAGCTGCAC | Antisense |
| *HBB* target-2 | SpCas9 | NGG | AGG | GATTATTCTGAGTCCAAGCT | Antisense |
| *HBB* target-2 | Nme2Cas9 and NsuCas9 | N_4_CC | CCAACC | CTCAGAATAATCCAGCCTTATC | Sense |
| *HBB* target-2 | PsuCas9 and GfoCas9 | N_4_ATAA | AACCATAA | CAGAATAATCCAGCCTTATCCC | Sense |
| *HBB* target-2 | PsuCas9 and GfoCas9 | N_4_GAAA | ATATGAAA | TGGATTGTAGCTGCTATTAGCA | Sense |
| *HBB* target-2 | PsuCas9 and GfoCas9 | N_4_ACAA | AGCTACAA | GAGGTTTCATATTGCTAATAGC | Antisense |
| *HBB* target-3 | SpCas9 | NGG | TGG | ATATCCCCCAGTTTAGTAGT | Sense |
| *HBB* target-3 | Nme2Cas9 and NsuCas9 | N_4_CC | TTCACC | GGCCCATCACTTTGGCAAAGAA | Antisense |
| *HBB* target-3 | PsuCas9 | N_4_ATAA | CTTCATAA | CAGAATCCAGATGCTCAAGGCC | Sense |
| *HBB* target-3 | GfoCas9 | N_4_GAAA | AATAGAAA | CTTAGGGAACAAAGGAACCTTT | Sense |
| *HBB* target-3 | PsuCas9 | N_4_ACAA | GGGAACAA | CCCAGTTTAGTAGTTGGACTTA | Sense |
| *HBB* target-4 | SpCas9 | NGG | AGG | TTCTTTAGAATGGTGCAAAG | Sense |
| *HBB* target-4 | Nme2Cas9 and NsuCas9 | N_4_CC | TGCACC | GATACAATGTATCATGCCTCTT | Antisense |
| *HBB* target-4 | PsuCas9 | N_4_ATAA | AGTGATAA | TTGCACCATTCTAAAGAATAAC | Antisense |
| *HBB* target-4 | GfoCas9 | N_4_GAAA | GAAAGAAA | TACATTGTATCATTATTGCCCT | Sense |
| *HBB* target-4 | PsuCas9 | N_4_ACAA | ATAAACAA | TAGGGAAAGTATTAGAAATAAG | Sense |
| *AIFM* target-1 | SpCas9 | NGG | TGG | GCTGGATGTGAGAGACAACA | Sense |
| *AIFM* target-1 | Nme2Cas9 and NsuCas9 | N_4_CC | ACTACC | GTTGTCTCTCACATCCAGCTGT | Antisense |
| *AIFM* target-1 | PsuCas9 and GfoCas9 | N_4_ATAA | TCAAATAA | TGGTGAAACTTAATGATGGCTC | Sense |
| *AIFM* target-1 | PsuCas9 and GfoCas9 | N_4_GAAA | TGGTGAAA | CAGCTGGATGTGAGAGACAACA | Sense |
| *AIFM* target-1 | PsuCas9 and GfoCas9 | N_4_ACAA | AGAGACAA | CCAGGTAGTACAGCTGGATGTG | Sense |
| *AIFM* target-2 | SpCas9 | NGG | AGG | CTAGTAACACCCTTGATTGA | Sense |
| *AIFM* target-2 | Nme2Cas9 and NsuCas9 | N_4_CC | TCAACC | GGTGTTATCTAAGTTGGTTCAC | Antisense |
| *AIFM* target-2 | PsuCas9 | N_4_ATAA | AAAGATAA | TTTAAGCTAGCAACCAAGGAAC | Antisense |
| *AIFM* target-2 | GfoCas9 | N_4_GAAA | TAAAGAAA | CTTTGTTCCTTGGTTGCTAGCT | Sense |
| *AIFM* target-2 | PsuCas9 | N_4_ACAA | TAAAACAA | CCTAAACTTGTAATGGCTTAAG | Antisense |
| *Casp-3* target-1 | SpCas9 | NGG | GGG | GCAATGCCAGTTTTCAGTCC | Sense |
| *Casp-3* target-1 | Nme2Cas9 and NsuCas9 | N_4_CC | TTTACC | AACTGCCTAGTTATGGATGAAT | Sense |
| *Casp-3* target-1 | PsuCas9 and GfoCas9 | N_4_ATAA | ATTAATAA | CCATCTGCTAGTTATTGGAAGC | Antisense |
| *Casp-3* target-1 | PsuCas9 and GfoCas9 | N_4_GAAA | AAAGGAAA | TTCCAATAACTAGCAGATGGAA | Sense |
| *Casp-3* target-1 | PsuCas9 and GfoCas9 | N_4_ACAA | GGGGACAA | ATGCAATGCCAGTTTTCAGTCC | Sense |
| *Casp-3* target-2 | SpCas9 | NGG | TGG | TCTTACACGTGAAGAAATTG | Sense |
| *Casp-3* target-2 | Nme2Cas9 and NsuCas9 | N_4_CC | ATAACC | GAATTGTGGAATTATACTGTAC | Sense |
| *Casp-3* target-2 | PsuCas9 | N_4_ATAA | ATGTATAA | ACTATGGTCTAGCAATAATTAT | Sense |
| *Casp-3* target-2 | GfoCas9 | N_4_GAAA | AGAAGAAA | GTGGAATTGATGCGTGATGGTA | Sense |
| *Casp-3* target-2 | PsuCas9 | N_4_ACAA | AGAAACAA | GAATTGATGCGTGATGGTAAGA | Sense |
| *EMX* target-1 | SpCas9 | NGG | CGG | TCGTAGAGTCCCATGTCTGC | Sense |
| *EMX* target-1 | Nme2Cas9 and NsuCas9 | N_4_CC | GTGACC | AGCTAGGATGCACAGCAGCTCT | Sense |
| *EMX* target-1 | PsuCas9 | N_4_ATAA | GATCATAA | CTCTACGAGTTTCTAGAGGAGA | Antisense |
| *EMX* target-1 | GfoCas9 | N_4_GAAA | TCTAGAAA | AAGAAGCGATTATGATCTCTCC | Sense |
| *EMX* target-1 | PsuCas9 | N_4_ACAA | TCAAACAA | AAGGGTGGTTTTCCTGTTCCTC | Antisense |

**Table S15.** **On-target genotyping primers and Illumina adapter primers used for adding sequencing adapters to target amplicons**

| **Primer Name** | **Sequence (5’---3’)** |
| --- | --- |
| *CLTA* region-1_Miseq_F | GCTCCTCAGAGCACAGTTGTA |
| *CLTA* region-1_Miseq_R | GGATGCAGGGAGTCACAAATC |
| *CLTA* region-2_Miseq_F | TTTGGTAATCACTGTGTGCTATTT |
| *CLTA* region-2_Miseq_R | ATGGTTCGAGTGATGCGGAA |
| *AIFM* target_Miseq_F | GGCTGACAACTTCTTATTGACC |
| *AIFM* target_Miseq_R | CCTTCCATAGAGAAGGCTGGAC |
| *HBB* target-2_Miseq_F | GCCACCACTTTCTGATAGGCA |
| *HBB* target-2_Miseq_R | GTATCATGCCTCTTTGCACCAT |
| *HBB* target-1_Miseq_F | GCAATCATTCGTCTGTTTCCCA |
| *HBB* target-1_Miseq_R | CTGCTGGTGGTCTACCCTTG |
| *Casp-3* taregt_Miseq_F | AGGCCTAGTAGGGTGTGTGA |
| *Casp-3* taregt_Miseq_R | TCAGGGCAGCCGAGAATAAC |
| *CLTA* target-3_Miseq F | TGAAGTTCTGGATTCTGGGCTG |
| *CLTA* target-3_Miseq R | GTCATGCTTCTTCCCTGCTGTA |
| *CLTA* target-4_Miseq F | AGCACCTTGCATTGCCAAAATC |
| *CLTA* target-4_Miseq R | TCTCCAGAGAAGATGCAGGTTT |
| *AIFM* target-2_Miseq F | CGATCAGCTTAGCAGGTCACAG |
| *AIFM* target-2_Miseq R | TGTCCGTTTGGGAGTCACATCA |
| *HBB* target-3_Miseq F | CTGACCTCCCACATTCCCTTTT |
| *HBB* target-3_Miseq R | ATGGTTGGGATAAGGCTGGATT |
| *HBB* target-4_Miseq F | TAGCTGCTATTAGCAATATGAAACC |
| *HBB* target-4_Miseq R | ACACAGTCTGCCTAGTACATTACTAT |
| *Casp-3* target-2_Miseq F | AAGGAATGACATCTCGGTCTGG |
| *Casp-3* target-2_Miseq R | CGGCAACACATCTCAAACCATT |
| *EMX* target-1_Miseq F | TATGGAAAAGAGCATGGGGCTG |
| *EMX* target-1_Miseq R | CTGCTCCCATGGGTCTAACATT |
| *CLTA* region-1_Adapter_F | TCGTCGGCAGCGTCAGATGTGTATAAGAGACAGGCTCCTCAGAGCACAGTTGTA |
| *CLTA* region-1_Adapter_R | GTCTCGTGGGCTCGGAGATGTGTATAAGAGACAGGGATGCAGGGAGTCACAAATC |
| *CLTA* region-2_Adapter_F | TCGTCGGCAGCGTCAGATGTGTATAAGAGACAGTTTGGTAATCACTGTGTGCTATTT |
| *CLTA* region-2_Adapter_R | GTCTCGTGGGCTCGGAGATGTGTATAAGAGACAGATGGTTCGAGTGATGCGGAA |
| *AIFM* target_Adapter_F | TCGTCGGCAGCGTCAGATGTGTATAAGAGACAGGGCTGACAACTTCTTATTGACC |
| *AIFM* target_Adapter_R | GTCTCGTGGGCTCGGAGATGTGTATAAGAGACAGCCTTCCATAGAGAAGGCTGGAC |
| *HBB* target-2_Adapter_F | TCGTCGGCAGCGTCAGATGTGTATAAGAGACAGGCCACCACTTTCTGATAGGCA |
| *HBB* target-2_Adapter_R | GTCTCGTGGGCTCGGAGATGTGTATAAGAGACAGGTATCATGCCTCTTTGCACCAT |
| *HBB* target-1_Adapter_F | TCGTCGGCAGCGTCAGATGTGTATAAGAGACAGGCAATCATTCGTCTGTTTCCCA |
| *HBB* target-1_Adapter_R | GTCTCGTGGGCTCGGAGATGTGTATAAGAGACAGCTGCTGGTGGTCTACCCTTG |
| *Casp-3* taregt_Adapter_F | TCGTCGGCAGCGTCAGATGTGTATAAGAGACAGAGGCCTAGTAGGGTGTGTGA |
| *Casp-3* taregt_Adapter_R | GTCTCGTGGGCTCGGAGATGTGTATAAGAGACAGTCAGGGCAGCCGAGAATAAC |
| *CLTA* target-3_Adapter F | TCGTCGGCAGCGTCAGATGTGTATAAGAGACAGTGAAGTTCTGGATTCTGGGCTG |
| *CLTA* target-3_Adapter R | GTCTCGTGGGCTCGGAGATGTGTATAAGAGACAGGTCATGCTTCTTCCCTGCTGTA |
| *CLTA* target-4_Adapter F | TCGTCGGCAGCGTCAGATGTGTATAAGAGACAGAGCACCTTGCATTGCCAAAATC |
| *CLTA* target-4_Adapter R | GTCTCGTGGGCTCGGAGATGTGTATAAGAGACAGTCTCCAGAGAAGATGCAGGTTT |
| *AIFM* target-2_Adapter F | TCGTCGGCAGCGTCAGATGTGTATAAGAGACAGCGATCAGCTTAGCAGGTCACAG |
| *AIFM* target-2_Adapter R | GTCTCGTGGGCTCGGAGATGTGTATAAGAGACAGTGTCCGTTTGGGAGTCACATCA |
| *HBB* target-3_Adapter F | TCGTCGGCAGCGTCAGATGTGTATAAGAGACAGCTGACCTCCCACATTCCCTTTT |
| *HBB* target-3_Adapter R | GTCTCGTGGGCTCGGAGATGTGTATAAGAGACAGATGGTTGGGATAAGGCTGGATT |
| *HBB* target-4_Adapter F | TCGTCGGCAGCGTCAGATGTGTATAAGAGACAGTAGCTGCTATTAGCAATATGAAACC |
| *HBB* target-4_Adapter R | GTCTCGTGGGCTCGGAGATGTGTATAAGAGACAGACACAGTCTGCCTAGTACATTACTAT |
| *Casp-3* target-2_Adapter F | TCGTCGGCAGCGTCAGATGTGTATAAGAGACAGAAGGAATGACATCTCGGTCTGG |
| *Casp-3* target-2_Adapter R | GTCTCGTGGGCTCGGAGATGTGTATAAGAGACAGCGGCAACACATCTCAAACCATT |
| *EMX* target-1_Adapter F | TCGTCGGCAGCGTCAGATGTGTATAAGAGACAGTATGGAAAAGAGCATGGGGCTG |
| *EMX* target-1_Adapter R | GTCTCGTGGGCTCGGAGATGTGTATAAGAGACAGCTGCTCCCATGGGTCTAACATT |

**Table S16. Predicted off-target sites and off-target amplicon sequences**

| **Off-Target Name** | **Chr** | **Off-Target Sequence (5’---3’)** | **Amplicon sequence (5’---3’)** |
| --- | --- | --- | --- |
| Psu-*HBB*-T3-N_4_ATAA-OFFT1 | chr1 | CAGAATCCA**tc**TGCTCAAG**tt**C | CAGCCCCAGGACCAAATTACCATCACAGGGTACAAAAAGAATACAGAAGCTACACAAGATACTATATTGATTA**TTATGGATGAACTTGAGCAGATGGATTCTG**AGGACGTTCCATTGGACATCCCACATCACTGGTGCCTGCAGCAAAAATCATGGATCCAAAATCGTGGATGAATTCAAAGTTGACATTTGTTTCCCACAGAGAGGAGGCCCAAACCCCAACTGTGGCACCATGATGAGGCTCCCAGAGAACATGGAGAAAGCCATTGACCACATCCTCAACCTAGAGGAGGAATACCTGGCTGATGTGGTAGATAGCAAGGTGCTGCAGGTA |
| Psu-*HBB*-T3-N_4_ATAA-OFFT2 | chr11 | **t**A**a**AATC**t**AGATGCTCAAG**c**CC | GTGAAGTACAACTGCCCCTCCAAATCCATGGACTCAACCAATGGTTGACTGAATATCAAAAATAGTCAGAAAAAAAAGTATAAAATTTAAAAATACAATATAACAACTATTTACATAGCATTCACATTGTTAAGTATTATAAGTAATATAGAGATGATTAAATTATACAAGAGGATGTCTGTAAGTTATATGCAAATACTATGCCATT**TTATATCAGGGCTTGAGCATCTAGATTTTA**GTATCCCTGGAGGTTGCTGTTGCGGGGAGGGTCCTGGAACCAATCCCCTGTGGATCAAGAGAGAAGACTGTACTATTCTCCATTTTACAGACGAGACTGAGGCCCAGAGAGGTGAAGCAAATTGCCCAAGACTACACAGCACAGACCTGTAAACCTCCACAACCT |
| Psu-*HBB*-T3-N_4_ATAA-OFFT3 | chr7 | CAG**c**A**a**CCA**t**ATG**a**TCAAGGCC | GCAGGAGGTAGTTTAAGGCCTACTCAGTGAGTAATTTAGTAAAGAACAGTGAGTTTTATGACTTAAATCTCATGAAATGACATTAATGTAAGCATGGTTCATGGAATGAGAAAGTGATGAAAATGATGTTGATTGTTCTTGGACTCAAGTGTGGTACTTTAATGTA**TTATAGGTGGCCTTGATCATATGGTTGCTG**CCTTCTTCTGACAAAGCTAGCAGGAGAAAGAAAGACACAAGTGGCTCTGTTTGGGCATTACAGTGTAGCCTGGAAATGCTACATTATTGAGGATGATCTTTTGAAAGTGAGAATCTGAGGCAACTGCCTG |
| Psu-*AIFM*-T1-N_4_ATAA-OFFT1 | chr1 | TGG**a**GAAA**g**TTAA**gc**ATGGCTC | AGTTGTATTACCTGGGTTGGAGGTAAGTGTTGGGGACACAAGTGTTAGAAACAGAACAGTTTGTTTAAAAGTTGGGGACAAAAAAGGACCAGAATCAAGATTACGGCACCC**TGGAGAAAGTTAAGCATGGCTCGGTCATAA**ATTAGGAGACAAGGAACAAATAGAGAAATACGTTTGGATAGAATAGTCAGATTATAGGCTGTGCTAAGTTGTTTGGGCTCCACTAGGGTTTCTAAATGCAGTTATTGGTAAACTGGACTTAGGAGAAGGGACAGGGTTGACAT |
| Psu-*AIFM*-T1-N_4_ATAA-OFFT2 | chrX | **g**GGTGA**t**A**a**TTAATGATG**a**CTC | AGGCCAGATTCTGAGGCCTAAGAAAACTAACTTATCTAGATCAGTTTCTTTACATCCCATTTTTATCTGACCTAAACTTTCAGGCACTGGATAAGATAATCTGGCTGCCTTCAGCCATATCCTTTTCTGAAGCTTTTGTAAAACTTCCCGGCCTTCCAAGAAGGTTTGCATCTTTCTACAATTTTTCCCACCACCCTGACTGATCTACAAGC**TTATTTTGGAGTCATCATTAATTATCACCC**ATTTTTTCTGTATTTATGATAATGACCTATTGGATCAAGGAATAGAGAGAGTAGAATTTGCTCTCATTCGCTCAGGTAAGAAGATGGAGATAATTTTTCTTACTCATATCCTAGTTGAGTGGCCCTTTCCTTACAACACA |
| Psu-*AIFM*-T1-N_4_ATAA-OFFT3 | chr10 | TGG**a**GAAA**a**T**a**AAT**a**ATG**t**CTC | AGCATTTGTGAAATAGGGACTTTCTAGCCTTTATTTGTGTTTAAGGAATCAGGGAATAAGTTCAAAATTGCCTTTCAAGAAATTTTTGGAACTCTCTTCTCACTAAGAAACTGTAAAGTC**TTATAAAAGAGACATTATTTATTTTCTCCA**AGTATTGCTTGCGAGGTGAATTGAAGGTTTTTTTTTTATCAACAGTTGTTTTATAAGATTGTTTGAGGACTAAAAGGGCTGATTGTAATCACCTGTAACATGTTACCCAGCAAGACATTCCTCACCAGG |
| Psu-*EMX*-T1-N_4_ATAA-OFFT1 | chrX | CTCT**t**C**t**AGTTTCTAGA**t**GAGA | ATGGATGTCTAGTTGCTCCAGCACCATTCATTGAAAAGTCTATTCTCCCTCCATTGAGTTCCTTTTGCAGTGAAAAATCAGTTGGGCAGATTTGTGCATCTCTAATTCTGGATTCTCTGTCC**CTCTTCTAGTTTCTAGATGAGACTGTATAA**GATTGCTGTTAATTCTTCCTTAAGTGTTAAAATTCTCCAGCAAACCTATTTGAGCTTGCAGATTTCTTTTATTGGAAACTTGATAATTATAATTACATATTCAATTTCTTTAATAATTACATGACTCTTAAAGTTGTTTATTTCATCTTGGTTAACTTTTGGTAGTTTGATTTTCAAGGAAATTGGTCCATTTCTTCT |
| Psu-*EMX*-T1-N_4_ATAA-OFFT2 | chr3 | CTCT**t**C**t**A**t**TTTCT**g**GAGGAGA | GATTGTTAGGTCTATTCTGGGAACTAAGATTGCTTATTTGGAATAAATCCTACTTTTTCATGGTGTATTACAAGATTTGTTAATGCTCTATAAAGGATTTTTGCTTCTAATTTTAGGAGAGATGTTTGTTTTTTTTTGTCGTTGTTGTTGTTTTGTAATGTCTTTGGCTTTGTACTGGCCTTCTAAATAAATTAGGAAATATTTC**CTCTTCTATTTTCTGGAGGAGATTATATAA**AGTTGGTGTTAATTCCCCTTTAAAGGTTTAGTAGAATTAGTCAGTGAGACCATCTGGGTCTGGGCCTGGAGATTTTA |
| Psu-*EMX*-T1-N_4_ATAA-OFFT3 | chr17 | CTCTA**t**GA**c**TTTCTA**c**AGGA**at** | GCATGGGGAGCTGTGAAAGAAGTGGGGTCAGAGGAGTCGTCAGGACTCAGATTATACGGGACTTTTTAAGTCATGGGAAAGACTGCATT**TTATTCCAATTCCTGTAGAAAGTCATAGAG**CTCTCTAACCATGCCTCAGGAGAAAACAACCACAACAACAGTCAACTGTTGTTTTCTGTTGCTTTTATTACTTGTTAATCAATGTTTAGCAATCCATTCAACAGTAAAGCAATTTCAGCTGAATTGACTGATCACTTCTGTGACTATTCAAACTGGTTTGTACGTGAACAACCAGAGTCTACACTTGAACTAGCTACCATAGAATGTGGATTCCAAATCAATCCACCCAGTGGA |
| Psu-*CLTA*-T3-N_4_ACAA-OFFT1 | chr1 | **t**AGTAGAGAGAGAG**a**CATAT**a**C | CCCTCCTCTGTCCCGTTTCTTCAAACTTACCCCCTACAAATGGGAATTTGTTCCTTCCTTCTCTCTGTTCCCAAAACATTTCATTTAAATCTAATTTACAACATCTTTTCAATTAAGATATTATAGTATAC**TTGTTTGAGTATATGTCTCTCTCTCTACTA**AGAGAACAGTGTTAAAAGGTATGGACATTAAATGCAGGACTCAATATAAGGTCTGGCAAAGAATAAACCCTTAATAGCTCTTGGATGGAAATAAAGGATGATATGAGATTATGAATGTCCTAAAGGAATCAGGCAGAAGAAAATCTCTTCTCAATTCCATGAGAACTTTCCTAGGTCTTGGTTCTCTAAACCATGGTG |
| Psu-*CLTA*-T3-N_4_ACAA-OFFT2 | chr1 | **t**AGTA**c**AGAGAGAG**a**CATAT**a**C | CCCTCCTCTGTCCCGTTTCTTCAAACTTACCCCCTACAAATGGGAATTTGTTCCTTCCTTCTCTCTGTTCCCAAAACATTTCATTTAAATCTAATTTACAACATCTTTTCAATGAAGATATTATAGTATAC**TTGTTTGAGTATATGTCTCTCTCTGTACTA**AGAGAACAGTGTTAAAAGGTATGGACATTAAATGCAGGACTCAATATAAGGTCCGGCAAAGAATAAACCCTTAATAGCTCTTGGATGGAAATAAAGGATGATATGAGATTATGAATGTCCTAAAGGAATCAGGCAGAAGAAAATCTCTTCTCAGTTCCATGAGAACTTTCCTAGGTCTTGGTTCTCTAAACCATGGTG |
| Psu-*CLTA*-T3-N_4_ACAA-OFFT3 | chr1 | GAG**c**AGAGA**a**AGA**t**CCAT**t**TTC | TCCCTGCCTCAGAGAAAAGACTGCATGCATGCACAATGTCATGCAAATCAGCTTTAATTTACTGCATGTTTTTCTGCTTCCACCAAAT**TTGTTGAAGAAAATGGATCTTTCTCTGCTC**ATAGCTGTAAATTCCTGCATCGTCAAAGCATACACAGATCAAGATGTTGTTTTTCAACTGATGCTTGAATAATAGCAGAAGAAAATTTCTATACAGAAATATCAAATGGATATTCCCATGGCAGCATTAATCTGGAAATATTCCCCACTGCACTATAAGTGGGTCTAGTTGGGTAATAGGAGCATGGGAAGGTCCTTATTTTGCACCCAAGGAGTTGCTC |
| Psu-*AIFM*-T1-N_4_ACAA-OFFT1 | chr19 | CCAGGTA**c**T**t**CA**t**CTGGAT**t**TG | CACCAGGCCCATGTCATATGTTTTTATGAGTTTGGACATTGGCATCTGGGA**CCAGGTACTTCATCTGGATTTGAACCACAA**TGTGGGTGTTTGTTTTTAATACCTTTGTGAAATAATTTTAAATTGCCTTATATATTTGGCATTGTCACTTTGCTGCCCCTACACACTGATGGAAAAATTGTGTTCTGCAGAAAACTAGAGTTCCTCACAGGGTGAGGAAGAGTTATGGTCATCTGCACAGTTGTAGCCCGTTGCGAGTGTTTTCCCATGTGCACAGTGATATCTAGGCAAGTGTGCCATGATTTCCTGGT |
| Psu-*AIFM*-T1-N_4_ACAA-OFFT2 | chr1 | CCAGG**c**AG**a**AC**g**GCTGG**c**T**t**TG | AACCATGAGCAGAGGAGGCAGGCCTGAACACATCTGTTAGAGGAGGGAATGATGTTTCTGCTACACAGCAAGTGTACCAACAAAACAAAACAAAACAAAACAAGCCCACGTTATTTGAATGAAACAGCAGTGAAGCAAAAAACACGCAACTGGAGAGGAGATGATGGTGGGTAAGGAATCTAGACAACAGGATGGATGGAGT**CCAGGCAGAACGGCTGGCTTTGGCAAACAA**GAGAGAAAGAAGAGGGATAGGCAGAAGCTGAGAAAGCAGGTTAAAGGTTGTGGTCAAGAGATGAGGGC |
| Psu-*AIFM*-T1-N_4_ACAA-OFFT3 | chr12 | C**gg**GGTA**c**T**g**CAGCTG**t**ATGTG | CTGAGGAGCCAGGCCCTGAACACCTCGGCCAGAGCCCTCCTCGGATGGTGACTATGCCAGGAGACACCGGGGAGACCTGCACATGGGCTGGAGTGTGTTCCAGAGACTTCACAGACTTGGCAACTATCAT**CGGGGTACTGCAGCTGTATGTGTTTCACAA**AAAGTCCTTATCTTAATTCAAAAATTAATCTGGTGTGGTGGTGTGCACCTGTAGTCCCAGCTACTCAGGAGGCTGAGGTGGGAGAATTACTTGTGCCCAAGAGGTTGAGGCTGCACTGAGCAAAG |
| Psu-*EMX*-T1-N_4_ACAA-OFFT1 | chr1 | AAGGGTGGTTTTCC**ca**T**c**CC**c**C | AAGCTTTCAGGAACGGGCTTTGCCCTCTCCCTCTTGCCTACCACGGACTTTCTTGCATTAAAAATTCTCATTGTGGGAGTCTCCGCTATTACTCACTTCTTTCAAAATGGAATGCTAGCAGCTGACAAGGGGCCCGCTCATCTCCAGCACCGGCTAGATCAGGAC**AAGGGTGGTTTTCCCATCCCCCTTTCACAA**AAGGGGAAACAGAGGCCTTCAACACACAAACGAGCCTGTGTTTGCTCATCTCAAAAGGTCTCCGTGGCCTCTCCTTACAAAACAACGAGCAGACTCCCATCTACCTGGTACACTCGGGAGGACTCTAAGACTGCT |
| Psu-*EMX*-T1-N_4_ACAA-OFFT2 | chrX | AA**c**GGTGGTTTTCCT**a**TT**g**CT**g** | AAAAAGAGTCAGTCCCTCTGGGCAGGGCCTCAAAGAGCACCACCCAGGTTG**AACGGTGGTTTTCCTATTGCTGGGCCACAA**GTGCCTTCCTATTGACCTTCACCTTCTCTAGACACTTGTTCAGGTGGCTGTGGGTCATCTCCATCTCCTCAATCTTGTCCAGTGCTGTTTATAAACCTCATTGTACTTTACCCTTTCTTACTTTAAGTTCACCTGCAACTTTCACTACTTTGCCTATTGGCATATGTCTGTATTTTGATGAACTGCAAATCTGAGTCTTTCTGCAGCTGC |
| Psu-*EMX*-T1-N_4_ACAA-OFFT3 | chr11 | **g**AGG**a**TGGTTTTCCTGTT**t**CT**g** | CCAGCTGGAGGGAAAAGGAAAGCCAAGCGGCACATACTGGACAAGTGTTAGGTTTCAAAAACCCAGGGAAATCCAGATACTGCCCAAGAAAACAGGTTCTAGAACTAGGATTTTCAAATTTAGAAAATTCAAAACATCAAGTGGAAAA**GAGGATGGTTTTCCTGTTTCTGGGAAACAA**AGGCCTCCCAGAAAACCAGAAACACAGGTGGAAGTCACGAAAGAAAATATAGGTTTGGTGGATGTCTTTTCATCTACCTGTGAATGTTAGGGGCTGTGGAAGGAGAAGAGGAAGCAGGTCTGGACATGG |
| Gfo-*HBB*-T3-N_4_GAAA-OFFT1 | chr11 | CTT**t**GGGAACA**g**AGG**g**AC**t**TTT | ACTCATGAACCACGCTCACCTTTACTCCTTTGTCCTGTCCAAATGCTCTTATTCTGGATGTGGCATAACTACTAGGAGCATAGGCTTTTATAAGGACACATCTGGAGATCAATGACAACCTGAAAAAAAAAATTATTCACGGTGTTGGGGAGAGGAGCA**CTTTGGGAACAGAGGGACTTTTGGAGGAAA**CCCTCAAGCTATCCAAATTAGGATCACTGTCCCTGCTCCCTCTACTTGGTGAGCTTTAGTGAGAAGGATAGTTGCCCAATTAGTCCCCAAACACCCCTTTCTGTTTTCGCATCCTTCCACTTGAGGACAGGGCTTCTCCTTTAGCTATCCTGGACCCCACCTAATTCTGAGTTAGGGCCCTGCTTCTT |
| Gfo-*HBB*-T3-N_4_GAAA-OFFT2 | chr3 | CTTAG**a**G**c**ACAAAGG**t**ACCTT**a** | CCACGCCCATCTTAATTTGTACTTTTAGTAGAGACGGGGTTTTGCCATGTTGCCCAGGCTGGTCTCAAACTCCTGACCTCAGGTGATCCACCCGCCTTGGCCTCCCAAAGTGCTGGGATTACACGTGTGAGCCACTGTGCCCAGCTTACTAGGTAATTT**TTTCAAAGTAAGGTACCTTTGTGCTCTAAG**GCTCACCCCTCTCCAACTACTCAATGAAAAAACAAAAAAATGCTAGTACCATGTGTATTTTGATTTCATGTTAATTACATATTTTATTTGCTCAGAGTACAAATTAGATAAAATTCCTTTTATTAAAATATTCCTTAGTAAAGAATCTAGTAATTTGC |
| Gfo-*HBB*-T3-N_4_GAAA-OFFT3 | chr7 | CT**gta**GGAACAAAG**a**AACCTTT | ATCCACTTAATTTGGGCAAGGTATTCCGTATTTTGATCTGCTAGAATTTTTTTAAAAATTAGTTTTGTTTTAATCATTCTATTATGTATTTTGCTTTG**CTGTAGGAACAAAGAAACCTTTCCAAGAAA**AGCCCTAGAAGGGAAATAGTAAAATTAAAAATGTAATACCCCATCAGTAATTTTTCTAGGGTAACTCTTAACTAATCTTAAGTGAAATATTCTATAAGTTATCATCTTGACAATTCAACTACAAAAATGGATTGGGTACATAAAATGCAGAGTAAAGAATAAAAATGATTTGAAGTAGGAAATTAGGTCAATTAGCAAATAGACCATTCTGCTTCTATTCACTAGAATTATGCAACAGCTTCTGGAGGC |
| Gfo-*AIFM*-T1-N_4_GAAA-OFFT1 | chr10 | **t**AGCTG**a**A**a**GTGAGAG**g**CAACA | AGACAGTGTGATTCAGACAGTGGAATGGAAAAAGAGAGAAGCATGTATTTCCAGCCACCTCTTTCTGTGTCTCTGCCCAGGACTTGCCTCACATTGAGAATGGTGTTGTGGCCATCCTCACGAGGTAGAAGATGGTA**TAGCTGAAAGTGAGAGGCAACACAGTGAAA**CTTAATGATGGCTCCCAAATAGCCTATGAAAACTGCTTGATTGCAACAAGAGGTCCTCCAAGAAGTCTGTCTGAAATTGATGAAGCTGGATCAGAGGTGAAGAGCAGAGCAACACTCTCTAGAAAGATTGGAGACTCTAGGACTCTGGAGAAGATTTCATGGAAGTCAAGTCAATTATGGTTATCGGTGGGGGCTTGCTTGGTGGTGAACTG |
| Gfo-*AIFM*-T1-N_4_GAAA-OFFT2 | chr17 | CAGCTGGA**ac**T**c**AGA**a**ACAACA | GAGGCCATCAGTTCCTTCCTGTGGAGAAGGGTCTGAAATGGAAGTCAGTGGTAGAAGGGGCTGGTCTGCTGGGCAGGGCTTACATCCACTGAGTTCTAAGATTCCTTTCCTGATCTGCACCTACGCCTGGTCTGTATGGTGGAATTTGT**CAGCTGGAACTCAGAAACAACAACTTGAAA**AAAAAATAATAATTAGAACATATTTGCATAAGATAGCTATTTACTCTGGAAACCAACAACTTTTGAGATTTCCCTTGCCCTGTGGACGCCCAGCTCCTGTCATCCTTCCTT |
| Gfo-*AIFM*-T1-N_4_GAAA-OFFT3 | chr22 | **a**AGC**a**GGATGTGAGA**c**ACAA**a**A | TCTCAGCACTGATCTCTCAGCCTGTTTCTTGCATAAATAAAACAGAGGCACTGGTTTGCATGAGAATCTCATAATGCGCTATGAAATCCATGCCAGAGTCAGGCATCTAATTTAAGAATTA**TTTCTCGATTTTGTGTCTCACATCCTGCTT**CTGCCTCTTTCTGCTGATTGGATTCTTCCCTTCCCTTTCCCTAACAAGCTTGTCTCCCCCTCTCCCTGCACCTCCTCCTCTGATCTTTCCCTCCTGCCTTCACCTCTTCCTCTCTCCCTAATAAAGTGACCCTGGCCTGGCTAGATATCTGTGAGTTGGCAGAGAGGTCAAGACAGGTCA |
| Nsu-*AIFM*-T1-N_4_CC-OFFT1 | chr10 | GT**a**GTCTCT**g**ACATCCAGCTGT | GCTACACTCAGCAACAAGTGATAGGGACAGGAGGCAGGGAAATTCTAGGCAAAAGAGGGCGGGCCTCTAGTGAGGGCCCCACCCTCAAGCCTGGAACCATGGCTTAAAGTGAGAACATACATTCCTGTTTTCCCACTCCAAAGCTGCCTTTTCCAAAACAACCCATGGCCCACCCCACCCTTCATCTTGTGCCCATAAAAACCCAAGGCTCCAGAGGCAGAAAAGAGGAGAAGA**GGAGAAACAGCTGGATGTCAGAGACTAC**AGTCCAGGAAAAGCGGCTTGACTTCAGAGGGACAGCTTGACGGGGTAGCTCTGGGGGAAGAATACCGTCCTGCTTCATCC |
| Nsu-*AIFM*-T1-N_4_CC-OFFT2 | chrX | GT**a**GTCTCTCACATCCAGCTG**c** | AAGGTAAGCTTCCCCTGGAGTCAAGCTGTTCCTCTGAAGTCAAGCTGCTTCTCTCCAATGTCCAACT**GTAGTCTCTCACATCCAGCTGCTTCTCC**TCTCTGCTGGCTGAGCCTGGGGTTTTTATGGGCACAGGATGGGAAGGTGGGGTGGGCCATGGGTGGTTTTGGAGAAGGCAAAATTCGAGGGGGAAAAGAAGGGTGTAAGTTCTCTCTTTGGGCCATGGTTCCAGGCTTTTTGGTTTGAGGGTGGGGACTTCACTGGGGACCCTCCCTTTTCTGCCCAGAATTTCCCTGCCTTCTATTTCTATCAACAGTATAGGAATGAGTCATACTACTTGGACTGTGGTTCTCCAAC |
| Nsu-*AIFM*-T1-N_4_CC-OFFT3 | chr11 | GT**a**GTCTC**a**CACATCCAGCT**a**T | AGGGTAGAGCATGATCTGGGCATCTCCAAATAGGAAAGTTCTTATATGAGTACTCCTAATGTGTCCACCCACCAAATCTATTTTATTGCTGATGCACGTTTTGCCCTGGCCTGCACCTTGCTGCCTAGGCCTCTTGAGCTCCTGGTCCC**GTAGTCTCACACATCCAGCTATGTGTCC**GTCACACCCCAGGCTCTTGTCCTTGACTATCTTCAGACACCTCTCAGTTTATTCTTCACAGTTTGCTCTTACTCCTCTCTTTCCTTGTTGCTTTGTTTTCTGGTCCTGGGTCACATCTTCAGTAGCTGCTTTTGGTCACTTTGGTGTGC |
| Nsu-*Casp-3*-T1-N_4_CC-OFFT1 | chr16 | AA**a**TGCCTAGT**g**ATGGAT**c**AAT | GCTTAGGGCATCAAGTCTGGGGTGTGATTTATTCCTGTTGATCTCAGACTGTTTACATGTGGAAAAAGTACAACAAGGTGTTTGGGAGTTCTGTATCTAAGTGTAGGGCTTGCTGGCTGGTGGCTTCCCTGT**GGGGTTATTGATCCATCACTAGGCATTT**CATTCTGGGATCCCCACATGGCAGAATCGGTGGATCTTTCCTCTTGGGGAAGTCCATGCCTTCAGGGAGAAATCTCCTAATTCCCTGCTTGGAGGGAGTGGGGATGTACAAGTCAACATCCTGGGAATCACTTAGCAGGAGAGACTTGGACC |
| Nsu-*Casp-3*-T1-N_4_CC-OFFT2 | chr1 | AACTGCC**att**TTATGGATGAA**a** | ATGAATACCTGTTTCCTGCTGTATATTTTATGTATATTACTAATAATTAGTAATCCTTACAACCCATAAAAATGAATTTTAATTAC**AACTGCCATTTTATGGATGAAACTGACC**TTCACAATCAGAATCAGAATTTTAAAATGAAAAACTTAAAAATTTAAAAAAATTAGAATTTAAACTTAGATTTTTTTTTTTGCTCCAATACCTTCACTTCTTCCATAATGCCAAGCTGACCTGATAAAATCAAGTCAGAAATAAACAGAGATTGAATTATCATGAAGGAACATGAGCATCGGAA |
| Nsu-*Casp-3*-T1-N_4_CC-OFFT3 | chr11 | AACTG**aa**TAGTTAT**a**GATGAA**a** | AGCCTCAATTCATGTGATGACTGCAAGTAATCTTACTGAAGACATTCTTTAGAATACAGTTTACAGAATGAACCTATTGAGTAAATATATTGTAAGA**AACTGAATAGTTATAGATGAAACTCTCC**ATCTTTATTTTTGTTATTTCACAATTACATGAATTTTTTTAAGTTAATCTATGAGTCAAAAAGTTCCAGAATTTCCAGGTATTTTTCTACTCTTCTCAGTTCACCAATAATAAAATATTCCAACTGGGGAATCATCACTAAAATATGCAAACCTTGATTTAATCAATGCCTGATATAATCTCCCTACTAAACTCCCATTTCCTAGCCTTTGCT |
| Nsu-*EMX*-T1-N_4_CC-OFFT1 | chr19 | AGC**c**AGGA**ga**CACAGCAGCTCT | CGTCCTCAGAGCCTTCCCAAATGCCCACACCGGGGGCACCCGGGTTTCCCCAAACACTCCAGGACAGAGCGGGGGCTTTGATCACAGGGAGGG**GGCTGCAGAGCTGCTGTGTCTCCTGGCT**CCTCTGCCCCATGCTCCCCACTCAGGGCTCCAGCAGTGGGCTCAGAGGACAGTGTGGCATTAGCATCCCCTGCCCGTCCAGGTTGTCACTCTCCTTGGCCCCTTGCTGCTCCCCAGTGGAAAAAAAAGGGGGAAATAAAGAATCGTAAAAAATACATATATTGGAACCTGTATGTTTTGTGATCTTCGCAACATTTCTCTGTAAACTCTCCTTCTTTAGGTTTTTGACAAAACCTGAATGCAGAAGTGGTCTCC |
| Nsu-*EMX*-T1-N_4_CC-OFFT2 | chr12 | AGCT**c**GGA**c**GC**g**CAGCAGCT**t**T | TAACCCTTAAGTGCTCCCGCTTCCTGCAGCCTCCACACACCACCAACTCTTCCAATCCTCCACTCTTCCATTCCTCCACTCTTCCAATCCTCGCTCTACAACCCCATCCTTACGATGTCATCATTCCTCCCTGGGTCGCGTTCCCACGGCAGCGCGCCTCTTCT**GGCTACAAAGCTGCTGCGCGTCCGAGCT**CTGCAGGGCGGAGACTCACCTACACAGCGGTATCCGAGGATGACCACCTTCCTGTAGCGGACTAGCGGCATGGCAGGAGCCCGCCCCAGGGGCTTGCGGAACTGCGGGCTCAGAGAGCCCGAAAACGAGGTCAGGGTGTGA |
| Nsu-*EMX*-T1-N_4_CC-OFFT3 | chr12 | **t**GCTAGGATGCA**ag**GCAGCTC**a** | GTGAGGGCTGAGGCTCTGACTTAGAAACCTTCCCAGTATGGTTAAAGAACAGCAAGGTCTACAGCAGGGGCAGTTGGGTATTAAGAGTGTCAGAGAGTTTATGACAGGAGAGTCTTGCAGATCAATATAGGGACTTTGTCTTTTTACTATGAATAAGTTAGGGAGTCCTTGGAGGATGTGGGCAAAGGA**GGCACATGAGCTGCCTTGCATCCTAGCA**GGATATCAGTCTGATGGCTGTGTTGAGGATATATGAAGAGGGCCAAGGGTAAAAGGCAGGGATACTAGTTAAGAAGCTGCTACAGAGGCAGGGGGAATAATTATGAAGCTGCTACAAAGAAACAATGGTGGTTTGGATCAGGCTGTTGGTAGTGAAGTCATGAG |

**Bold pink: mismatched nucleotides**

**Bold blue: off-target sequences**

**Bold blue: PAM sequences**

**Table S17. Off-target PCR primer sequences**

| **Primer Name** | **Sequence (5’---3’)** |
| --- | --- |
| Gfo-*AIFM*-T1-N_4_GAAA-OFFT1_Fwd | AGACAGTGTGATTCAGACAGTGG |
| Gfo-*AIFM*-T1-N_4_GAAA-OFFT1_Rev | CAGTTCACCACCAAGCAAGCC |
| Gfo-*AIFM*-T1-N_4_GAAA-OFFT2_Fwd | GAGGCCATCAGTTCCTTCCTGT |
| Gfo-*AIFM*-T1-N_4_GAAA-OFFT2_Rev | AAGGAAGGATGACAGGAGCTGG |
| Gfo-*AIFM*-T1-N_4_GAAA-OFFT3_Fwd | TCTCAGCACTGATCTCTCAGCC |
| Gfo-*AIFM*-T1-N_4_GAAA-OFFT3_Rev | TGACCTGTCTTGACCTCTCTGC |
| Gfo-*HBB*-T3-N_4_GAAA-OFFT1_Fwd | ACTCATGAACCACGCTCACCTT |
| Gfo-*HBB*-T3-N_4_GAAA-OFFT1_Rev | AAGAAGCAGGGCCCTAACTCAG |
| Gfo-*HBB*-T3-N_4_GAAA-OFFT2_Fwd | CCACGCCCATCTTAATTTGTAC |
| Gfo-*HBB*-T3-N_4_GAAA-OFFT2_Rev | GCAAATTACTAGATTCTTTACTAAGG |
| Gfo-*HBB*-T3-N_4_GAAA-OFFT3_Fwd | ATCCACTTAATTTGGGCAAGGT |
| Gfo-*HBB*-T3-N_4_GAAA-OFFT3_Rev | GCCTCCAGAAGCTGTTGCATAA |
| Nsu-*AIFM*-T1-N_4_CC-OFFT1_Fwd | GCTACACTCAGCAACAAGTGATAG |
| Nsu-*AIFM*-T1-N_4_CC-OFFT1_Rev | GGATGAAGCAGGACGGTATTCTTC |
| Nsu-*AIFM*-T1-N_4_CC-OFFT2_Fwd | AAGGTAAGCTTCCCCTGGAGTC |
| Nsu-*AIFM*-T1-N_4_CC-OFFT2_Rev | GTTGGAGAACCACAGTCCAAGTA |
| Nsu-*AIFM*-T1-N_4_CC-OFFT3_Fwd | AGGGTAGAGCATGATCTGGGCA |
| Nsu-*AIFM*-T1-N_4_CC-OFFT3_Rev | GCACACCAAAGTGACCAAAAGC |
| Nsu-*Casp-3*-T1-N_4_CC-OFFT1_Fwd | GCTTAGGGCATCAAGTCTGGGG |
| Nsu-*Casp-3*-T1-N_4_CC-OFFT1_Rev | GGTCCAAGTCTCTCCTGCTAAGT |
| Nsu-*Casp-3*-T1-N_4_CC-OFFT2_Fwd | ATGAATACCTGTTTCCTGCTGT |
| Nsu-*Casp-3*-T1-N_4_CC-OFFT2_Rev | TTCCGATGCTCATGTTCCTTCA |
| Nsu-*Casp-3*-T1-N_4_CC-OFFT3_Fwd | AGCCTCAATTCATGTGATGACTGC |
| Nsu-*Casp-3*-T1-N_4_CC-OFFT3_Rev | AGCAAAGGCTAGGAAATGGGAGT |
| Nsu-*EMX*-T1-N_4_CC-OFFT1_Fwd | CGTCCTCAGAGCCTTCCCAAA |
| Nsu-*EMX*-T1-N_4_CC-OFFT1_Rev | GGAGACCACTTCTGCATTCAGGT |
| Nsu-*EMX*-T1-N_4_CC-OFFT2_Fwd | TAACCCTTAAGTGCTCCCGCTT |
| Nsu-*EMX*-T1-N_4_CC-OFFT2_Rev | TCACACCCTGACCTCGTTTTCG |
| Nsu-*EMX*-T1-N_4_CC-OFFT3_Fwd | GTGAGGGCTGAGGCTCTGACT |
| Nsu-*EMX*-T1-N_4_CC-OFFT3_Rev | CTCATGACTTCACTACCAACAGCC |
| Psu-*AIFM*-T1-N_4_ATAA-OFFT1_Fwd | AGTTGTATTACCTGGGTTGGAGGT |
| Psu-*AIFM*-T1-N_4_ATAA-OFFT1_Rev | ATGTCAACCCTGTCCCTTCTCC |
| Psu-*AIFM*-T1-N_4_ATAA-OFFT2_Fwd | AGGCCAGATTCTGAGGCCTAAGAA |
| Psu-*AIFM*-T1-N_4_ATAA-OFFT2_Rev | TGTGTTGTAAGGAAAGGGCCACTCA |
| Psu-*AIFM*-T1-N_4_ATAA-OFFT3_Fwd | AGCATTTGTGAAATAGGGACTTTC |
| Psu-*AIFM*-T1-N_4_ATAA-OFFT3_Rev | CCTGGTGAGGAATGTCTTGCT |
| Psu-*AIFM*-T1-N_4_ACAA-OFFT1_Fwd | CACCAGGCCCATGTCATATGTT |
| Psu-*AIFM*-T1-N_4_ACAA-OFFT1_Rev | ACCAGGAAATCATGGCACACTTG |
| Psu-*AIFM*-T1-N_4_ACAA-OFFT2_Fwd | AACCATGAGCAGAGGAGGCAG |
| Psu-*AIFM*-T1-N_4_ACAA-OFFT2_Rev | GCCCTCATCTCTTGACCACAAC |
| Psu-*AIFM*-T1-N_4_ACAA-OFFT3_Fwd | CTGAGGAGCCAGGCCCTGAA |
| Psu-*AIFM*-T1-N_4_ACAA-OFFT3_Rev | CTTTGCTCAGTGCAGCCTCAACCTC |
| Psu-*CLTA*-T3-N_4_ACAA-OFFT1_Fwd | CCCTCCTCTGTCCCGTTTCTTC |
| Psu-*CLTA*-T3-N_4_ACAA-OFFT1_Rev | CACCATGGTTTAGAGAACCAAGACC |
| Psu-*CLTA*-T3-N_4_ACAA-OFFT2_Fwd | CCCTCCTCTGTCCCGTTTCTTC |
| Psu-*CLTA*-T3-N_4_ACAA-OFFT2_Rev | CACCATGGTTTAGAGAACCAAGACC |
| Psu-*CLTA*-T3-N_4_ACAA-OFFT3_Fwd | TCCCTGCCTCAGAGAAAAGACTG |
| Psu-*CLTA*-T3-N_4_ACAA-OFFT3_Rev | GAGCAACTCCTTGGGTGCAAAA |
| Psu-*EMX*-T1-N_4_ATAA-OFFT1_Fwd | ATGGATGTCTAGTTGCTCCAGCACC |
| Psu-*EMX*-T1-N_4_ATAA-OFFT1_Rev | AGAAGAAATGGACCAATTTCCTTG |
| Psu-*EMX*-T1-N_4_ATAA-OFFT2_Fwd | GATTGTTAGGTCTATTCTGGGAAC |
| Psu-*EMX*-T1-N_4_ATAA-OFFT2_Rev | TAAAATCTCCAGGCCCAGACC |
| Psu-*EMX*-T1-N_4_ATAA-OFFT3_Fwd | GCATGGGGAGCTGTGAAAGAAG |
| Psu-*EMX*-T1-N_4_ATAA-OFFT3_Rev | TCCACTGGGTGGATTGATTTGG |
| Psu-*EMX*-T1-N_4_ACAA-OFFT1_Fwd | AAGCTTTCAGGAACGGGCTTTG |
| Psu-*EMX*-T1-N_4_ACAA-OFFT1_Rev | AGCAGTCTTAGAGTCCTCCCGA |
| Psu-*EMX*-T1-N_4_ACAA-OFFT2_Fwd | AAAAAGAGTCAGTCCCTCTGGGC |
| Psu-*EMX*-T1-N_4_ACAA-OFFT2_Rev | GCAGCTGCAGAAAGACTCAGAT |
| Psu-*EMX*-T1-N_4_ACAA-OFFT3_Fwd | CCAGCTGGAGGGAAAAGGAAAG |
| Psu-*EMX*-T1-N_4_ACAA-OFFT3_Rev | CCATGTCCAGACCTGCTTCCTCT |
| Psu-*HBB*-T3-N_4_ATAA-OFFT1_Fwd | CAGCCCCAGGACCAAATTACCA |
| Psu-*HBB*-T3-N_4_ATAA-OFFT1_Rev | TACCTGCAGCACCTTGCTATCT |
| Psu-*HBB*-T3-N_4_ATAA-OFFT2_Fwd | GTGAAGTACAACTGCCCCTCCA |
| Psu-*HBB*-T3-N_4_ATAA-OFFT2_Rev | AGGTTGTGGAGGTTTACAGGTCT |
| Psu-*HBB*-T3-N_4_ATAA-OFFT3_Fwd | GCAGGAGGTAGTTTAAGGCCTACT |
| Psu-*HBB*-T3-N_4_ATAA-OFFT3_Rev | CAGGCAGTTGCCTCAGATTCTC |
| Illumina-Gfo-*AIFM*-T1-N_4_GAAA-OFFT1_Fwd | TCGTCGGCAGCGTCAGATGTGTATAAGAGACAGAGACAGTGTGATTCAGACAGTGG |
| Illumina-Gfo-*AIFM*-T1-N_4_GAAA-OFFT1_Rev | GTCTCGTGGGCTCGGAGATGTGTATAAGAGACAGCAGTTCACCACCAAGCAAGCC |
| Illumina-Gfo-*AIFM*-T1-N_4_GAAA-OFFT2_Fwd | TCGTCGGCAGCGTCAGATGTGTATAAGAGACAGGAGGCCATCAGTTCCTTCCTGT |
| Illumina-Gfo-*AIFM*-T1-N_4_GAAA-OFFT2_Rev | GTCTCGTGGGCTCGGAGATGTGTATAAGAGACAGAAGGAAGGATGACAGGAGCTGG |
| Illumina-Gfo-*AIFM*-T1-N_4_GAAA-OFFT3_Fwd | TCGTCGGCAGCGTCAGATGTGTATAAGAGACAGTCTCAGCACTGATCTCTCAGCC |
| Illumina-Gfo-*AIFM*-T1-N_4_GAAA-OFFT3_Rev | GTCTCGTGGGCTCGGAGATGTGTATAAGAGACAGTGACCTGTCTTGACCTCTCTGC |
| Illumina-Gfo-*HBB*-T3-N_4_GAAA-OFFT1_Fwd | TCGTCGGCAGCGTCAGATGTGTATAAGAGACAGACTCATGAACCACGCTCACCTT |
| Illumina-Gfo-*HBB*-T3-N_4_GAAA-OFFT1_Rev | GTCTCGTGGGCTCGGAGATGTGTATAAGAGACAGAAGAAGCAGGGCCCTAACTCAG |
| Illumina-Gfo-*HBB*-T3-N_4_GAAA-OFFT2_Fwd | TCGTCGGCAGCGTCAGATGTGTATAAGAGACAGCCACGCCCATCTTAATTTGTAC |
| Illumina-Gfo-*HBB*-T3-N_4_GAAA-OFFT2_Rev | GTCTCGTGGGCTCGGAGATGTGTATAAGAGACAGGCAAATTACTAGATTCTTTACTAAGG |
| Illumina-Gfo-*HBB*-T3-N_4_GAAA-OFFT3_Fwd | TCGTCGGCAGCGTCAGATGTGTATAAGAGACAGATCCACTTAATTTGGGCAAGGT |
| Illumina-Gfo-*HBB*-T3-N_4_GAAA-OFFT3_Rev | GTCTCGTGGGCTCGGAGATGTGTATAAGAGACAGGCCTCCAGAAGCTGTTGCATAA |
| Illumina-Nsu-*AIFM*-T1-N_4_CC-OFFT1_Fwd | TCGTCGGCAGCGTCAGATGTGTATAAGAGACAGGCTACACTCAGCAACAAGTGATAG |
| Illumina-Nsu-*AIFM*-T1-N_4_CC-OFFT1_Rev | GTCTCGTGGGCTCGGAGATGTGTATAAGAGACAGGGATGAAGCAGGACGGTATTCTTC |
| Illumina-Nsu-*AIFM*-T1-N_4_CC-OFFT2_Fwd | TCGTCGGCAGCGTCAGATGTGTATAAGAGACAGAAGGTAAGCTTCCCCTGGAGTC |
| Illumina-Nsu-*AIFM*-T1-N_4_CC-OFFT2_Rev | GTCTCGTGGGCTCGGAGATGTGTATAAGAGACAGGTTGGAGAACCACAGTCCAAGTA |
| Illumina-Nsu-*AIFM*-T1-N_4_CC-OFFT3_Fwd | TCGTCGGCAGCGTCAGATGTGTATAAGAGACAGAGGGTAGAGCATGATCTGGGCA |
| Illumina-Nsu-*AIFM*-T1-N_4_CC-OFFT3_Rev | GTCTCGTGGGCTCGGAGATGTGTATAAGAGACAGGCACACCAAAGTGACCAAAAGC |
| Illumina-Nsu-*Casp-3*-T1-N_4_CC-OFFT1_Fwd | TCGTCGGCAGCGTCAGATGTGTATAAGAGACAGGCTTAGGGCATCAAGTCTGGGG |
| Illumina-Nsu-*Casp-3*-T1-N_4_CC-OFFT1_Rev | GTCTCGTGGGCTCGGAGATGTGTATAAGAGACAGGGTCCAAGTCTCTCCTGCTAAGT |
| Illumina-Nsu-*Casp-3*-T1-N_4_CC-OFFT2_Fwd | TCGTCGGCAGCGTCAGATGTGTATAAGAGACAGATGAATACCTGTTTCCTGCTGT |
| Illumina-Nsu-*Casp-3*-T1-N_4_CC-OFFT2_Rev | GTCTCGTGGGCTCGGAGATGTGTATAAGAGACAGTTCCGATGCTCATGTTCCTTCA |
| Illumina-Nsu-*Casp-3*-T1-N_4_CC-OFFT3_Fwd | TCGTCGGCAGCGTCAGATGTGTATAAGAGACAGAGCCTCAATTCATGTGATGACTGC |
| Illumina-Nsu-*Casp-3*-T1-N_4_CC-OFFT3_Rev | GTCTCGTGGGCTCGGAGATGTGTATAAGAGACAGAGCAAAGGCTAGGAAATGGGAGT |
| Illumina-Nsu-*EMX*-T1-N_4_CC-OFFT1_Fwd | TCGTCGGCAGCGTCAGATGTGTATAAGAGACAGCGTCCTCAGAGCCTTCCCAAA |
| Illumina-Nsu-*EMX*-T1-N_4_CC-OFFT1_Rev | GTCTCGTGGGCTCGGAGATGTGTATAAGAGACAGGGAGACCACTTCTGCATTCAGGT |
| Illumina-Nsu-*EMX*-T1-N_4_CC-OFFT2_Fwd | TCGTCGGCAGCGTCAGATGTGTATAAGAGACAGTAACCCTTAAGTGCTCCCGCTT |
| Illumina-Nsu-*EMX*-T1-N_4_CC-OFFT2_Rev | GTCTCGTGGGCTCGGAGATGTGTATAAGAGACAGTCACACCCTGACCTCGTTTTCG |
| Illumina-Nsu-*EMX*-T1-N_4_CC-OFFT3_Fwd | TCGTCGGCAGCGTCAGATGTGTATAAGAGACAGGTGAGGGCTGAGGCTCTGACT |
| Illumina-Nsu-*EMX*-T1-N_4_CC-OFFT3_Rev | GTCTCGTGGGCTCGGAGATGTGTATAAGAGACAGCTCATGACTTCACTACCAACAGCC |
| Illumina-Psu-*AIFM*-T1-N_4_ATAA-OFFT1_Fwd | TCGTCGGCAGCGTCAGATGTGTATAAGAGACAGAGTTGTATTACCTGGGTTGGAGGT |
| Illumina-Psu-*AIFM*-T1-N_4_ATAA-OFFT1_Rev | GTCTCGTGGGCTCGGAGATGTGTATAAGAGACAGATGTCAACCCTGTCCCTTCTCC |
| Illumina-Psu-*AIFM*-T1-N_4_ATAA-OFFT2_Fwd | TCGTCGGCAGCGTCAGATGTGTATAAGAGACAGAGGCCAGATTCTGAGGCCTAAGAA |
| Illumina-Psu-*AIFM*-T1-N_4_ATAA-OFFT2_Rev | GTCTCGTGGGCTCGGAGATGTGTATAAGAGACAGTGTGTTGTAAGGAAAGGGCCACTCA |
| Illumina-Psu-*AIFM*-T1-N_4_ATAA-OFFT3_Fwd | TCGTCGGCAGCGTCAGATGTGTATAAGAGACAGAGCATTTGTGAAATAGGGACTTTC |
| Illumina-Psu-*AIFM*-T1-N_4_ATAA-OFFT3_Rev | GTCTCGTGGGCTCGGAGATGTGTATAAGAGACAGCCTGGTGAGGAATGTCTTGCT |
| Illumina-Psu-*AIFM*-T1-N_4_ACAA-OFFT1_Fwd | TCGTCGGCAGCGTCAGATGTGTATAAGAGACAGCACCAGGCCCATGTCATATGTT |
| Illumina-Psu-*AIFM*-T1-N_4_ACAA-OFFT1_Rev | GTCTCGTGGGCTCGGAGATGTGTATAAGAGACAGACCAGGAAATCATGGCACACTTG |
| Illumina-Psu-*AIFM*-T1-N_4_ACAA-OFFT2_Fwd | TCGTCGGCAGCGTCAGATGTGTATAAGAGACAGAACCATGAGCAGAGGAGGCAG |
| Illumina-Psu-*AIFM*-T1-N_4_ACAA-OFFT2_Rev | GTCTCGTGGGCTCGGAGATGTGTATAAGAGACAGGCCCTCATCTCTTGACCACAAC |
| Illumina-Psu-*AIFM*-T1-N_4_ACAA-OFFT3_Fwd | TCGTCGGCAGCGTCAGATGTGTATAAGAGACAGCTGAGGAGCCAGGCCCTGAA |
| Illumina-Psu-*AIFM*-T1-N_4_ACAA-OFFT3_Rev | GTCTCGTGGGCTCGGAGATGTGTATAAGAGACAGCTTTGCTCAGTGCAGCCTCAACCTC |
| Illumina-Psu-*CLTA*-T3-N_4_ACAA-OFFT1_Fwd | TCGTCGGCAGCGTCAGATGTGTATAAGAGACAGCCCTCCTCTGTCCCGTTTCTTC |
| Illumina-Psu-*CLTA*-T3-N_4_ACAA-OFFT1_Rev | GTCTCGTGGGCTCGGAGATGTGTATAAGAGACAGCACCATGGTTTAGAGAACCAAGACC |
| Illumina-Psu-*CLTA*-T3-N_4_ACAA-OFFT2_Fwd | TCGTCGGCAGCGTCAGATGTGTATAAGAGACAGCCCTCCTCTGTCCCGTTTCTTC |
| Illumina-Psu-*CLTA*-T3-N_4_ACAA-OFFT2_Rev | GTCTCGTGGGCTCGGAGATGTGTATAAGAGACAGCACCATGGTTTAGAGAACCAAGACC |
| Illumina-Psu-*CLTA*-T3-N_4_ACAA-OFFT3_Fwd | TCGTCGGCAGCGTCAGATGTGTATAAGAGACAGTCCCTGCCTCAGAGAAAAGACTG |
| Illumina-Psu-*CLTA*-T3-N_4_ACAA-OFFT3_Rev | GTCTCGTGGGCTCGGAGATGTGTATAAGAGACAGGAGCAACTCCTTGGGTGCAAAA |
| Illumina-Psu-*EMX*-T1-N_4_ATAA-OFFT1_Fwd | TCGTCGGCAGCGTCAGATGTGTATAAGAGACAGATGGATGTCTAGTTGCTCCAGCACC |
| Illumina-Psu-*EMX*-T1-N_4_ATAA-OFFT1_Rev | GTCTCGTGGGCTCGGAGATGTGTATAAGAGACAGAGAAGAAATGGACCAATTTCCTTG |
| Illumina-Psu-*EMX*-T1-N_4_ATAA-OFFT2_Fwd | TCGTCGGCAGCGTCAGATGTGTATAAGAGACAGGATTGTTAGGTCTATTCTGGGAAC |
| Illumina-Psu-*EMX*-T1-N_4_ATAA-OFFT2_Rev | GTCTCGTGGGCTCGGAGATGTGTATAAGAGACAGTAAAATCTCCAGGCCCAGACC |
| Illumina-Psu-*EMX*-T1-N_4_ATAA-OFFT3_Fwd | TCGTCGGCAGCGTCAGATGTGTATAAGAGACAGGCATGGGGAGCTGTGAAAGAAG |
| Illumina-Psu-*EMX*-T1-N_4_ATAA-OFFT3_Rev | GTCTCGTGGGCTCGGAGATGTGTATAAGAGACAGTCCACTGGGTGGATTGATTTGG |
| Illumina-Psu-*EMX*-T1-N_4_ACAA-OFFT1_Fwd | TCGTCGGCAGCGTCAGATGTGTATAAGAGACAGAAGCTTTCAGGAACGGGCTTTG |
| Illumina-Psu-*EMX*-T1-N_4_ACAA-OFFT1_Rev | GTCTCGTGGGCTCGGAGATGTGTATAAGAGACAGAGCAGTCTTAGAGTCCTCCCGA |
| Illumina-Psu-*EMX*-T1-N_4_ACAA-OFFT2_Fwd | TCGTCGGCAGCGTCAGATGTGTATAAGAGACAGAAAAAGAGTCAGTCCCTCTGGGC |
| Illumina-Psu-*EMX*-T1-N_4_ACAA-OFFT2_Rev | GTCTCGTGGGCTCGGAGATGTGTATAAGAGACAGGCAGCTGCAGAAAGACTCAGAT |
| Illumina-Psu-*EMX*-T1-N_4_ACAA-OFFT3_Fwd | TCGTCGGCAGCGTCAGATGTGTATAAGAGACAGCCAGCTGGAGGGAAAAGGAAAG |
| Illumina-Psu-*EMX*-T1-N_4_ACAA-OFFT3_Rev | GTCTCGTGGGCTCGGAGATGTGTATAAGAGACAGCCATGTCCAGACCTGCTTCCTCT |
| Illumina-Psu-*HBB*-T3-N_4_ATAA-OFFT1_Fwd | TCGTCGGCAGCGTCAGATGTGTATAAGAGACAGCAGCCCCAGGACCAAATTACCA |
| Illumina-Psu-*HBB*-T3-N_4_ATAA-OFFT1_Rev | GTCTCGTGGGCTCGGAGATGTGTATAAGAGACAGTACCTGCAGCACCTTGCTATCT |
| Illumina-Psu-*HBB*-T3-N_4_ATAA-OFFT2_Fwd | TCGTCGGCAGCGTCAGATGTGTATAAGAGACAGGTGAAGTACAACTGCCCCTCCA |
| Illumina-Psu-*HBB*-T3-N_4_ATAA-OFFT2_Rev | GTCTCGTGGGCTCGGAGATGTGTATAAGAGACAGAGGTTGTGGAGGTTTACAGGTCT |
| Illumina-Psu-*HBB*-T3-N_4_ATAA-OFFT3_Fwd | TCGTCGGCAGCGTCAGATGTGTATAAGAGACAGGCAGGAGGTAGTTTAAGGCCTACT |
| Illumina-Psu-*HBB*-T3-N_4_ATAA-OFFT3_Rev | GTCTCGTGGGCTCGGAGATGTGTATAAGAGACAGCAGGCAGTTGCCTCAGATTCTC |

**Dataset S1 (separate file):** Detailed information of the sources of NsuCas9, PsuCas9 and GfoCas9; sequences of reference proteins used to conduct the phylogenetic analysis.
